# A *trans*-translation inhibitor kills *Mycobacterium tuberculosis* by targeting ribosomal protein bL12

**DOI:** 10.1101/2025.06.02.656638

**Authors:** Akanksha Varshney, John N. Alumasa, Amber Miller, Kenneth C. Keiler

## Abstract

New antibiotics with novel mechanisms of action are needed to treat infections by multidrug-resistant strains of *Mycobacterium tuberculosis*. Here, we show that KKL-1005, an anti-tubercular triazole-based molecule, binds to ribosomal protein bL12 and specifically inhibits the *trans*-translation ribosome rescue pathway, a process essential for the survival of *M. tuberculosis.* Our data demonstrate that KKL-1005 binds to the N terminal domain of bL12, both *in vitro* and in bacterial cells, and specifically inhibits *trans*-translation and not normal translation. These results suggest that tmRNA-SmpB interacts with bL12 differently from tRNA, and raise the possibility of developing antibiotics targeting bL12.

## INTRODUCTION

Infections caused by *M. tuberculosis* kill over 1.5 million annually (WHO 2024). Despite the availability of anti-tubercular drugs, the emergence and rapid spread of multidrug-resistant strains threaten public health (Nimmo et al. 2022; Serajian et al. 2025). To address this growing threat, the discovery of novel anti-tubercular agents that act on previously unexploited molecular targets is essential. Beyond their clinical potential, antibiotics with new mechanisms of action are valuable as chemical probes to dissect fundamental aspects of bacterial physiology.

The *trans*-translation pathway for ribosome rescue is a potential antibiotic target because it is essential in many human pathogens, including *M. tuberculosis*, and is not present in humans (Personne and Parish 2014; Alumasa et al. 2017). *trans*-Translation rescues ribosomes that are stalled at the 3’ end of mRNA lacking a stop codon (Keiler et al. 1996; Keiler 2015). Because the frequency of these “non-stop” ribosomes is high, bacteria cannot survive unless non-stop ribosomes are rescued (Ito et al. 2011; Keiler 2015). The key molecules for *trans*-translation are a specialized RNA known as transfer-messenger RNA (tmRNA), and a small protein, SmpB (Keiler et al. 1996; Karzai et al. 1999). During *trans*-translation, the tRNA-like domain of tmRNA is charged with alanine by alanyl-tRNA synthase and EF-Tu delivers the tmRNA-SmpB complex to the A site of the non-stop ribosome (Barends et al. 2000). The nascent polypeptide is transpeptidated onto tmRNA and tmRNA-SmpB is translocated into the P site of the ribosome (Ramrath et al. 2012). During translocation, a specialized reading frame within tmRNA is inserted into the mRNA channel. Translation resumes using tmRNA as the template and terminates at the stop codon at the end of the tmRNA reading frame, releasing the ribosome and the nascent polypeptide, which now contains the tmRNA-encoded tag peptide (Keiler et al. 1996; Keiler 2015). Because the tag peptide is recognized by several proteases, the tagged protein is rapidly degraded (Keiler et al. 1996; Gottesman et al. 1998; Herman et al. 1998; Flynn et al. 2001).

We previously found that some small molecule inhibitors of *trans*-translation have potent antibiotic properties, including against *M. tuberculosis* (Ramadoss et al. 2013; Alumasa et al. 2017; Varshney et al. 2025). To search for additional *trans*-translation inhibitors with anti-mycobacterial activity, we screened a compound library from TB Alliance comprising molecules that inhibited the growth of *M. tuberculosis* in culture but did not have a known mechanism of action. Here, we show that one hit from this library, KKL-1005, inhibits *trans-*translation by binding ribosomal protein bL12. Analysis of KKL-1005 activity suggests a that tmRNA-SmpB interacts with the ribosome in a fundamentally different way from tRNAs and these differences provide opportunities for antibiotic development.

## RESULTS

### A luciferase-based screen identified new inhibitors of *trans*-translation

We screened a library of 1,608 antitubercular small molecules for their ability to inhibit *trans*-translation using a luciferase reporter in *E. coli*. In this *luc-trpAt* reporter, the firefly luciferase gene has a trpA transcriptional terminator before the stop codon, such that luciferase is translated from a non-stop mRNA (Ramadoss et al. 2013). When *trans*-translation is active the luciferase will be tagged and degraded, but when *trans*-translation is inhibited active luciferase will accumulate, resulting in a luminescent signal when luciferin is added. Luciferase activity was measured after growth in the presence of compounds from the library using the known *trans*-translation inhibitor KKL-35 as a positive control (Fig. 1A, Table S1) (Ramadoss et al. 2013; Varshney et al. 2025) and those 2 standard deviations above the average were scored as positive. Among the positive compounds, the triazole furan carboxamide compound KKL-1005 (*N*-(5-benzamido-2-phenyl-1,2,4-triazol-3-yl)furan-2-carboxamide) was chosen for further investigation here. To confirm that the observed activity of KKL-1005 in *E. coli* is due to inhibition of *trans*-translation and not an effect on luciferase activity, we repeated the *trans*-translation inhibition assays using an mCherry-trpAt reporter that is analogous to the luc-trpAt reporter in that it produces mCherry from a non-stop mRNA (Ramadoss et al. 2013). Incubation of *E. coli* expressing mCherry-trpAt with KKL-1005 resulted in a dose-dependent increase in fluorescence, consistent with inhibition of *trans*-translation (Fig. 1B). Fluorescence did not reach the level seen when *trans*-translation activity was eliminated by deleting *ssrA*, the gene encoding tmRNA, but the IC_50_ for inhibition by KKL-1005 was <1.5 µM, indicating potent activity (Fig. 1B). KKL-1005-treated cells grew to the same level as the DMSO-treated control, indicating that KKL-1005 did not inhibit the growth of the cells expressing mCherry at the concentrations used in the experiment (Fig. S1). These results demonstrate that KKL-1005 inhibits *trans*-translation in *E. coli* cells.

**Fig. 1:**
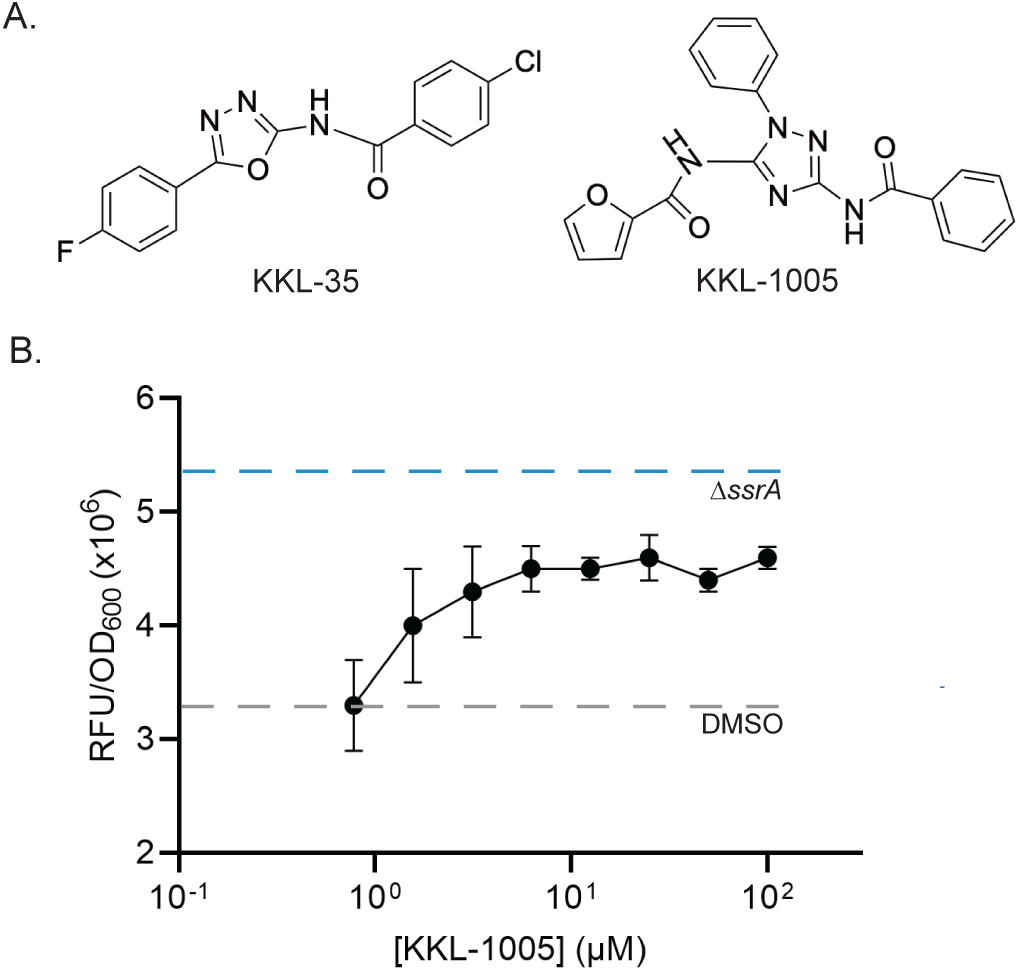
KKL-1005 inhibits *trans*-translation in *E. coli.* A) Chemical structures of a previously identified *trans*-translation inhibitor, KKL-35 and KKL-1005. B) Inhibition of *trans*-translation in *E. coli* was monitored using the *mCherry-*trpAt reporter. Wild-type *E. coli* was treated with KKL-1005 (circles) or DMSO and *E. coli ΔssrA* was treated with DMSO and the mCherry signal per OD600 was measured. Data are the mean with the error bars representing the standard deviation for three biological replicates.

### KKL-1005 has anti-tubercular activity

KKL-1005 has previously been identified by Southern Research Institute in a high-throughput screen for inhibitors of *M. tuberculosis* growth (PubChem AID 1626). In this screen KKL-1005 inhibited growth of strain H37Rv by 95.5% when present at 10 µM (3.7 µg/ml). To assess whether our synthesis of KKL-1005 also had anti-tubercular activity we performed broth dilution and plating assays (Table 1). Consistent with the published data PubChem AID 1626, KKL-1005 inhibited growth of the attenuated *M. tuberculosis* strain H37Rv Δ*RD1 ΔpanCD* with a MIC = 4.7 µg/ml. Plating assays showed that KKL-1005 is bactericidal against this strain with a MBC = 4.7 µg/mL. Assays against other bacterial species indicated preferential antibiotic activity against *M. tuberculosis* (Table 1). However, KKL-1005 was reported to be cytotoxic against Vero cells with a CC_50_ = 3.3 µg/mL (8.8 µM) (PubChem AID 1626). We measured the cytotoxicity of KKL-1005 against HeLa cells and found a CC_50_ = 7.5 µg/mL (20 ± 2 µM) (Fig. S2). Because there is no *trans*-translation in eukaryotes, the cytotoxicity must me due to off-target effects. Nevertheless, the cytotoxicity would have to be greatly reduced to develop molecules like KKL-1005 as antibiotics.

**Table 1:** KKL-1005 inhibits bacterial growth.

| Compound | *M. tuberculosis*<br>H37Rv $\Delta RD1$<br>$\Delta panCD$ | | *M. avium* | | *B. anthracis* | | *E. coli* $\Delta tolC$ | |
| --- | --- | --- | --- | --- | --- | --- | --- | --- |
|  | MIC<sup>a</sup> | MBC<sup>b</sup> | MIC | MBC | MIC | MBC | MIC | MBC |
| KKL-1005 | 4.7<br>(12.5) | 4.7<br>(12.5) | 18.6<br>(50) | 18.6<br>(50) | 9.3<br>(25) | ND<sup>c</sup> | 9.3<br>(25) | 9.3<br>(25) |
| Rifampicin | 0.12<br>(0.14) | 0.12<br>(0.14) | 0.5<br>(0.6) | 1<br>(1.2) | ND | ND | ND | ND |

<sup>a</sup> MIC $\mu\text{g/mL}$ ( $\mu\text{M}$ ) values from at least three broth microdilution assays.
<sup>b</sup> MBC; $\mu\text{g/mL}$ ( $\mu\text{M}$ ) values from at least three plating assays.
<sup>c</sup> ND: Not determined.

### KKL-1005 inhibits *M. tuberculosis trans*-translation in *in vitro*

To determine if KKL-1005 inhibits *trans*-translation on *M. tuberculosis* ribosomes, we used an *in vitro trans*-translation assay that comprised purified *M. tuberculosis* ribosomes, tmRNA-SmpB, and translation factors, combined with purified *E. coli* tRNAs, tRNA aminoacyl-tRNA synthetases, methionyl-tRNA formyltransferase, and nucleoside diphosphate kinase, and purchased T7 RNA polymerase, nucleoside triphosphates, amino acids, and salts (Varshney et al. 2025). The *trans*-translation reaction was programmed with a PCR product encoding the dihydrofolate reductase (DHFR) protein that contained a T7 RNA polymerase promoter and no in-frame stop codon *(dhfr-ns*).

Synthesized DHFR protein was detected by incorporation of [^35^S]-methionine followed by SDS-PAGE and phosphorimaging. As previously observed with this assay, *in vitro trans*-translation resulted in both full-length DHFR protein (DHFR-ns), and DHFR with the tmRNA-encoded peptide tag (Varshney et al. 2025) (Fig. 2A). Including KKL-1005 in the translation reaction decreased the fraction of the protein that was tagged in a dose-dependent manner, with an IC_50_ = 37 ± 2 μM (Fig. 2A & B). When a stop codon was added to the gene encoding DHFR (*dhfr-stop*) and tmRNA-SmpB was omitted from the reaction, KKL-1005 did not inhibit translation of DHFR-stop (Fig. 2C). Conversely, adding chloramphenicol in these reactions completely inhibited the synthesis of DHFR-stop (Fig. 2C). *In vitro trans*-translation assays performed with purified *E. coli* components show that KKL-1005 also inhibits *trans*-translation on *E. coli* ribosomes (Fig. S3). These results demonstrate that KKL-1005 specifically inhibits *trans*-translation and not normal translation.

**Fig. 2:**
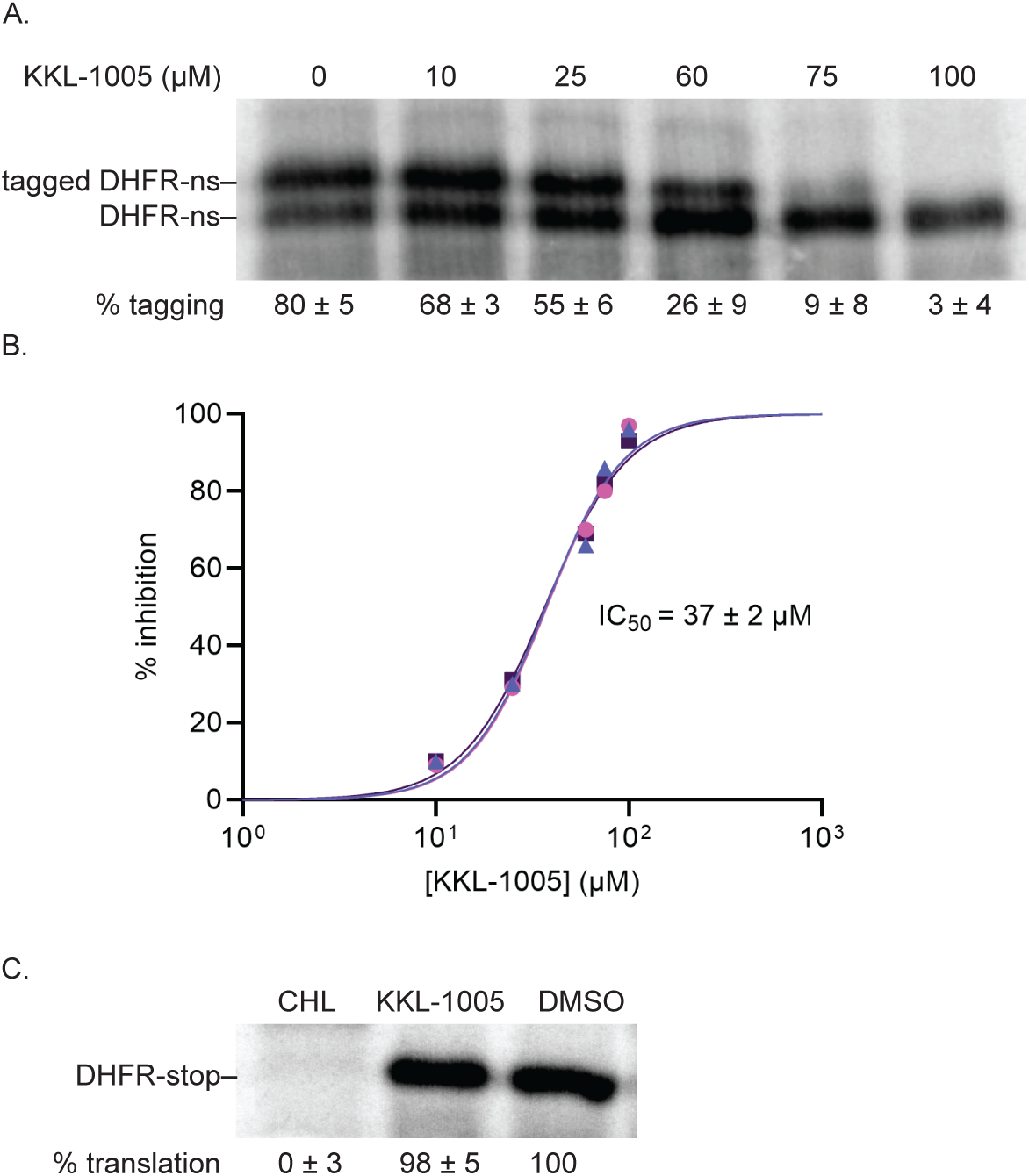
KKL-1005 inhibits *M. tuberculosis trans*-translation *in vitro.* A) A gene encoding DHFR without a stop codon was expressed in the presence of *M. tuberculosis* tmRNA-SmpB and varying concentrations of KKL-1005. Synthesized protein was detected by incorporation of ^35^S-methionine, followed by SDS-PAGE and phosphorimaging. Bands corresponding to tagged and untagged DHFR are indicated, and the average percentage of DHFR protein found in the tagged band for three repeats is shown with the standard deviation. B) Data from gels as in (A) were plotted and fit with a sigmoidal function to determine the IC50. C) *In vitro* translation was assayed from the expression of a gene encoding DHFR with a stop codon in the presence of DMSO, 100 μM chloramphenicol (CHL), or 100 μM KKL-1005, and a representative experiment is shown. The percentage of DHFR with respect to the amount in the DMSO-treated control is shown as the average from two independent repeats with the standard deviation.

### KKL-1005 binds the N-terminal domain of ribosomal protein bL12

To identify the molecular target of KKL-1005, we synthesized KKL-2108, an analog of KKL-1005 containing an azide functional group for photoactivatable crosslinking and an alkyne moiety for subsequent click bio-conjugation to assist in target identification (Fig. 3A) (Alumasa and Keiler 2015). To ensure that KKL-2108 retains the biochemical activity of KKL-1005 and would therefore be expected to bind the same target, we measured the growth inhibition of *E.coli ΔtolC* after treatment with KKL-2108 and KKL-1005. The dose-response curves for growth inhibition of *E. coli ΔtolC* for KKL-2108 and KKL-1005 were similar, suggesting that the structural changes did not disrupt the antibacterial properties (Fig. S4). Likewise, the MIC values for KKL-2108 against *E. coli ΔtolC* and *B. anthracis* were similar to KKL-1005 (Table S2). To facilitate isolation of the molecular target, we also synthesized a trifunctional probe, KKL-2107, that incorporated an azide group, a biotin group to allow affinity purification, and a fluorescent dansyl group for visualization (Alumasa et al. 2017) (Fig. 3A). To identify molecules that bind to KKL-2108 we incubated *B. anthracis* cells with KKL-2108 and initiated cross-linking by irradiating the cells with UV light (Fig. 3B). Cells were lysed and KKL-2107 was covalently linked to KKL-2108 using a copper-catalyzed reaction. StrepTactin resin was used to pull down KKL-2107 with the covalently linked KKL-2108 and any molecules that cross-linked to KKL-2108 inside the cell (Fig. 3B). Analysis of the affinity-purified proteins using SDS-PAGE revealed a single fluorescent band in the eluate (Fig. 3C & D). Mass spectrometry analysis of this band identified it as ribosomal protein bL12 (Table S3). The *in vivo* labeling was repeated with *E. coli ΔtolC*, and ribosomal protein bL12 was again identified as the cross-linked species (Fig. S5).

**Fig. 3:**
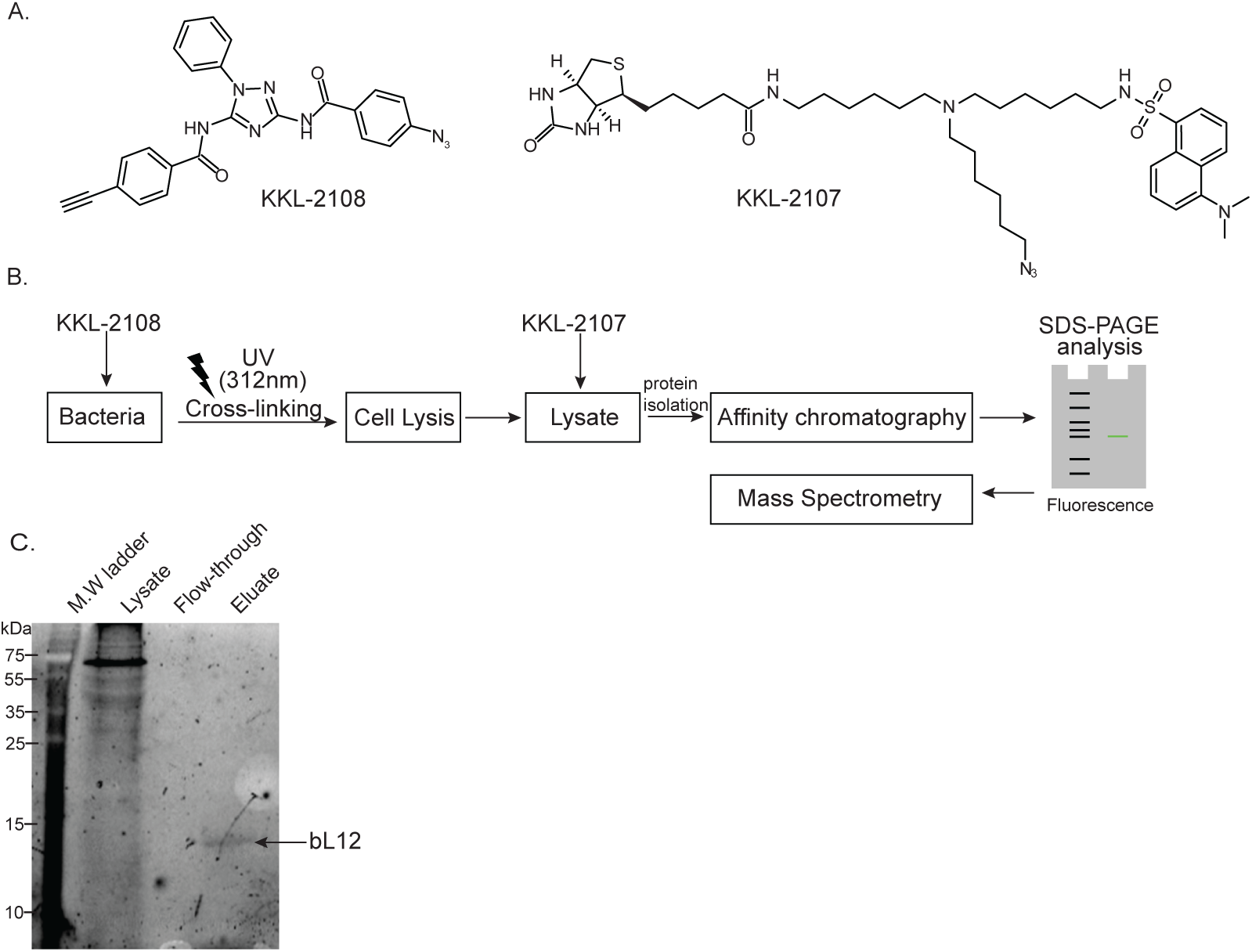
KKL-1005 binds *E. coli* bL12 *in vivo.* A) Chemical structures of cross-linkable KKL-1005 analog, KKL-2108, and a tri-functional reagent, KKL-2107, containing biotin and azide functional groups and a fluorescent dansyl moiety. B) Target identification workflow. The photolabile probe KKL-2108 was added to a growing *B. anthracis* culture. Cells were irradiated with UV light to activate the probe and enable cross-linking. Cells were lysed, and the lysate was subjected to click chemistry with KKL-2107 and analyzed by SDS-PAGE. C) Coomassie stained gel analysis of the affinity chromatography shown in (B). D) Fluorescence scan of a gel analysis of the samples from (C) with controls for the lysate after the click reaction. The band identified by mass spectrometry is indicated.

We measured the binding of KKL-1005 with purified *E. coli* bL12 *in vitro* using microscale thermophoresis (MST) and observed dose-dependent binding with equilibrium binding constant (K_d_) = 8 ± 2 µM (Fig. 4A). KKL-1005 also binds with purified *M. tuberculosis* bL12 with K_d_ = 4 ± 0.2 µM (Fig. S6). Structural studies have shown that bL12 consists of globular N-terminal and C-terminal domains connected by a flexible linker (Liljas and Gudkov 1987; Bocharov et al. 2004; Mulder et al. 2004). The NTD is responsible for dimerization of bL12 and anchors bL12 to the ribosome through interaction with bL10 (Wahl et al. 2000; Diaconu et al. 2005). The CTD interacts with the translation factors, IF2, EF-Tu, EF-G, and RF-3, and plays a key role in stimulating their GTPase activities (Savelsbergh et al. 2000; Wahl and Moller 2002; Bocharov et al. 2004; Helgstrand et al. 2007; Huang et al. 2010; Ge et al. 2018). To determine which domain binds with KKL-1005, we repeated the MST experiments using purified *E. coli* bL12 NTD or CTD. KKL-1005 binds with the NTD with K_d_ = 11 ± 3 µM, similar to the full length bL12 (Fig. 4B). Conversely, little interaction was observed between KKL-1005 and the CTD (Fig. 4C). Binding did not reach saturation within the solubility range of KKL-1005 so an accurate K_d_ could not be calculated, but the value was >1000 µM (Fig. 4C). These data indicate that KKL-1005 binds the NTD of bL12 and demonstrate that KKL-1005 can bind free bL12 in the absence of the ribosome.

**Fig. 4:**
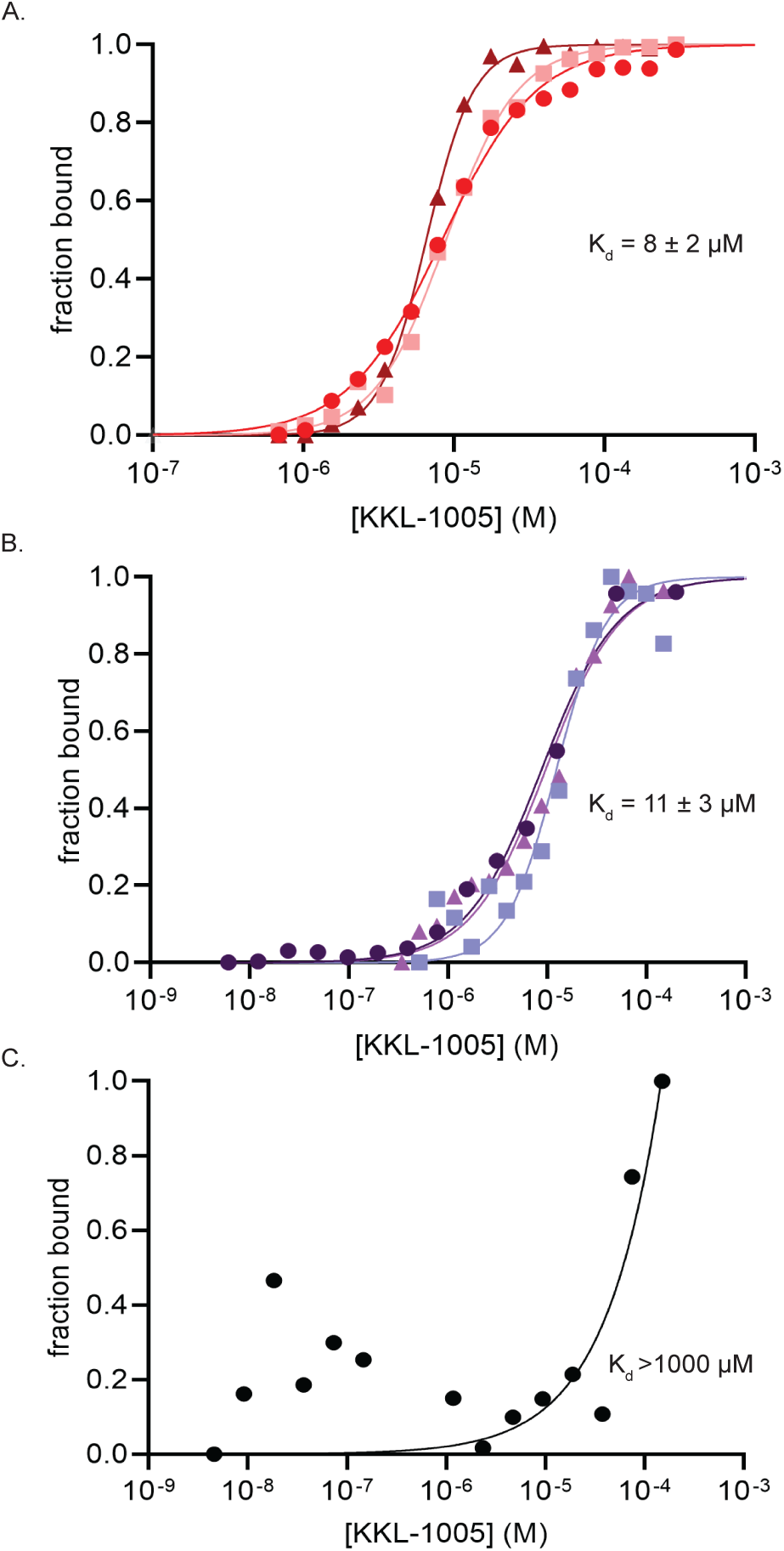
KKL-1005 binds to the N-terminal domain of bL12. A) MST was used to measure the binding of *E. coli* bL12 and KKL-1005 *in vitro*. Change in fluorescence was measured for fluorescently labeled bL12 with different concentrations of KKL-1005, and the fraction of bL12 bound with KKL-1005 was plotted. The dissociation constant (Kd) was calculated by non-linear curve fitting, and the Kd with standard deviations for at least three repeats is shown. B) MST as in (A) to measure the binding of bL12 NTD with KKL-1005. C) MST as in (A) to measure the binding of bL12 CTD with KKL-1005.

### Over-expression of bL12 suppresses growth inhibition by KKL-1005

To determine whether bL12 is the biological target responsible for growth inhibition by KKL-1005, we investigated the effects of over-expressing bL12 on the antibacterial activity of KKL-1005. We over-expressed bL12 in *E. coli ΔtolC* cells that had been transformed with a plasmid containing the *rplL* gene under the control of an IPTG-inducible promoter (Fig. S7A). We first demonstrated that over-expression of bL12 did not alter the growth kinetics of *E. coli* (Fig. S7B). To assess the effect of over-expressing bL12 on the antibacterial activity of KKL-1005, we cultured the bL12-expressing *E. coli ΔtolC* strain with and without the addition of IPTG and in the presence of varying concentrations of KKL-1005 (0 – 15 µg/ml). Optical density measurements of the uninduced strain in the presence of KKL-1005 showed significant inhibition and impaired growth rates at KKL-1005 concentrations ≥ 8 µg/ml (0.8X MIC), (Fig. 5A-F). Induction of bL12 increased growth rates in the presence of intermediate (8 – 11 µg/ml) concentrations of KKL-1005 (Fig. 5C-D & F). Higher concentrations of KKL-1005 inhibited growth even when bL12 was over-expressed (Fig. 5E & F). Over-expression of bL12 did not alter growth inhibition by KKL-35, which has an unrelated molecular structure and binds the 23s rRNA near the peptidyl transfer center (Fig. 5G–L). These results indicate that over-expression of bL12 specifically impairs the activity of KKL-1005 but does not rescue growth inhibition by other *trans*-translation inhibitors, consistent with bL12 acting as the molecular target for KKL-1005.

**Fig. 5:**
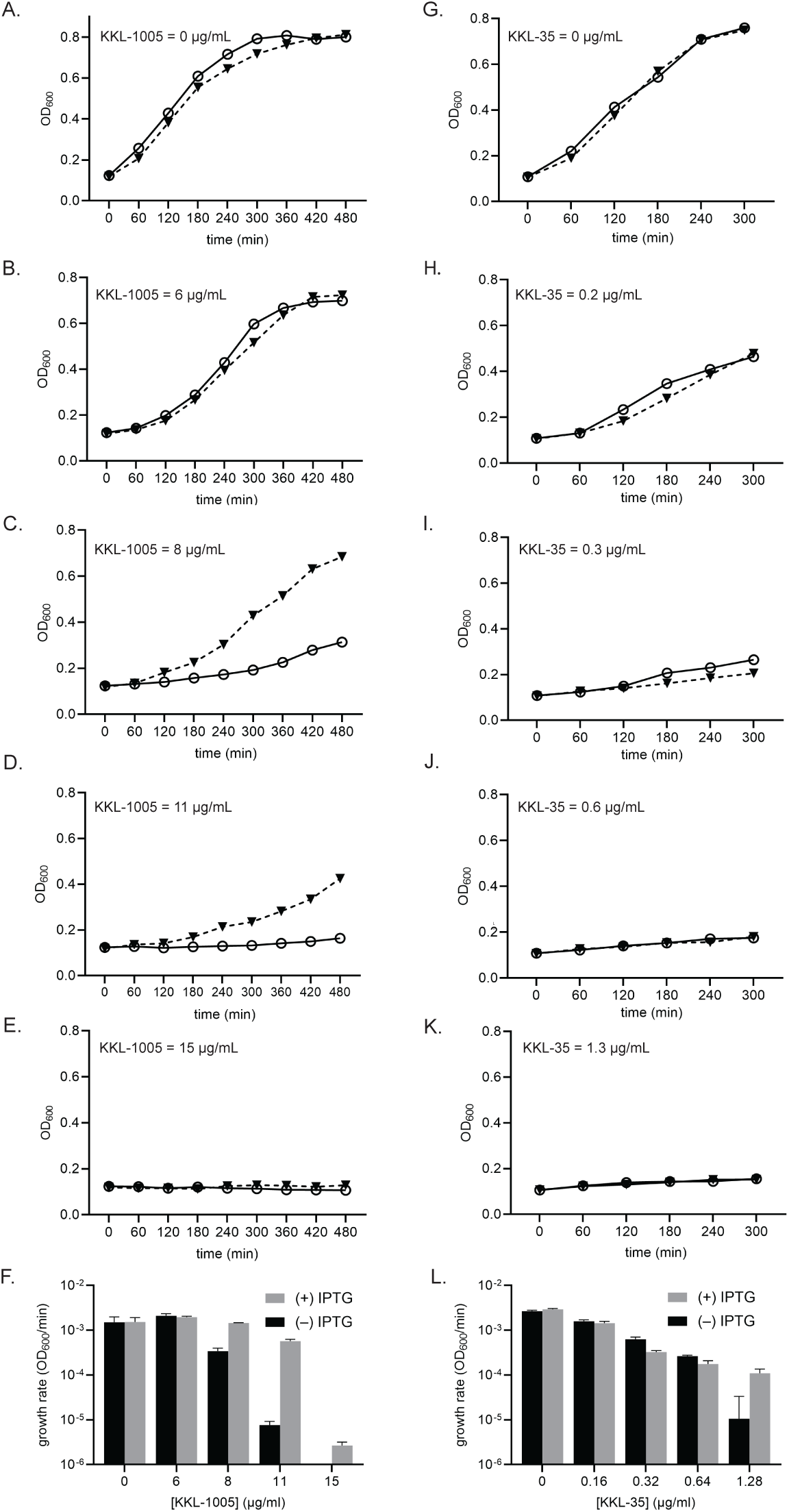
Over-expression of bL12 suppresses growth inhibition by KKL-1005. Growth curves for cultures of *E. coli ΔtolC prplL* treated with *trans*-translation inhibitors. Cultures induced to express bL12 (filled triangles) or without induction (open circles) were grown in the presence of KKL-1005 (A-E) or KKL-35 (G-K). Growth rates from exponential phase (120 – 300 min) for cultures treated with KKL-1005 (F) and KKL-35 (L) were calculated from the profiles three biological replicates and the averages with standard deviations were plotted.

### bL12 counteracts inhibition of *trans*-translation by KKL-1005 *in vitro*

If KKL-1005 inhibits bacterial growth by binding bL12 and thereby inhibits *trans*-translation, then bL12 should counteract the effect of KKL-1005 on *M. tuberculosis trans*-translation *in vitro*. To test this prediction, we added purified *M. tuberculosis* bL12 to *in vitro trans*-translation assays in the presence of KKL-1005. In the presence of 25 µM KKL-1005, the addition of 25 µM bL12 decreased *trans*-translation inhibition from 51% to 9% (Fig. 6A & B). These data suggest that KKL-1005 inhibits *trans*-translation by binding bL12, but excess bL12 can counteract the inhibitory effect of KKL-1005.

**Fig. 6:**
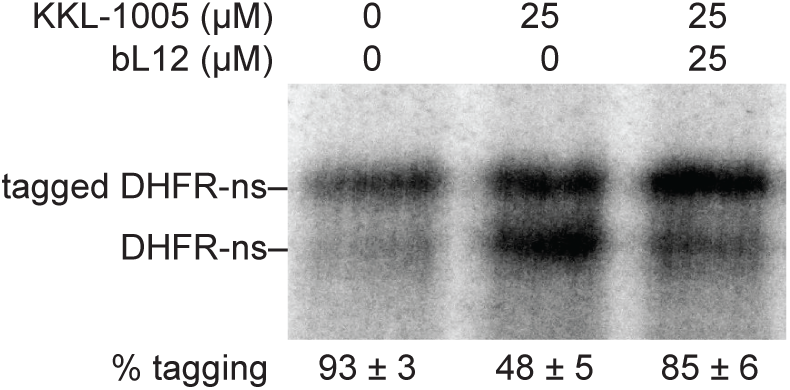
bL12 counteracts inhibition of *trans*-translation by KKL-1005 *in vitro.* *in vitro trans*-translation assays as in Figure 3, with the indicated concentrations of bL12 and KKL-1005. Reactions treated with 25 µM bL12 suppressed the inhibition of *trans*-translation by 25 µM KKL-1005. Data are the average with standard deviations from at least two experiments.

## DISCUSSION

The data presented here demonstrate that KKL-1005 inhibits *trans*-translation and kills *M. tuberculosis*. Binding and competition studies indicate that KKL-1005 exerts both effects through binding the ribosomal protein bL12 NTD. bL12 is essential in *M. tuberculosis*, and the bactericidal activity of KKL-1005 against *M. tuberculosis* suggests that the bL12 NTD is a potential target for new anti-tuberculosis drugs, although the cytotoxicity of KKL-1005 against human cells would have to be attenuated to generate useful leads based on this structure. The ability of KKL-1005 to specifically inhibit *trans*-translation and not normal translation also indicates that bL12 plays a different role with tmRNA-SmpB than with a tRNA.

How could KKL-1005 inhibit *trans*-translation and not normal translation through bL12? bL12 forms homodimers that bind uL10 and form a highly flexible stalk that is involved in the initial binding of the translation factors IF2, EF-Tu, EF-G, and RF, and stimulates their GTPase activity to promote translation initiation, elongation, and termination (Savelsbergh et al. 2000; Wahl and Moller 2002; Bocharov et al. 2004; Diaconu et al. 2005; Helgstrand et al. 2007; Huang et al. 2010; Ge et al. 2018; Younkin et al. 2020). The NTD, where KKL-1005 binds, contains the dimerization interface and uL10 binding region (Wahl et al. 2000; Diaconu et al. 2005). The CTD, connected to the NTD by a flexible linker, interacts with the GTPase translation factors (Savelsbergh et al. 2000; Wahl and Moller 2002; Bocharov et al. 2004; Kothe et al. 2004; Diaconu et al. 2005; Helgstrand et al. 2007; Huang et al. 2010; Ge et al. 2018). Mutations in the bL12 CTD impair GTPase recruitment and reduce translation fidelity (Kothe et al. 2004; Diaconu et al. 2005; Savelsbergh et al. 2005). The bL12 NTD can rotate around the L10 base and has different preferred orientations when interacting with different factors (Diaconu et al. 2005). It has been proposed that these structural changes lead to different functional states of the ribosome (Diaconu et al. 2005). Because the flexible linker between the bL12 NTD and CTD would limit the ability of a small molecule bound to the NTD to affect the interactions between the CTD and a GTPase, we think it is likely that KKL-1005 acts by changing the orientation of the NTD around L10 such that the stalk adopts a conformation that prevents *trans*-translation.

Both EF-Tu and EF-G interact with tmRNA-SmpB at points in *trans*-translation that could lead to differential inhibition through bL12. Because IF2 and RF3 are not required for *trans*-translation, it is unlikely that KKL-1005 acts by altering bL12 interaction with either of these factors. Although the tRNA-like domain of tmRNA is structurally similar to tRNA^Ala^, there are key differences in the size and flexibility of tRNA and tmRNA-SmpB that alter their interactions with the ribosome and EF-Tu. Structural studies show small differences in the path of tRNA and tmRNA along the surface of EF-Tu, but these differences are significant enough to be targeted by a different *trans*-translation inhibitor, KKL-55 (Marathe et al. 2023). KKL-55 specifically disrupts binding of EF-Tu with tmRNA but not with tRNA (Marathe et al. 2023). There are also subtle but significant differences in the timing of activation of GTP hydrolysis on EF-Tu during *trans*-translation compared to canonical translation, suggesting that there are different interactions between EF-Tu and the ribosome during these processes (Kurita et al. 2014; Miller and Buskirk 2014). It is also possible that KKL-1005 may selectively impair the EF-G–mediated translocation of the tmRNA-SmpB complex during *trans*-translation. tmRNA-SmpB is much larger than a tRNA, and translocation involves more complex movements within the ribosome (Ramrath et al. 2012). During translocation the C terminal residues of SmpB must flip within the mRNA and the tmRNA reading frame must pass through the intersubunit latches to be inserted in the mRNA channel (Ramrath et al. 2012; Guyomar et al. 2021). The role of EF-G in this process has not been studied in mechanistic detail, but it would not be surprising if there were different interactions required during *trans*-translation. We cannot exclude the possibility that KKL-1005 acts through different or multiple factors, but the activity of this inhibitor demonstrates that there are still unknown aspects of the *trans*-translation mechanism. Additional structural and biochemical experiments will be required to understand the molecular mechanism of KKL-1005.

## METHODS

### Bacterial strains, plasmids, and growth conditions

Bacterial strains, plasmids, and primer sequences are shown in Table S4. *B. anthracis* was cultured in Brain Heart Infusion Broth (Difco, Becton Dickinson, Franklin Lakes, NJ) and *E. coli* was grown in Lysogeny Broth (LB; Difco). *M. avium* strain 2285 Smooth and *M. tuberculosis* H37Rv ΔRD1 Δ*panCD* were grown in Middlebrook 7H9 medium (Difco) supplemented with 10% (v/v) OADC enrichment (Difco), 0.2% (w/v) glycerol, and 0.05% (w/v) tyloxapol. Pantothenate (50 mg/L) was added to *M. tuberculosis* cultures. *Escherichia coli* strain DH5α was used for the propagation of plasmids, and strain BL21 (DE3) was used for over-expression and purification of the ten *M. tuberculosis* translation factors, and grown in LB supplemented with 50 µg/mL kanamycin.

### Luciferase reporter screen in *E. coli*

The luciferase screen was performed as previously described (Ramadoss et al. 2013). An overnight culture of *E. coli pLC1021* was diluted to OD_600_ = 0.02 and grown to OD_600_ = 0.4 at 37 °C. luc-trpAt expression was induced with the addition of 1 mM isopropyl ß-D-1-thiogalactopyranoside (IPTG). 48 µL of cells were transferred to white 96-well plates, and 2 µL of DMSO or 20 µM compound, or 20 µM KKL-35 was added. Plates were sealed with adhesive film, vortexed gently to mix, and incubated at room temperature for 2 h. 50µL Bright-Glo reagent (Promega, Madison, WI) was added to each well, and the plates were sealed with adhesive film, vortexed vigorously, and incubated for an additional 10 min at room temperature. Luminescence was measured at 560 nm using a SpectraMax i3 microplate reader.

### MIC assays

MIC values were determined using broth dilution assays in 96-well microtiter plates per CLSI guidelines (Jorgensen et al. 2007). Plates were incubated at 37 °C (20 h for *E. coli* and *B. anthracis*, three days for *M. avium,* and one week for *M. tuberculosis*), and MIC was recorded as the lowest concentration resulting in no visible growth. Results were recorded from at least three biological repeats.

### Cytotoxicity assay

HeLa cells (ATCC) were cultured in Minimal essential media α (MEM α; Thermo Fisher Scientific, Waltham, MA) supplemented with 10% fetal bovine serum (FBS; Thermo Fisher Scientific) at 37 °C and 5% CO₂. 2,000 cells were seeded per well in a white, clear-bottom 96-well tissue culture plate (Costar) containing 100 μL of DMEM supplemented with 10% FBS and allowed to adhere overnight at 37 °C and 5% CO₂. Cells were then treated with 4 μL KKL-1005 or DMSO and incubated for 20 h at 37 °C and 5% CO₂. 74 µL media was removed from each well and 30 μL of CellTiter-Glo reagent (Promega) was added. Plate was covered with aluminum foil, gently rocked for 15 min at room temperature. Plate was allowed to equilibrate for 10 mins and luminescence was measured at 560 nm using a SpectraMax i3 microplate reader. GraphPad Prism was used to plot and fit the data to a sigmoidal function and determine the CC_50_ from three biological repeats.

### mCherry *trans*-translation reporter assays in *E. coli*

The mCherry reporter assays were performed as described previously (Ramadoss et al. 2013). Overnight cultures of *E. coli* pLC1063 *and E. coli ΔssrA* pLC1063 were diluted to OD_600_ = 0.02 and allowed to grow till OD_600_ = 0.2 at 37 °C. mCherry-trpAt expression was induced with the addition of 1 mM IPTG, and 48 µL *E. coli* pLC1063 cells or *E. coli ΔssrA* pLC1063 cells were transferred to a black 96-well plate with a clear bottom. 2 µL DMSO was added to *E. coli ΔssrA* pLC1063, and 2 µL DMSO or a varying concentration of KKL-1005 was added to *E. coli pLC*1063. The plates were incubated with shaking at 37 °C for 4 h, and mCherry fluorescence readings were recorded at α_ex_ = 560 nm & α_em_ = 617 nm on a SpectraMax i3 microplate reader. The data obtained was quantified using GraphPad Prism.

### *In vitro* translation and *trans*-translation assays

*M. tuberculosis* and *E. coli* assays were performed using purified *M. tuberculosis* and *E. coli* components based on previous protocols (Lavickova and Maerkl 2019; Varshney et al. 2025). Energy solution and *M. tuberculosis* protein solution were prepared as described previously (Varshney et al. 2025). *E. coli* protein solution was prepared as described previously (Lavickova and Maerkl 2019). *E. coli* tmRNA and *E. coli* SmpB were prepared based on published protocol (Ramadoss et al. 2013). *M. tuberculosis* tmRNA was *in vitro* transcribed using MTB ssrA F and MTB ssrA R primers based on a previous protocol. 6His-tagged *M. tuberculosis* SmpB and 6His-tagged *M. tuberculosis* EF-Tu were purified from *E. coli* BL21(DE3) pET28aTBsmpb-His_6_ and *E. coli* BL21(DE3) pET28atuf-His_6_, respectively, based on a published protocol (Varshney et al. 2025). *M. tuberculosis* ribosomes were prepared from H37Rv *ΔRD1 ΔpanCD* as described previously (Varshney et al. 2025). *E. coli* ribosomes were prepared from *E. coli* MRE600 cells using the same protocol described for the purification of *M. tuberculosis* ribosomes. *In vitro* translation reactions were set up by expressing full-length DHFR from a DHFR gene with a stop codon (DHFR-stop). Translation assays were set up using the energy solution (2 µL), *M. tuberculosis* or *E. coli* protein solution (1 µL), *M. tuberculosis* or *E. coli* EF-Tu (10 µM), *M. tuberculosis* or *E. coli* ribosomes (1.28 µM), DHFR-stop template (9 ng/µL), and [^35^S]-methionine (0.42 µCi/µL). Reactions were incubated at 37 °C for 2 h, precipitated with acetone, analyzed by SDS-PAGE followed and visualized by phosphor imaging (GE Healthcare, Chicago, IL). When necessary, KKL-1005, chloramphenicol, or DMSO was added to the reactions. Relative translation activity was calculated with respect to the DMSO-treated control. *In vitro, trans*-translation reactions were set up with some modifications to the translation assay. Full-length DHFR was expressed from a DHFR gene without an in-frame stop codon (DHFR-ns). DHFR-ns template was prepared via PCR, as described previously, and 9 ng/µL was added to the reactions. *M. tuberculosis* or *E. coli* tmRNA and SmpB were added to the reactions at final concentrations of 2.75 μM. Reactions were incubated at 37 °C for 2.5 h, precipitated with acetone, analyzed by SDS-PAGE, and visualized by phosphor imaging. When necessary, 25 µM purified *M. tuberculosis* bL12 or bL12 storage buffer was added to the reactions. Tagging efficiency was calculated as the percentage of total DHFR tagged by tmRNA-SmpB from at least 3 repeats. Dose-dependent inhibition of *trans*-translation by KKL-1005 was determined from at least three repeats. GraphPad Prism was used to plot and fit the data to a sigmoidal function and determine the IC_50_.

### Cross-linking and click conjugation

Intracellular photo-affinity labeling was performed based on a previous protocol with some modifications (Alumasa et al. 2017). 9.3 µg/mL KKL-2108 was added to a 50 mL culture of *B. anthracis* Sterne or *E. coli ΔtolC* growing in exponential phase at OD_600_ = 0.6. The culture was incubated with shaking in a water bath at 37 °C for 1 h, cells were harvested by centrifugation at 2716 *g* and resuspended in PBS (19 mM Na_2_HPO_4_ (pH 7.4), 137 mM NaCl, 2.5 mM KCl, and 2 mM KH_2_PO_4_). The suspension was irradiated with UV light (312 nm) for 10 min, cells were recovered by centrifugation at 2716 *g*, and the pellet was resuspended in lysis buffer (100 mM NaH_2_PO_4_ (pH 7.5), 100 mM NaCl, 0.1% SDS, and 2 mM β-ME). Cells were lysed by sonication and centrifuged at 22,000 *g* for 10 min. A copper-catalyzed azide−alkyne Huisgen cycloaddition click conjugation was used with KKL-2107, as described previously. Samples were analyzed by SDS-PAGE and visualized by Coomassie staining. The gel was scanned to visualize fluorescence bands using the green filter at 535 nm on Typhoon Biomolecular Imager (GE Healthcare).

### Sample preparation for Mass Spectroscopy Analysis

The excised gel fragment containing the fluorescent band was diced into smaller pieces. The gel fragments were incubated in 1 mL 45 mM DTT dissolved in 10 mM ammonium bicarbonate, pH 7.8) at 55°C for 90 min. The solution was discarded, and the gel fragments were incubated in 1 mL 100 mM iodoacetamide at room temperature in the dark for 60 min. The gel fragment was washed with 500 µl 50% 10 mM ammonium bicarbonate in acetonitrile with gentle rocking for 30 min. The liquid was discarded, and the gel fragment was air dried at room temperature in the dark. 10 mM ammonium bicarbonate containing 5 pmol trypsin was added, followed by 10 mM ammonium bicarbonate buffer to saturate the gel. Addition of this buffer continued over a 2 h period until the gel pieces were fully swollen and covered by the buffer. The sample was incubated overnight at 37°C, and an aliquot containing 5 - 50% of the solution phase was isolated. Mass mapping to identify the protein was performed in conjunction with the Huck Life Sciences Institute at Penn State University. The sample was analyzed by Matrix-assisted Laser Desorption Ionization Mass Spectroscopy (MALDI-MS), followed by MS/MS on a Thermo LTQ Orbitrap Velos ETD.

### Over-expression of L12 and time-course assays to assess growth inhibition by KKL-1005

*E. coli ΔtolC* p*rplL* was grown in LB at 37° to OD_600_ = 0.6, and over-expression of bL12 was induced by the addition of IPTG to 1 mM for 4 h at 37°C. Growth was monitored by recording OD_600_ for 6 h, and aliquots of the cultures were analyzed on 15% SDS-polyacrylamide gels. For assays involving KKL-1005, an overnight culture of this strain was diluted to OD_600_ = 0.02 and allowed to grow to OD_600_ = 0.1. The culture was and split into two equal portions and one of the flasks was induced with 1 mM IPTG. 5 ml induced or uninduced culture was dispensed into glass culture tubes containing different concentrations of the test compounds (KKL-1005: 0, 6, 8, 11, 15 µg/ml or KKL-35: 0, 0.2, 0.3, 0.6, 1.3 µg/ml), and incubated with shaking at 37 °C. Aliquots from these cultures were extracted every hour, and the OD_600_ was measured. Corresponding growth rates were calculated from the exponential phase of these profiles between T_120 min_ – T_300 min,_ and GraphPad Prism was used to plot the data.

### Western Blot Analysis

Western transfer on PVDF membrane was performed for 1 h, and the membrane was washed twice with 10 mL TBS (50 mM Tris-HCl and 150 mM sodium chloride, pH 7.6) for 10 min. The membrane was blocked with 3% w/v Bovine serum albumin (BSA) dissolved in 10 mL TBS for 1 h at room temperature and was washed thrice with 10 mL TBS. The membrane was incubated overnight at 4 °C with the 0.5 µg/mL primary antibody (Monoclonal Anti-His IgG, MilliporeSigma, Burlington, MA) dissolved in 3% w/v BSA in 10 mL TBS. Membrane was washed thrice with 10 mL TBS and incubated with the 0.03 µg/mL secondary antibody (Anti-mouse IgG A3562 GAM goat anti-mouse alkaline phosphatase, MilliporeSigma) in 1% w/v non-fat dry milk dissolved in 10 mL TBS for 1h at room temperature. Finally, the membrane was washed thrice with 10 mL TBS and incubated with 500 µL of 0.6 mg/mL ECF substrate (Cytiva, Marlborough, MA) for 2 min at room temperature. The membrane was scanned at 565 nm on the on Biorad Chemidoc XRS+ Imager.

### Purification of bL12, bL12 NTD and bL12 CTD

*M. tuberculosis* bL12 was constructed via HiFi assembly (New England Biolabs, Ipswich, MA). The template was amplified from *M. tuberculosis* H37Rv ΔRD1 Δ*panCD* genomic DNA by PCR using primers TB_L12_F and TB_L12_R and ligated into pET28a that had been digested with NcoI and XhoI. The NTD (residues 1-32) and CTD (residues 52-120) of *E. coli* bL12 were constructed via HiFi assembly. The templates were amplified from *E. coli* genomic DNA by PCR using primers L12_NTD_F, L12_NTD_R, and L12_CTD_F, L12_CTD_R, respectively. The assembled vectors were transformed into *E. coli* BL21-DE3. Overnight cultures of *E. coli prplL* or *E. coli* BL21(DE3) pET28aTB*rplL-*His_6_ or *E. coli* BL21(DE3) pET28a*rplL*-NTD-His_6_ or *E. coli* BL21(DE3) pET28a*rplL*-CTD-His_6_ was sub-cultured into 1 L terrific broth (24 g/L yeast extract, 20 g/L tryptone, 4 mL/L glycerol, 0.017 M KH_2_PO4, 0.072 M K_2_HPO_4_) and allowed to grow to OD_600_ = 0.6 with shaking at 37 °C. 6His-tagged *M. tuberculosis* bL12 or 6His-tagged *E. coli* bL12 or 6His-tagged *E. coli* NTD or 6His-tagged *E. coli* CTD was overexpressed presence of 1 mM IPTG for 3 h. The cells were harvested by centrifugation at 6,953 × g for 10 min at 4°C and resuspended in lysis buffer (50 mM Tris-HCl (pH 7.5), 60 mM NH_4_Cl, 7 mM MgCl_2_, and 5 mM β-ME). Cells were lysed by sonication, and debris was removed by centrifugation at 22,000 × g for 15 min at 4°C. The lysate was incubated with 750 µL HisPur Ni-NTA agarose resin (Thermo Fisher Scientific) for 1 h while rocking gently at 4 °C. This suspension was passed through a column, and the resin was washed thrice with wash buffer (lysis buffer at pH 6.3). Bound protein was eluted off the resin with elution buffer (lysis buffer at pH 4.0) at 4°C. Further purification was performed on a Sepharose^TM^ 10/300 size exclusion column using the lysis buffer. Purified protein was first dialyzed overnight at 4°C in bL12 storage buffer (50 mM Tris-HCl (pH 7.5), 60 mM NH_4_Cl, 7 mM MgCl_2_, 25% (v/v) glycerol, and 7 mM β-ME). Purified protein was finally dialyzed in PBS at 4°C before the MST experiments.

### Microscale Thermophoresis Experiments

600nM of 6His-tagged *E. coli L12* or 6His-tagged *M. tuberculosis* L12 or 6His-tagged *E. coli* NTD or 6His-tagged *E. coli* L12 CTD was incubated with 150nM Red-Tris-NTA dye Generation 2 (NanoTemper Technologies, Watertown, MA, USA) in binding buffer (PBS, 0.01% Tween-20, and 100mM NaCl) for 30 min at room temperature, followed by centrifugation at 21,000 *g* for 10 min at 4°C. 5 µL protein-dye supernatant was added to serially diluted 5 µL KKL-1005 samples and incubated at room temperature for 2 h. This mixture was loaded onto Monolith capillaries (Nanotemper Technologies), and MST was measured in a Monolith NT.115 (Nanotemper Technologies). Plots of concentration of KKL-1005 vs change in fluorescence of the protein were fit to the hyperbolic function y = c/ (1 + K_d_/x) in GraphPad Prism, and the K_d_ was determined from at least two repeats.

## ACKNOWLEDGEMENTS

We thank TB Alliance, Anna Upton, Khisi Mdluli, and Takushi Kaneko for providing the compound library and for many helpful discussions and insights on this project. This work was supported by NIH grants GM121650 and AI158706 and by TB Alliance.

