## Supplemental Information for "A *trans*-translation inhibitor kills *Mycobacterium tuberculosis* by targeting ribosomal protein bL12"

### Supporting Information

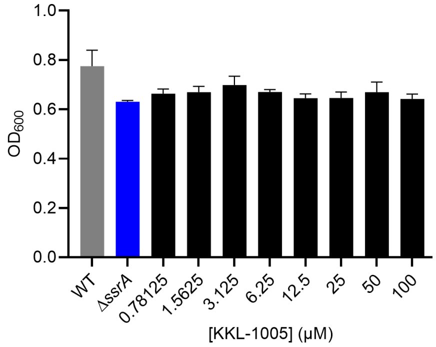


**Fig. S1: KKL-1005 does not affect cell growth of *E. coli* in mCherry assays.** *E. coli* Δ*ssrA* and wild-type were treated with DMSO, and the growth was compared to that of wild-type *E. coli* treated with KKL-1005. The final $OD_{600}$ for all cultures used in Figure 1B. Data are the mean with the error bars representing the standard deviation for three biological replicates.

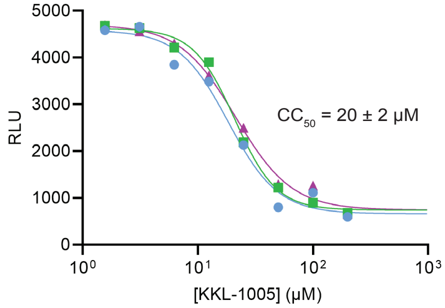


**Fig. S2: KKL-1005 is cytotoxic against HeLa cells.** HeLa cells were treated with KKL-1005 or DMSO. Relative luminescence units (RLU) represent the luminescence relative to that of the DMSO treated control. Data were plotted and fit to a sigmoidal function and the mean $CC_{50}$ with standard deviation from three biological replicates is shown.

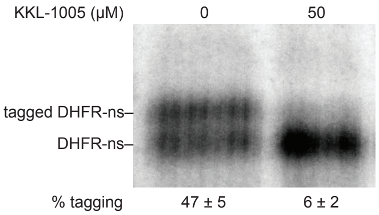


**Fig. S3: KKL-1005 inhibits *E. coli trans*-translation *in vitro.*** *In vitro trans*-translation assay consisting of *E. coli* components and DHFR without a stop codon was incubated with 50μM KKL-1005 or DMSO***.*** Bands corresponding to tagged and untagged DHFR are indicated, and the average percentage of DHFR protein found in the tagged band for two repeats is shown with standard deviation.

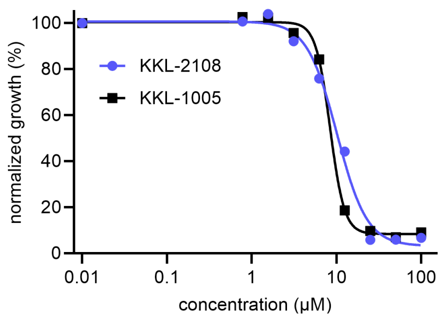


**Fig. S4: Growth of *E. coli* ∆*tolC* is inhibited by both KKL-2108 and KKL-1005.** The dose-response curves for the growth inhibition of *E. coli* after treatment with KKL-2108 or KKL-1005.

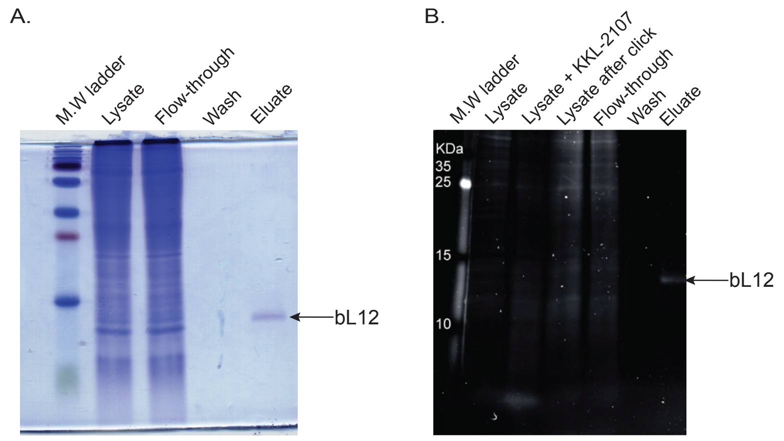


**Fig. S5: KKL-1005 binds to *E. coli* bL12 *in vivo*.** Click chemistry as in Fig. 3 with *E. coli* ∆*tolC*. A) Coomassie stained gel analysis of the affinity chromatography shown in Fig. 3B. C) Fluorescence scan of a gel analysis of the samples from (A) with controls for the lysate premixed with KKL-2107 only and lysate after the click reaction.

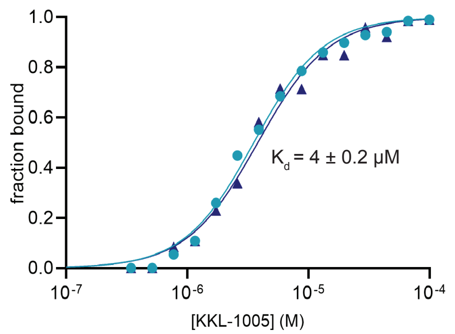


**Fig. S6: KKL-1005 binds *M. tuberculosis* bL12 *in vitro*.** MST was used to measure the binding of KKL-1005 with *M. tuberculosis* bL12. Change in fluorescence was measured for fluorescently labeled bL12 with KKL-1005, and the fraction of bL12 bound to KKL-1005 was calculated. Data were plotted and fit to a sigmoidal function in GraphPad Prism and the mean $K_d$ with standard deviation from three technical replicates is shown.

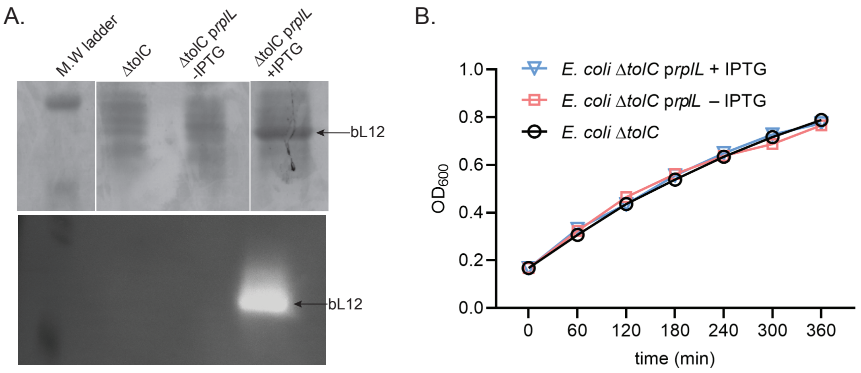


**Fig. S7: Over-expression of bL12 does not interfere with the growth of *E. coli*.** A) SDS-PAGE analysis of *E. coli* Δ*tolC* and *E. coli* Δ*tolC* p*rplL* lysates showing a Coomassie-stained gel (top) and a western blot (bottom) confirming over-expression of bL12 in the IPTG-induced culture. B) Growth curves for the *E. coli* Δ*tolC* and *E. coli* Δ*tolC* p*rplL* with and without IPTG.

### Supplementary tables

#### Table S1. Luciferase luminescence activity normalized (%) to KKL-35

| Compound I.D | SMILES | average (%)[1] |
|---|---|---|
| KKL-0035 | FC1=CC=C(C2=NN=C(NC(=O)C3=CC=C(Cl)C=C3)O2)C=C1 | 100 |
| KKL-1005 | O=C(Nc1nc(NC(=O)c2ccccc2)nn1-c1ccccc1)c1ccco1 | 44 |
| KKL-0003 | OC1=CC([N+]([O-])=O)=CC2=CC=CN=C12 | 46 |
| KKL-0062 | COC1=CC=CC=C1C(=O)NNC(=S)NC(=O)C1=CC=CS1 | 45 |
| KKL-0063 | CC(=O)NC(=S)NNC(=O)C1=CC=C(C2=CC=CC=C2)C=C1 | 99 |
| KKL-0065 | CC(=O)NC(=S)NNC(=O)C1=CC=C(C)C(Br)=C1 | 186 |
| KKL-0093 | OC1=CC=C(Br)C2=CC=CN=C12 | 371 |
| KKL-0209 | FC(F)(F)c1cccc(C(=O)Nc2nnc(-c3ccc(Cl)cc3)o2)c1 | 24 |
| KKL-0287 | Cc1nc(CSc2nc(C)nc3sc4c(c23)CCCC4)cs1 | 0 |
| KKL-0288 | CN(C)C(S)=Nc1nc2ccccc2s1 | 2 |
| KKL-0289 | Cc1cc(C)nc(/N=C(\N)Nc2cc(Cl)ccc2Oc2ccccc2)n1 | 0 |
| KKL-0290 | Cc1cc(C(=O)Oc2ccc(Cc3ccccc3)cc2)ccc1[N+]([O-])=O | 0 |

| | | |
|---|---|---|
| KKL-0291 | [O-][N+](=O)c1ccc2[nH]c(CSc3nc4ccccc4o3)nc2c1 | 0 |
| KKL-0292 | Cn1c(Sc2cnn(-c3ccccc3)c(=O)c2Br)nnc1-c1ccccc1 | 0 |
| KKL-0293 | Cc1cccc(OCc2nnc(SCC(=O)NCCc3ccccc3)o2)c1 | 0 |
| KKL-0294 | CCCS(=O)(=O)c1nnnn1-c1cccc(OC)c1 | 0 |
| KKL-0295 | CCn1c(/C=C/c2ccc([N+]([O-])=O)o2)nc2ccc(I)cc2c1=O | 0 |
| KKL-0296 | Oc1ccc([N+]([O-])=O)cc1/C=C/C(=O)c1ccc(Cl)cc1 | 0 |
| KKL-0297 | CCCCc1cc(=O)oc2cc(OC(C)C(=O)OCC)c(Cl)cc12 | 0 |
| KKL-0298 | CCOC(=O)c1c(CSc2ccccc2)n(-c2ccccc2)c2ccc(O)c(CN(C)C)c12 | 0 |
| KKL-0299 | Cc1ccnc(NC(=O)COc2ccc([N+]([O-])=O)cc2)c1 | 0 |
| KKL-0300 | CCOC(=O)C(Oc1ccc2oc(C)c(C(=O)OC)c2c1)c1ccccc1 | 0 |
| KKL-0301 | CC(C)CC(=O)OCC1=CO[C@@H](OC(=O)CC(C)C)C2C1C[C@H ](OC(C)=O)[C@]21CO1 | 1 |
| KKL-0302 | COC(=O)[C@@H]1Cc2c([nH]c3ccccc23)[C@H]2C[C@H](NCc3c ccnc3)C[C@@H](c3ccc(F)cc3)N12 | 0 |
| KKL-0303 | CN(C)c1ccc(- c2cc([C@H]3CN4CC[C@H]3C[C@@H]4CNC(=S)Nc3ccc(C)cc3 [illegible] | 0 |
| KKL-0304 | CC(C)[C@H]1OC(=O)[C@H](Cc2ccccc2)N(C)C(=O)[C@@H](C( C)C)OC(=O)[C@H](Cc2ccccc2)N(C)C(=O)[C@@H](C(C)C)OC(= [illegible] | 0 |
| KKL-0305 | COC(=O)[C@@H]1Cc2c([nH]c3ccccc23)[C@H]2C[C@@H](N3C COCC3)C[C@@H](c3ccc(Cl)c(F)c3)N12 | 0 |
| KKL-0306 | COC(=O)[C@@H]1Cc2c([nH]c3ccccc23)[C@H]2C[C@H](NCCN C(C)=O)C[C@@H](c3ccc(OC(F)(F)F)cc3)N12 | 20 |
| KKL-0307 | FC(F)(F)c1ccc(C(=O)NC[C@H]2C[C@@H]3CCN2C[C@@H]3C N2CCC(Cc3ccccc3)CC2)cc1 | 0 |
| KKL-0308 | COC(=O)[C@@H]1Cc2c([nH]c3ccccc23)[C@H]2C[C@H](NCCN 3CCOCC3)C[C@@H](c3ccc(OC(F)(F)F)cc3)N12 | 6 |
| KKL-0309 | CN(C)c1ccc(CN[C@H]2C[C@H](c3c(C)nn(- c4ccc(C(F)(F)F)cc4)c3C)C=C2)cc1 | 0 |
| KKL-0310 | CC(C)(C)c1ccc(C(=O)Nc2ccc(NC(=S)NC(=O)c3cccs3)cc2)cc1 | 0 |
| KKL-0311 | CCCCc1ccc(NC(=S)NC(=O)c2ccc(OCC)c(Cl)c2)cc1 | 0 |
| KKL-0312 | COC(=O)[C@@H]1Cc2c([nH]c3ccccc23)[C@H]2C[C@@H](N3C C[C@@H](O)C3)C[C@@H](c3ccc(C(F)(F)F)cc3)N12 | 2 |
| KKL-0313 | COC(=O)[C@@H]1Cc2c([nH]c3ccccc23)[C@H]2C[C@@H](N3C COCC3)C[C@@H](c3ccc(OC(F)(F)F)cc3)N12 | 4 |
| KKL-0314 | COC(=O)[C@@H]1Cc2c([nH]c3ccccc23)[C@H]2C[C@H](NCCN 3CCOCC3)C[C@@H](c3ccc(Cl)c(F)c3)N12 | 9 |
| KKL-0315 | COc1cccc(- c2cc(CC3(O)CCN(Cc4[nH]c5ccc(C)cc5c4C)CC3)on2)c1 | 0 |
| KKL-0316 | COc1ccc(-c2ncc(CN3CCCC(C(=O)c4ccccc4SC)C3)cn2)cc1 | 0 |
| KKL-0317 | CSc1ncc(CNC2CCCc3c2cnn3-c2ccc(C(C)(C)C)cc2)cn1 | 0 |
| KKL-0318 | Cc1ccc(NC2CCCN(C3CSCCSC3)C2)cc1C | 6 |
| KKL-0319 | CN(Cc1cc(C)[nH]n1)Cc1cn(-c2ccc(C)cc2)nc1-c1ccc(F)cc1 | 0 |

| | | |
|---|---|---|
| KKL-0320 | Cc1ccc(-n2cc(CN3CCN(c4ccccn4)CC3)c(-c3ccc(F)cc3)n2)cc1 | 0 |
| KKL-0321 | COc1ccc(CNCc2cn(-c3cccc(F)c3)nc2-c2ccc(C)o2)cc1 | 0 |
| KKL-0322 | C1CCC(c2ccc(-c3cnnc(N4CCN(c5ccncc5)CC4)n3)cc2)CC1 | 0 |
| KKL-0323 | Cn1nc(-c2ccc3ccccc3c2)cc1[C@H]1CN2CC[C@H]1C[C@@H]2COC(=O [illegible] | 0 |
| KKL-0324 | COc1ccc(-c2cc([C@H]3CN4CC[C@H]3C[C@@H]4COC(=O)Nc3cccc(C(F)( [illegible] | 0 |
| KKL-0325 | Cc1nc(C)c(-c2csc(Nc3ccc(Br)cc3)n2)s1 | 0 |
| KKL-0326 | NC1CCc2nc(-c3ccncc3)nc(Nc3ccc(Cl)cc3)c2C1 | 797 |
| KKL-0327 | COc1ccccc1CN(CCCn1ccnc1)Cc1ccc(Cl)c(Cl)c1 | 16 |
| KKL-0328 | COc1ccc2c(c1)NC1(COC3(C1)CCN(Cc1cccc(Cl)c1)CC3)c1cccn1-2 | 0 |
| KKL-0329 | Cc1ccc(OC(=O)c2ccccc2Cl)cc1C | 0 |
| KKL-0330 | Cc1cc(CN2CCC(O)(c3ccccc3)C2)ccc1N1CCCC1 | 0 |
| KKL-0331 | Fc1ccccc1-c1csc(NC(=O)c2cnc(N3CCCCC3)nc2)n1 | 2 |
| KKL-0332 | CCN(C(=O)C1CCCCC1)c1nc2c(s1)CCc1oc(C)cc1-2 | 0 |
| KKL-0333 | OC1(c2ccccc2)CCN(Cc2ccc(N3CCCC3)c(F)c2)C1 | 0 |
| KKL-0334 | CCCn1nc(C)c(-c2cc(C(=O)Nc3ccc(CN4CCCCCC4)cc3)[nH]n2)c1C | 0 |
| KKL-0335 | Cc1nc(C2CCNCC2)ncc1-c1nc(N2CCCCCC2)ncc1-c1ccc(F)cc1 | 0 |
| KKL-0336 | O[C@@H]1CCCN(C(=O)Nc2ccc(OC(F)(F)F)cc2)C[C@H]1NC(=O)c1ccc(OCCN2CCCC2)cc1 | 0 |
| KKL-0337 | Cc1ccc(C(=O)NC[C@H]2C[C@@H]3CCN2C[C@@H]3c2cc(-c3ccc4ccccc4c3)nn2C)cc1 | 0 |
| KKL-0338 | COc1ccc2c(c1)C=C(CN(CCCn1ccnc1)Cc1ccc(C)o1)CO2 | 0 |
| KKL-0339 | CSc1ccccc1C(=O)C1CCCN(C(=O)c2ccc3[n+]([O-])onc3c2)C1 | 0 |
| KKL-0340 | Cc1c(C(=O)NCCN2CCC(O)(c3ccc(C)cc3)CC2)oc2ccccc12 | 1 |
| KKL-0341 | COc1cc(CN(CCCn2ccnc2)Cc2ccc(C)s2)cc2c1OCO2 | 9 |
| KKL-0342 | Cc1nn(-c2ccc(F)cc2)c2nc(O)c(CNCc3ccccc3F)cc12 | 0 |
| KKL-0343 | CC(C)OC1c2ncc(-c3ccccc3)cc2Sc2nc(-c3cccnc3)nn21 | 0 |
| KKL-0344 | CSc1ncc(CN2CCCC(C(=O)c3ccc(-c4ccccc4)c(F)c3)C2)cn1 | 0 |
| KKL-0345 | COc1ccc(CN2CCC(O)(c3cccnc3)CC2)cc1COc1ccccc1F | 3 |
| KKL-0346 | CSc1nccc(N2CCCC(CCC(=O)NCc3ccc(F)c(F)c3)C2)n1 | 0 |
| KKL-0347 | CC(C)(C)C(Cn1ccnc1)NCc1c[nH]nc1-c1ccc(-c2ccccc2)cc1 | 0 |
| KKL-0348 | CC1CCc2onc(C(=O)NC3CCCc4c3cnn4Cc3ccccc3Cl)c2C1 | 0 |

| | | |
|---|---|---|
| KKL-0349 | Cc1nn(-c2ccccc2)c2nc(O)c(CNc3cc4c(cc3C)OCCO4)cc12 | 0 |
| KKL-0350 | Cc1nn(-c2ccccc2)c2nc(O)c(CNC(c3ccco3)c3ccccc3)cc12 | 0 |
| KKL-0351 | CN(CCn1ccnc1)Cc1cn(-c2ccc(C)cc2)nc1-c1cccc(C)c1 | 0 |
| KKL-0352 | COc1ccc(C(=O)C(CN2CCOCC2)c2ccccc2)cc1 | 0 |
| KKL-0353 | Nc1ncnc(Oc2cccc3cccnc23)c1[N+]([O-])=O | 0 |
| KKL-0354 | COc1cc(CNCc2ccccn2)ccc1OCc1ccc(Cl)c(Cl)c1 | 0 |
| KKL-0355 | CC(C)(C)c1ccc(-c2nc(NCc3cccnc3)c3ccccc3n2)cc1 | 1 |
| KKL-0356 | [O-<br>][N+](=O)c1ccc(C(=O)Nc2ccc(N3CCN(C(=O)c4ccc(Cl)cc4)CC3)c<br>[illegible] | 0 |
| KKL-0357 | CN1CCN(c2ccc(NC(=S)NC(=O)c3ccc(-c4cccc(Cl)c4)o3)cc2)CC1 | 18 |
| KKL-0358 | Cc1cc2c(cc1C)N1C(=NCCC1)N2Cc1ccc(C(C)(C)C)cc1 | 0 |
| KKL-0359 | CCn1c(NC(=O)c2ccccc2)nc2ccccc12 | 6 |
| KKL-0360 | COc1ccccc1/C=C/C(=O)Oc1cccc2cccnc12 | 0 |
| KKL-0361 | C1CN(c2nc(NC3C4CC5CC(C4)CC3C5)c3ccccc3n2)CCO1 | 0 |
| KKL-0362 | [O-<br>][N+](=O)c1cc(C(=O)N(CCCCN2C(=O)c3ccccc3C2=O)c2ccc(Cl)c<br>[illegible] | 0 |
| KKL-0363 | CC1=NN(c2nc3ccccc3s2)C(=O)C1Cc1cccc2ccccc12 | 9 |
| KKL-0364 | Cc1ccc(Nc2nc(NCc3ccco3)c3ccccc3n2)cc1 | 19 |
| KKL-0365 | [O-][N+](=O)c1ccc2c3c(cccc13)C(=O)N(Cc1cccnc1)C2=O | 0 |
| KKL-0366 | Brc1ccc(/C=C(/NC(=O)c2ccccc2)C(=O)NCC(=O)OCc2ccccc2)cc<br>1 | 2 |
| KKL-0367 | OC[C@@H](NC(=O)C(Cl)Cl)[C@H](O)c1ccc([N+]([O-])=O)cc1 | 0 |
| KKL-0368 | Brc1ccc(-c2nnc(SCC#CCOC(=O)c3cccs3)o2)cc1 | 0 |
| KKL-0369 | C(Nc1nc(-c2ccccn2)nc2ccccc12)c1ccco1 | 0 |
| KKL-0370 | CC(=O)n1cc(N(C(=O)CCl)c2ccccc2)c2ccccc12 | 0 |
| KKL-0371 | CN(C)CCCNc1cc(C(C)(C)C)nc2c(-c3ccccc3)cnn12 | 0 |
| KKL-0372 | CCOC(=O)c1cnc(-n2nc(C)cc2C)nc1N(CC)CC | 0 |
| KKL-0373 | CCCCCCCn1c(NC(=O)CC)nc2ccccc12 | 27 |
| KKL-0374 | Cn1c(-c2ccccc2)cc2c1ccc1[n+]([O-])onc21 | 0 |
| KKL-0375 | C1CN(c2nc(-c3ccccn3)nc3ccccc23)CCN1c1ccccc1 | 0 |
| KKL-0376 | Cc1cc2ncn(N=Cc3ccc([N+]([O-])=O)s3)c2cc1C | 32 |
| KKL-0377 | Cc1ccc(-c2c[n+](CC(=O)Nc3nc(-c4ccc(F)cc4)cs3)c3n2CCC3)cc1 | 21 |

| | | |
|---|---|---|
| KKL-0378 | Clc1ccc2c(c1)C1=C(C(c3ccncc3)O2)C(c2ccc(Br)cc2)n2c(ncn2)N1 | 0 |
| KKL-0379 | CC(=O)Nc1nc2ccccc2n1Cc1ccccc1 | 0 |
| KKL-0380 | CCCCOC(=O)CSc1nc(N)c(C#N)c(-c2ccc(OC)cc2)c1C#N | 3 |
| KKL-0381 | Clc1ccc(COC(Cn2ccnc2)c2ccc(Cl)cc2Cl)c(Cl)c1 | 0 |
| KKL-0382 | [O-][N+](=O)c1ccc2nc(NC(NC(=O)c3ccccc3Cl)(C(F)(F)F)C(F)(F)F)sc[illegible] | 0 |
| KKL-0383 | Clc1ccc(/C=C/C(=O)c2cccnc2)c(Cl)c1 | 0 |
| KKL-0384 | OC12CC3CC(C1)CC(C(=O)Nc1nc(-c4ccccn4)cs1)(C3)C2 | 0 |
| KKL-0385 | CCOC(=O)C1=C(c2ccccc2)N=C(N)NC1c1ccc(OCc2ccc(Cl)cc2)c(OC)c1 | 6 |
| KKL-0386 | [O-][N+](=O)c1cc(C(=O)OC(c2ccccc2)c2nccc3ccccc23)cc([N+]([O-[illegible] | 1 |
| KKL-0387 | Cc1cccc(Nc2nnc(SCC(=O)Nc3cccnc3Cl)s2)c1 | 0 |
| KKL-0388 | Brc1ccccc1-c1nnc(SCC(=O)N2CCc3ccccc3C2)o1 | 0 |
| KKL-0389 | Cc1ccc2nc(Cl)c(/C=C/C(=O)c3ccco3)cc2c1 | 0 |
| KKL-0390 | Clc1cnc(C(=O)Nc2nc(-c3ccccn3)cs2)c(Cl)c1Cl | 10 |
| KKL-0391 | Oc1c(CN2CCc3ccccc3C2)cc(Cl)c2cccnc12 | 42 |
| KKL-0392 | O=C(/C=C/c1ccc(C#N)cc1)c1cccnc1 | 0 |
| KKL-0393 | Clc1ccc(/C=C/C(=O)c2cccnc2)cc1 | 0 |
| KKL-0394 | CCOC(=O)C1CCN(C2=NC(=O)/C(=C\c3ccc(Cl)cc3Cl)S2)CC1 | 0 |
| KKL-0395 | COc1cc2cc(-c3nc4ccccc4[nH]3)c(N)nc2cc1OC | 22 |
| KKL-0396 | OC(CC(=O)Nc1cccc(C(F)(F)F)c1)(C(F)(F)F)C(F)(F)F | 28 |
| KKL-0397 | CC[C@@H]1C[C@H]2CC[C@H](O2)[C@H](C)C(=O)O[C@@H](C)C[C@@H]2CC[C@@H](O2)[C@@H](C)C(=O)O[C@H](C)C[C[illegible] | 0 |
| KKL-0398 | Cc1ccc(C(=O)N2CCN(c3ccc(NC(=O)COc4c(C)cc(C)cc4Br)cc3)CC2)cc1 | 3 |
| KKL-0399 | [O-][N+](=O)c1ccc(NCCNc2nc3ccccc3nc2N2CCOCC2)cc1 | 9 |
| KKL-0400 | Cc1cc2ncn(CCOc3ccc(Cl)cc3)c2cc1C | 2 |
| KKL-0401 | Cc1ccc(NCCC(=O)c2ccc([N+]([O-])=O)cc2)cc1C | 14 |
| KKL-0402 | COc1cc2c(cc1NC(=O)Nc1ccc(Cl)cc1)oc1ccccc21 | 5 |
| KKL-0403 | CC(C)COc1ccc(-c2nn(-c3ccccc3)cc2C(=O)Nc2nccs2)cc1 | 5 |
| KKL-0404 | CCOC(=O)COc1ccc(OCc2ccccc2)cc1 | 1 |
| KKL-0405 | Brc1ccc(-c2nn(-c3ccccc3)cc2CNCc2ccccc2)cc1 | 0 |
| KKL-0406 | Cc1ccc(Cl)cc1N1CCN(CC(O)COc2ccc(C(=O)c3ccccc3)cc2)CC1 | 0 |

| | | |
|---|---|---|
| KKL-0407 | COC(=O)COc1ccc2c(-c3ccc(OC)cc3)cc(=O)oc2c1 | 0 |
| KKL-0408 | Fc1ccc(NC(=S)NC(=O)c2ccc(Br)cc2)cc1 | 0 |
| KKL-0409 | Cc1cc(C)c(C(=O)/C=C/c2cccc([N+]([O-])=O)c2)cc1C | 0 |
| KKL-0410 | CC1=CC(C)(C)Nc2ccc(OC(=O)c3cccs3)cc21 | 0 |
| KKL-0411 | CCCOc1ccccc1NC(=O)Nc1ccc(Cl)c(Cl)c1 | 2 |
| KKL-0412 | CCN(c1ccccc1)c1ccc([N+]([O-])=O)c2nonc12 | 0 |
| KKL-0413 | Clc1cccc(NC(=O)CSc2ccc(Br)cc2)c1 | 8 |
| KKL-0414 | COC(=O)C(C)Oc1ccc2c(c1)oc(C)c(-c1ccc(Br)cc1)c2=O | 9 |
| KKL-0415 | Clc1ccc(OC(=O)c2cccs2)c2ncccc12 | 4 |
| KKL-0416 | CC(=NNC1=NC(=O)C(Cc2ccc(F)cc2)S1)c1ccc(Cl)cc1 | 10 |
| KKL-0417 | Clc1ccc(OCCCNCc2ccccc2)c2ccccc12 | 48 |
| KKL-0418 | COc1ccc(-c2[nH]c(-c3ccc([N+]([O-])=O)cc3)nc2-c2ccccc2)cc1 | 11 |
| KKL-0419 | CCC(C)Oc1ccc(C(=O)NC(=S)Nc2ccc(N3CCOCC3)cc2)cc1 | 0 |
| KKL-0420 | COC(C)(C)C#CC(=O)c1cccc2ccccc12 | 0 |
| KKL-0421 | Clc1ccc(OCCCCCn2ccnc2)c(Cl)c1 | 0 |
| KKL-0422 | CC(=NNc1nc(C)c(C)s1)c1ccc(Br)cc1 | 0 |
| KKL-0423 | CC(C)c1ccc(OCCCCn2ccnc2)cc1C | 8 |
| KKL-0424 | [O-][N+](=O)c1ccc(C(=O)Nc2ccccc2Br)o1 | 0 |
| KKL-0425 | Cc1nn(-c2ccc([N+]([O-<br>])=O)cc2)c(C)c1N1C(=O)c2cccc3cc(Br)cc(c23)C1=O | 0 |
| KKL-0426 | COc1ccccc1-c1c(C)oc2cc(OCC(=O)OCc3ccccc3)ccc2c1=O | 0 |
| KKL-0427 | CCC(C)NC(=O)c1ccc(NC(=O)C2(c3ccccc3)CCCC2)cc1 | 0 |
| KKL-0428 | COc1ccc(-c2nnc(SCC(=O)NCCc3ccccc3)o2)cc1 | 4 |
| KKL-0429 | CC(C)c1ccc(C)c(OCCCCn2ccnc2)c1 | 8 |
| KKL-0430 | CC(=O)Oc1cccc2ccc(/C=C/c3cccc(Br)c3)nc12 | 14 |
| KKL-0431 | FC(F)C(F)(F)Oc1ccccc1NC(=O)Nc1cccc(Cl)c1 | 21 |
| KKL-0432 | Cc1ccccc1/C=C/c1ccc2cccc(O)c2n1 | 11 |
| KKL-0433 | [O-][N+](=O)c1ccc(-c2nnc(SCC#C)o2)cc1 | 8 |
| KKL-0434 | CCOC(=O)COc1ccc2c(c1)oc(C)c(-c1ccc(OC)cc1)c2=O | 0 |
| KKL-0435 | Fc1ccccc1C(=O)Oc1cccc2cccnc12 | 0 |

| | | |
|---|---|---|
| KKL-0436 | CCC(C)c1ccc(O)c(NC(=S)NC(=O)c2cccnc2)c1 | 17 |
| KKL-0437 | FC(F)(F)c1cc(NCCC(=O)c2cccs2)ccc1Cl | 0 |
| KKL-0438 | Clc1ccc(C2C=C(c3ccc(Cl)cc3)Nc3nnnn32)cc1 | 0 |
| KKL-0439 | Cc1ccc(CN(CCCn2ccnc2)Cc2cccn2-c2cccc(F)c2)o1 | 9 |
| KKL-0440 | Cc1c(CN2CCC3(CC2)NC(=O)c2ccccc2O3)[nH]c2ccc(F)cc12 | 0 |
| KKL-0441 | O=C1Oc2ccccc2N1C1CCN(Cc2nc3ccccc3n2Cc2ccccc2)CC1 | 2 |
| KKL-0442 | O=C(CC(c1ccccc1)c1ccccc1)N1CCN(Cc2ccc3c(c2)OCO3)CC1 | 7 |
| KKL-0443 | OC(CN1CCCCCC1)Cn1cc(/C=C/C(=O)c2cccs2)c2ccccc12 | 0 |
| KKL-0444 | Clc1ccc(N2CCN(CCCNC(=S)Nc3cccc(Cl)c3)CC2)cc1 | 10 |
| KKL-0445 | CC(C)Cn1ncc2c1CCc1sc(N(C)C(=O)c3noc4c3CCCC4)nc1-2 | 0 |
| KKL-0446 | COc1ccc2c(c1)NC1(COC3(C1)CCN(Cc1ccccc1Cl)CC3)c1cccn1-2 | 3 |
| KKL-0447 | Fc1ccc(CNc2nc(-c3ccncc3)cc3ncccc23)cc1 | 0 |
| KKL-0448 | CC1=CCC(c2cnc(-c3ccccc3)c(C)c2)NC1c1ccccc1 | 0 |
| KKL-0449 | CN(C)C[C@@H]1CN(Cc2ccc(OCC=C)cc2)C[C@H]1c1ccnc(N2CCCC2)n1 | 0 |
| KKL-0450 | O=C(Oc1cccc2cccnc12)c1ccccc1 | 3 |
| KKL-0451 | CC1=Nc2c(ccc3cccnc23)C1(C)C | 0 |
| KKL-0452 | COC(=O)CSc1cc(SCC(=O)OC)c(C#N)c(SCC(=O)OC)c1 | 0 |
| KKL-0453 | Cc1nn(CCCC(=O)Nc2cc(Oc3ccc(C)cc3)cc([N+]([O-])=O)c2)c(C)c1[N+]([O-])=O | 0 |
| KKL-0454 | CCOc1cc(/C=C2\SC(=N)N(c3nccs3)C2=O)ccc1OCCN1CCOCC1 | 0 |
| KKL-0455 | Oc1c(C(=O)Nc2cccnc2)c(=O)n2c3c(cccc13)CC2 | 0 |
| KKL-0456 | Cc1ccc(C(=O)CC(=O)c2cc(Cl)ccc2O)cc1 | 0 |
| KKL-0457 | O=C(Nc1cccc(-c2ccco2)c1)C1CCN(C2Cc3ccccc3C2)CC1 | 0 |
| KKL-0458 | Cc1ccc(-c2nn(-c3cccc(F)c3)cc2CNCc2cnn(Cc3ccccc3)c2)o1 | 0 |
| KKL-0459 | CCn1c(C)cc2cc(NC(=O)c3cnc(SC)nc3)ccc12 | 3 |
| KKL-0460 | CCOc1ccc2[nH]c3c(c2c1)CN(Cc1cccc(OCC#C)c1)CC3 | 26 |
| KKL-0461 | Cc1ccc2c(c1)C(O)CC1(O2)CCN(CC(O)c2ccc3c(c2)CCCC3)CC1 | 0 |
| KKL-0462 | O=C(Cn1cccc2cc(-c3ccco3)nc12)NCCOc1ccccc1 | 4 |
| KKL-0463 | COc1ccc(Cc2cnc(NC(=O)c3ccccc3Cn3ccnc3)s2)cc1 | 6 |
| KKL-0464 | CC(C)C1=Nc2sc(C)c(C)c2C(=O)N1C1CCN(C(=O)c2ccncc2)CC1 | 14 |

| | | |
|---|---|---|
| KKL-0465 | CSc1ncc(C(=O)NC2(c3ccccc3)CCCC2)cn1 | 0 |
| KKL-0466 | COc1cc(C)c(-c2csc(NC(=O)c3cnc(SC)nc3)n2)cc1C | 0 |
| KKL-0467 | CCOc1cccc(-c2csc(NC(=O)c3cnc(SC)nc3)n2)c1 | 0 |
| KKL-0468 | COc1ccccc1CCCNC(=O)c1cc(-c2c(C)nn(C)c2C)[nH]n1 | 0 |
| KKL-0469 | COc1ccccc1C1(O)CCN(Cc2[nH]c3ccc(F)cc3c2C)CC1 | 4 |
| KKL-0470 | COc1ccc(C)cc1CN1CCC(c2cc3ncccc3cn2)CC1 | 0 |
| KKL-0471 | O=C(Nc1ccccc1)C1COc2ccccc2O1 | 6 |
| KKL-0472 | CN(C)CCOc1ccccc1C(=O)Nc1nc(-c2ccc(F)cc2)c(C)s1 | 0 |
| KKL-0473 | COc1ccc2[nH]c(CN3CCC4(CC3)NC(=O)c3ccccc3O4)c(C)c2c1 | 9 |
| KKL-0474 | CSc1ncc(C(=O)Nc2nc(-c3c[nH]c4cc(C)ccc34)cs2)cn1 | 0 |
| KKL-0475 | CN(Cc1nccs1)Cc1cn(-c2ccc(C)cc2)nc1-c1ccc(F)cc1 | 2 |
| KKL-0476 | COc1ccc(F)c(-c2nc(CN3CCC4(CC3)C=Cc3ccccc34)c(C)o2)c1 | 6 |
| KKL-0477 | O=C(Nc1ccc(-n2cccn2)cc1)C1CCCCN1Cc1cccc2nonc12 | 1 |
| KKL-0478 | COc1ccc(F)c(-c2nc(CN3CCN(c4ncc5ccccc5n4)CC3)c(C)o2)c1 | 3 |
| KKL-0479 | Clc1ccc(C(=O)C2CCCN(Cc3ccc4nonc4c3)C2)cc1 | 0 |
| KKL-0480 | CCOc1ccc(C2N(Cc3cnc(SC)nc3)CCc3c2[nH]c2ccccc32)cc1 | 0 |
| KKL-0481 | COc1ccc(NC(=O)N2CCCC(C(=O)c3cc(F)ccc3OC)C2)cc1Cl | 0 |
| KKL-0482 | Cc1ccc(NC2CCCN(C(=O)c3ccc4[n+]([O-])onc4c3)C2)cc1C | 3 |
| KKL-0483 | C(N1CCN(c2nc(-c3ccccc3)nc3c2CCC3)CC1)c1cc2ccccc2[nH]1 | 0 |
| KKL-0484 | Cc1oc(-c2cccs2)nc1CN1CCc2c([nH]c3ccccc23)C1c1ccccn1 | 6 |
| KKL-0485 | Cc1ccc(-<br>c2cc(C(=O)N3CCCC(N4CCN(c5ccccc5C)CC4)C3)[nH]n2)s1 | 0 |
| KKL-0486 | CCOC(=O)C1CCCN(Cc2nc(-c3ccc(Cl)cc3)no2)C1 | 2 |
| KKL-0487 | COc1ccc(-<br>c2nc(CN3CCC4(CC3)C=Cc3ccccc34)c(C)o2)c(OC)c1C | 0 |
| KKL-0488 | Cc1nc2sccn2c1C(=O)N1CCc2c([nH]c3ccccc23)C1c1ccc(F)cc1 | 4 |
| KKL-0489 | CN(Cc1cn(-c2ccc(C)cc2)nc1-c1ccc(F)cc1)Cc1ccccn1 | 1 |
| KKL-0490 | CCC(CC)C(=O)N1CCCC(C(=O)Nc2cccc(-c3ccc(F)cc3)c2)C1 | 3 |
| KKL-0491 | Cn1nc2c(c1C(=O)N1CCc3c([nH]c4ccccc34)C1c1ccccc1F)CCCC<br>2 | 0 |
| KKL-0492 | COC(=O)c1ccc(-c2noc(CN3CCCC(c4nc5ccccc5o4)C3)n2)cc1 | 3 |
| KKL-0493 | CSc1ccc(C(=O)C2CCCN(C(=O)c3ccc4[n+]([O-])onc4c3)C2)cc1 | 3 |

| | | |
|---|---|---|
| KKL-0494 | COc1ccccc1CN1CCCCC1C(=O)Nc1ccc(-n2cccn2)cc1 | 3 |
| KKL-0495 | COC(=O)[C@@H](NC(=O)c1cc(COc2ccc(F)cc2Cl)on1)c1ccccc1 | 2 |
| KKL-0496 | COC(=O)[C@@H](NC(=O)c1cc(COc2c(F)cccc2F)on1)c1ccccc1 | 8 |
| KKL-0497 | COC(=O)C(CCSC)NC(=O)c1cc(COc2ccc(F)cc2Cl)on1 | 9 |
| KKL-0498 | Cc1cccc(-n2cc(CN3CCN(c4ccncc4)CC3)c(-c3cccc(F)c3)n2)c1 | 0 |
| KKL-0499 | CSCCC(=O)N1CCCC(C(=O)Nc2cccc(-c3cccc(C)c3)c2)C1 | 0 |
| KKL-0500 | CC(NCc1cn(-c2ccc(C)cc2)nc1-c1cccc(C)c1)c1csc(C)n1 | 0 |
| KKL-0501 | COc1cccc(CNCc2cn(-c3ccc(C)cc3)nc2-c2ccc(F)cc2)c1OC | 0 |
| KKL-0502 | CC(C)(C)c1ccc(OCC(=O)N2CCCC(c3cc4ncccc4cn3)C2)cc1 | 0 |
| KKL-0503 | C[C@H]1CN(C(=O)c2cc(-c3cccs3)on2)[C@@H](C)CN1Cc1ccco1 | 0 |
| KKL-0504 | COc1ccc2c(c1)CCCN2Cc1ccnc(N2CCCC2)n1 | 4 |
| KKL-0505 | COC(=O)[C@@H](NC(=O)c1cc(COc2cncc(Cl)c2)on1)c1ccccc1 | 12 |
| KKL-0506 | CC(C(=O)N1CCC(Oc2ncnc3n(C4CCCC4)c(C)c(C)c23)CC1)c1ccccc1 | 0 |
| KKL-0507 | CC(C)Cc1cc(CNC(=O)c2cc(-c3c(C)nn(C)c3C)[nH]n2)on1 | 0 |
| KKL-0508 | COc1cccc(O)c1CN1CCC(C(=O)Nc2ccc(-c3cccc(C)c3)cc2)CC1 | 0 |
| KKL-0509 | Cc1ccc(CN(CCCn2ccnc2)Cc2cc3c(cc2Cl)OCO3)s1 | 2 |
| KKL-0510 | Cc1c[nH]c(CCNCc2cn(-c3cccc(C)c3)nc2-c2cccc(F)c2)n1 | 9 |
| KKL-0511 | Cc1cc2nc(C3CCCN(C(=O)c4[nH]c5ccccc5c4C)C3)cc(C(F)(F)F)n2n1 | 0 |
| KKL-0512 | COc1ccc(C(C)NC(=O)c2cc(-c3ccccc3OC)[nH]n2)cc1F | 0 |
| KKL-0513 | CCc1c(C)sc2c1C(=O)N(C1CCN(C(=O)c3nccc4ccccc34)CC1)C=N2 | 3 |
| KKL-0514 | COc1cccc(CN2CCC(c3c[nH]c4ccc(OC)cc34)C2)c1 | 2 |
| KKL-0515 | CCC1N(Cc2cnn(Cc3ccccc3)c2C)CCn2c(C)ccc21 | 0 |
| KKL-0516 | CCC1N(Cc2cccc(OCC#C)c2)CCn2cccc21 | 0 |
| KKL-0517 | COc1ccc2[nH]c(CN3CCc4c([nH]c5ccccc45)C3C)cc2c1 | 5 |
| KKL-0518 | Fc1ccccc1CN(CCCn1ccnc1)CC1=Cc2ccccc2OC1 | 4 |
| KKL-0519 | CCOC(=O)C1(Cc2ccc(Cl)cc2)CCN(C(=O)Nc2ccc(SC)cc2)CC1 | 2 |
| KKL-0520 | COc1ccc(-c2cnnc(NCC(C)Oc3cccnc3)n2)c(OC)c1 | 3 |
| KKL-0521 | Cc1ccc(CNCc2c(C(=O)N3CCCCCCC3)nc3ccc(Cl)cn23)o1 | 2 |
| KKL-0522 | CN(Cc1cn(-c2ccc(C)cc2)nc1-c1cccc(C)c1)Cc1nccs1 | 8 |

| | | |
|---|---|---|
| KKL-0523 | CN(C)CC(C)(C)CNc1cc(-c2ccccc2)nc2cc(-c3cccc(F)c3)nn12 | 0 |
| KKL-0524 | Cc1cc(OCC(=O)NC[C@H]2CC[C@H](c3nnc(-c4cccs4)o3)CC2)ccc1Cl | 0 |
| KKL-0525 | CC1CCc2c(sc3nc(C(C)(C)C)nc(N4CCNCC4)c23)C1 | 0 |
| KKL-0526 | NC1CCc2nc(-c3cccnc3)nc(Nc3ccc(C(F)(F)F)cc3)c2C1 | 137 |
| KKL-0527 | COc1ccc(C2(C)NN=C3Sc4ccccc4N23)cc1 | 5 |
| KKL-0528 | COc1ccc(Cl)cc1NC(=O)CN1CCC(c2nc3cc(Cl)ccc3[nH]2)CC1 | 0 |
| KKL-0529 | COc1cc(/C=C/C2=CCN(C)CC2)ccc1OCC(=O)Nc1cccc(Cl)c1 | 3 |
| KKL-0530 | NC1CCc2nc(-c3cccnc3)nc(Nc3cccc(C(F)(F)F)c3)c2C1 | 124 |
| KKL-0531 | CCc1c(C)nc2cc(-c3cccs3)nn2c1NCc1ccccc1OC | 0 |
| KKL-0532 | Cc1ccc2[nH]c(-c3ccc(NCc4cc5cc(C)ccc5nc4O)cc3)nc2c1 | 0 |
| KKL-0533 | Cc1ccc2n(C)c(C(=O)N3CCCC(c4cc5[nH]cccc5n4)C3)cc2c1 | 0 |
| KKL-0534 | Cc1ccc2[nH]c(-c3ccc(NCc4cc5ccc(C)cc5nc4O)cc3)nc2c1 | 0 |
| KKL-0535 | CC(Cc1nc2ccccc2[nH]1)Cc1cc(C)cc(C)c1 | 1 |
| KKL-0536 | CCOC(=O)c1ccc2ncc(C(=O)OCC)c(NCc3cccnc3)c2c1 | 0 |
| KKL-0537 | COc1ccc(CN2CCC(c3cc4nc(C)cc(C(F)(F)F)n4n3)C2)c(OC)c1 | 0 |
| KKL-0538 | Cc1ccc(-c2csc(Nc3ccc(N)cc3)n2)cc1 | 0 |
| KKL-0539 | NC1CCc2nc(-c3ccncc3)nc(Nc3ccc(C(F)(F)F)cc3)c2C1 | 620 |
| KKL-0540 | NC1CCc2nc(-c3cccnc3)nc(Nc3ccc(Cl)cc3)c2C1 | 119 |
| KKL-0541 | NC1CCc2nc(-c3ccncc3)nc(Nc3cccc(C(F)(F)F)c3)c2C1 | 802 |
| KKL-0542 | OC(CN1CCNCC1)Cn1c2ccc(Cl)cc2c2cc(Cl)ccc12 | 0 |
| KKL-0543 | Cc1ccc(CN(CCCn2ccnc2)Cc2oc3ccccc3c2C)s1 | 3 |
| KKL-0544 | Cc1oc2ccccc2c1CN(CCCn1ccnc1)Cc1cccc(F)c1 | 0 |
| KKL-0545 | Cc1c(CN(CCCn2ccnc2)Cc2ccc(F)cc2)[nH]c2ccc(F)cc12 | 2 |
| KKL-0546 | C[C@H]1CN(C(=O)c2cc(-c3ccc4c(c3)OCO4)on2)[C@@H](C)CN1Cc1ccco1 | 0 |
| KKL-0547 | COCc1nnc(NC(=O)c2nnn(-c3ccccc3F)c2C)s1 | 0 |
| KKL-0548 | COc1cccc(N2CCCN(Cc3cn(C=C)nc3C)CC2)c1 | 0 |
| KKL-0549 | O[C@@H]1CCCN(C(=O)Nc2ccc(OC(F)(F)F)cc2)C[C@H]1NC(=O)c1ccc(OCc2ccccc2)cc1 | 1 |
| KKL-0550 | C[C@H]1[C@@H]2CC[C@@]3(C)Cc4sc(NC(=O)Nc5ccc(F)cc5F)nc4[C@@H](C)[C@@H]3[C@H]2OC1=O | 0 |
| KKL-0551 | COC(=O)[C@@H]1Cc2c([nH]c3ccccc23)[C@H]2C[C@@H](NCCN(C)C)C[C@@H](c3ccc(C(C)C)cc3)N12 | 0 |

| | | |
|---|---|---|
| KKL-0552 | OC(CN1CCC2(CC1)CC(O)c1cc(Cl)ccc1O2)c1ccc2ccccc2c1 | 0 |
| KKL-0553 | COc1ccccc1C1(O)CCN(Cc2cnn(-c3ccc(C)cc3)c2)CC1 | 7 |
| KKL-0554 | FC(F)(F)c1cccc(Nc2ncccc2C(=O)NCCCn2ccnc2)c1 | 15 |
| KKL-0555 | CSc1ncc(C(=O)N2CCc3c([nH]c4ccc(Cl)cc34)C2)cn1 | 0 |
| KKL-0556 | COc1ccc2c(c1)C(O)CC1(O2)CCN(CC(O)c2ccc3c(c2)CCCC3)CC1 | 0 |
| KKL-0557 | Cc1c(CN2CCC(O)(c3cccc(C)c3)CC2)[nH]c2ccccc12 | 2 |
| KKL-0558 | Fc1ccc(OCc2cc(C(=O)N3CCCCO3)no2)c(Cl)c1 | 0 |
| KKL-0559 | COc1ccccc1-c1csc(NC(=O)c2cccc(Cn3nc(C)cc3C)c2)n1 | 3 |
| KKL-0560 | Cc1ccc2oc(C(=O)N3CCC(Oc4cccnc4)CC3)cc2c1 | 4 |
| KKL-0561 | COc1ccccc1C(=O)NNC(=S)NC(=O)c1ccc(-c2ccccc2)cc1 | 11 |
| KKL-0562 | COc1ccc(NC(=O)c2ccc(C#CC(C)(C)O)s2)cc1 | 1 |
| KKL-0563 | CCOC(=O)c1c(NC(=S)NC(=O)N(C)C)scc1-c1ccc(C)o1 | 0 |
| KKL-0564 | CC(C)(C)c1ccc(C(=O)NC(=S)NCc2ccco2)cc1 | 0 |
| KKL-0565 | COC(=O)COc1ccc2cc(-c3ccc(OC)c(OC)c3)c(=O)oc2c1 | 3 |
| KKL-0566 | Clc1ccc(Cn2ccc(NC(=O)c3ccc4c(c3)OCO4)n2)cc1 | 0 |
| KKL-0567 | Cc1cc(NC(=O)CSc2nnnn2-c2ccccc2C)no1 | 0 |
| KKL-0568 | FC(F)(F)C1CC(c2cccs2)Nc2c(C(=O)NCc3ccc4c(c3)OCO4)cnn21 | 0 |
| KKL-0569 | CC1(C)C2(C)CCC1(C(=O)Nc1ccc(OC(F)(F)F)cc1)CC2=O | 11 |
| KKL-0570 | Brc1ccccc1-c1nc2sc3c(c2c(=O)o1)CCCC3 | 0 |
| KKL-0571 | O=C(Oc1cccc2cccnc12)c1cc2ccccc2oc1=O | 8 |
| KKL-0572 | CSc1nsc(SCC(=O)N2CCN(c3ccccc3)CC2)n1 | 0 |
| KKL-0573 | C(N1CNC(Nc2nc3ccccc3s2)=NC1)c1ccc2c(c1)OCO2 | 2 |
| KKL-0574 | Cc1cc(C)cc(NC(=O)COc2cccc(C=O)c2)c1 | 1 |
| KKL-0575 | Brc1ccc(C(=O)Nc2nc3c(s2)CCCC3)s1 | 0 |
| KKL-0576 | Cc1cccc(-n2nc([N+]([O-])=O)c(NCCC3=CCCCC3)[n+]2[O-])c1 | 6 |
| KKL-0577 | Clc1ccc(NC(=O)NC2CC3CCC2C3)cc1 | 0 |
| KKL-0578 | CN1CCC(O)(c2cccs2)C(C(=O)c2cccs2)C1 | 3 |
| KKL-0579 | Cc1ccc2[nH]c(/C(=C\c3c[nH]c4ccc(Br)cc34)C#N)nc2c1 | 0 |
| KKL-0580 | Fc1ccc(C(=O)NC(=S)N(CC=C)CC=C)cc1 | 20 |

| | | |
|---|---|---|
| KKL-0581 | O=C(Oc1ccccc1)c1ccc(S(=O)(=O)N2CCCCC2)cc1 | 2 |
| KKL-0582 | O=C(NC(=S)N(CCC#N)Cc1ccccc1)c1ccccc1 | 14 |
| KKL-0583 | CSc1ccccc1NC(=O)c1ccc(Br)s1 | 0 |
| KKL-0584 | CCOC(=O)c1ccc(NC(=O)c2ccc(C)s2)cc1 | 0 |
| KKL-0585 | O=C(Oc1ccccc1)c1cccc2cccnc12 | 0 |
| KKL-0586 | O=C(Nc1nccs1)c1ccc(CSc2ccccc2)o1 | 0 |
| KKL-0587 | CCOC(=O)CSC1=C(C#N)C(c2ccccc2[N+]([O-])=O)C(C(C)=O)=C(C)N1 | 0 |
| KKL-0588 | CC(=O)NC(=S)NNC(=O)c1ccc(C(C)(C)C)cc1 | 143 |
| KKL-0589 | CCOC(=O)CSc1n[nH]c(-c2ccc(Cl)cc2)n1 | 0 |
| KKL-0590 | COc1nc(SC)nc2sc3c(c12)CCC3 | 4 |
| KKL-0591 | CC(NC(=O)NNC(=O)c1ccncc1)(C(F)(F)F)C(F)(F)F | 0 |
| KKL-0592 | CCOC(=O)COc1c(Cl)cc(/C=C(/C#N)c2nc3cc(OC)ccc3[nH]2)cc1Cl | 0 |
| KKL-0593 | CC(C)NS(=O)(=O)c1ccc(NC(=S)NC(=O)c2ccc(C)c(Br)c2)cc1 | 0 |
| KKL-0594 | COc1cccc(NC(=O)c2c(NC(=O)c3cccnc3)sc3c2CCC3)c1 | 0 |
| KKL-0595 | O=C(CSc1ccccc1)Nc1ccc(-c2nc3ccccc3c(=O)o2)cc1 | 1 |
| KKL-0596 | O=C(CCc1ccccc1)NC(=S)Nc1ccc(N2CCCCC2)cc1 | 1 |
| KKL-0597 | [O-][N+](=O)c1ccc(/C=C(/C#N)c2ccc(C#N)cc2)o1 | 19 |
| KKL-0598 | COC(=O)CSc1nnc(-c2ccc(Cl)cc2)o1 | 0 |
| KKL-0599 | Clc1ccc(C(=O)NC(=S)N2CCN(C/C=C/c3ccccc3)CC2)cc1 | 33 |
| KKL-0600 | COc1ccccc1NC(=O)CSc1nc2cc3ccccc3cc2[nH]1 | 1 |
| KKL-0601 | CCN(CC)c1cccc(Oc2nc([N+]([O-])=O)nn2C)c1 | 0 |
| KKL-0602 | CCOC(=O)Cn1cc(/C=C(/C#N)c2nc3cc(OC)ccc3[nH]2)c2ccccc12 | 2 |
| KKL-0603 | COc1ccc2[nH]cc(CCN=C(S)Nc3ccc(Nc4ccccc4)cc3)c2c1 | 2 |
| KKL-0604 | Cc1ccc(C(=O)Nc2ccc([N+]([O-])=O)cc2)s1 | 0 |
| KKL-0605 | CSc1ncc(C(=O)Nc2cccc(C(F)(F)F)c2)cn1 | 3 |
| KKL-0606 | CCc1nc2sc(C(=O)NCc3ccc(F)cc3)c(N)c2c(C(F)(F)F)c1C | 1 |
| KKL-0607 | O=C(CSc1nc2c(cc1C#N)CCC2)OCc1ccccc1 | 0 |
| KKL-0608 | Cn1c2c(c(=O)n(C)c1=O)C(C(F)(F)F)(C(F)(F)F)NC(c1ccc(Cl)cc1)=N2 | 10 |
| KKL-0609 | O=C(Oc1cccc2cccnc12)c1ccc(S(=O)(=O)N2CCCCC2)cc1 | 0 |

| | | |
|---|---|---|
| KKL-0610 | COc1cccc(-n2nnnc2S(C)(=O)=O)c1 | 0 |
| KKL-0611 | CCc1ccc(Oc2nc([N+]([O-])=O)nn2C)cc1 | 0 |
| KKL-0612 | O=C(CSc1nc2cc3ccccc3cc2[nH]1)Nc1ccccc1C#N | 0 |
| KKL-0613 | Cc1scc(C(=O)NCc2ccco2)c1-c1ccccc1 | 1 |
| KKL-0614 | CCC(C(=O)Nc1ccc(CC(=O)OC)cc1)c1ccccc1 | 9 |
| KKL-0615 | Cc1ccc(-c2nnc3sc(Cc4ccccc4)nn23)cc1 | 0 |
| KKL-0616 | O=C(/C=C/c1cccs1)N1CCc2ccc(NC(=O)c3ccncc3)cc2C1 | 0 |
| KKL-0617 | Fc1ccc(OCc2cc(C(=O)N3CCCCO3)no2)cc1F | 0 |
| KKL-0618 | CN(CC(O)COc1ccc(CNCc2cc3ccccc3s2)cc1)Cc1ccccc1 | 0 |
| KKL-0619 | CCOC(=O)C1(CCOC)CCN(Cc2ccc(N(C)c3ccccc3)cc2)CC1 | 2 |
| KKL-0620 | Cc1ccccc1C1N(C(=O)c2nn(C)c(=O)c3ccccc23)CCc2c1[nH]c1ccccc21 | 1 |
| KKL-0621 | CN(Cc1cccc(-n2cccn2)c1)C(=O)c1ccc2nc(C3CCCCC3)oc2c1 | 0 |
| KKL-0622 | CSCCC(=O)N1CCC(Oc2ccc(C(=O)NCCc3ccccc3)cc2)CC1 | 4 |
| KKL-0623 | Fc1ccccc1C1N(C(=O)c2cc(=O)c3ccccc3o2)CCc2c1[nH]c1ccccc21 | 0 |
| KKL-0624 | OC(CC(=O)Nc1ccc(Cl)c(Cl)c1)(C(F)(F)F)C(F)(F)F | 36 |
| KKL-0625 | Cc1ccccc1N1CCN(C2CCCN(C(=O)c3ccc4[n+]([O-])onc4c3)C2)CC1 | 2 |
| KKL-0626 | [O-][n+]1onc2cc(C(=O)N3CCCC(N4CCN(c5ccc(F)cc5)CC4)C3)ccc1[illegible] | 4 |
| KKL-0627 | COc1cccc(OC)c1-c1cnnc(NCc2cccc(SC)c2)n1 | 1 |
| KKL-0628 | CC(C)n1cc(CN2CCC(C(=O)Nc3cccc(-c4cccc(Cl)c4)c3)CC2)cn1 | 0 |
| KKL-0629 | CSc1ccc(-c2nc(CN3CCN(c4ccc(Cl)cn4)CC3)c(C)o2)cc1 | 0 |
| KKL-0630 | CSc1nccc(N2CCCC(CCC(=O)NCc3ccc(C)o3)C2)n1 | 0 |
| KKL-0631 | [O-][N+](=O)c1ccc(C(=O)Nc2ccc(N3CCN(C(=O)c4ccccc4)CC3)cc2)[illegible] | 0 |
| KKL-0632 | OC1(Cc2cc(-c3ccccc3)no2)CCN(Cc2ccc(OC3CCCC3)cc2)CC1 | 0 |
| KKL-0633 | COc1ccc(-c2ncc(CN3CCCC(C(=O)c4cccc(OC(C)C)c4)C3)cn2)cc1 | 0 |
| KKL-0634 | Fc1cccc(-n2cc(CNC(=O)c3cc(COc4cccc5cccnc45)on3)cn2)c1 | 2 |
| KKL-0635 | CSCCC(=O)N1CCCC(C(=O)Nc2cccc(-c3ccc(F)cc3)c2)C1 | 9 |
| KKL-0636 | CC1(C)CC(NC(=O)c2ccc3[n+]([O-])onc3c2)c2cnn(-c3cccc(F)c3)c2C1 | 5 |
| KKL-0637 | O=C(N1CCCCC1c1cccnc1)c1ccc2oc(Cc3ccccc3)nc2c1 | 4 |
| KKL-0638 | COc1cccc(-c2cc(CC3(O)CCN(CC4=CC[C@@H](C(C)=C)CC4)CC3)on2)c1 | 1 |

| | | |
|---|---|---|
| KKL-0639 | CC(C)Cc1ccc(CN2CCCC(NC(=O)c3ccc4[n+]([O-])onc4c3)C2)cc1 | 5 |
| KKL-0640 | Cn1cc(CNCc2ccc3c(c2)OCO3)c(-c2ccc(C3CCCCC3)cc2)n1 | 0 |
| KKL-0641 | CSCCC(=O)N1CCCC(CNC(=O)c2ccc(-c3ccccc3)cc2)C1 | 2 |
| KKL-0642 | Cc1cn2c(C(=O)N3CCc4c([nH]c5ccccc45)C3c3cccc(C)c3)csc2n1 | 3 |
| KKL-0643 | Fc1cccc(Cl)c1C1N(Cc2ncc[nH]2)CCc2c1[nH]c1ccccc21 | 6 |
| KKL-0644 | COc1ccc(C2N(Cc3cc4ccccc4nc3O)CCc3c2[nH]c2ccccc32)cc1C | 0 |
| KKL-0645 | Cc1cccc(C2N(Cc3ccc4cc(F)ccc4n3)CCc3c2[nH]c2ccccc32)n1 | 0 |
| KKL-0646 | COc1ccc(-c2nc(CN3CCC4(CC3)C=Cc3ccccc34)c(C)o2)c(F)c1 | 2 |
| KKL-0647 | COc1ccc(-c2cnnc(NCCc3cccnc3)n2)cc1 | 2 |
| KKL-0648 | CN(Cc1cn(-c2cccc(C)c2)nc1-c1cccc(F)c1)Cc1cccnc1 | 12 |
| KKL-0649 | CN(CCn1ccnc1)Cc1cn(Cc2ccccc2)nc1-c1cc2ccccc2o1 | 0 |
| KKL-0650 | COc1cccc(CNC(=O)CCC2CCCN(c3ccnc(SC)n3)C2)c1 | 9 |
| KKL-0651 | Cc1ccc(-n2cc(CN3CCC(CO)(Cc4cccc(C(F)(F)F)c4)CC3)cn2)cc1 | 0 |
| KKL-0652 | CSc1ccc(NC(=O)N2CCC(CO)(CCOc3ccccc3)CC2)cc1 | 3 |
| KKL-0653 | COc1cccc(-c2nn(-c3ccccc3)cc2CNCc2ccc(-n3cncn3)cc2)c1 | 2 |
| KKL-0654 | CN(CCn1ccnc1)Cc1cn(-c2ccc(C)cc2)nc1-c1ccc(F)cc1 | 3 |
| KKL-0655 | CCOc1ccccc1C1N(Cc2cc3ccccc3nc2O)CCc2c1[nH]c1ccccc21 | 9 |
| KKL-0656 | Cc1ccc(-c2nn(-c3cccc(F)c3)cc2CNCc2[nH]nc3c2CCCCC3)o1 | 0 |
| KKL-0657 | CCn1ccnc1CN(C)Cc1cn(-c2ccc(C)cc2)nc1-c1ccc(F)cc1 | 0 |
| KKL-0658 | CSCCCNC(=O)C1CCN(c2nc(-c3ccccc3)nc3c2CCC3)CC1 | 2 |
| KKL-0659 | Cc1oc(-c2ccc3ccccc3c2)nc1CN1CCN(c2cnccn2)CC1 | 0 |
| KKL-0660 | COc1ccc(O)c(CN2CCCC(N3CCN(c4cccc(C(F)(F)F)c4)CC3)C2)c1 | 1 |
| KKL-0661 | Cc1ccc(-c2[nH]ncc2CNCc2ccc(OC(F)(F)F)cc2)cc1C | 9 |
| KKL-0662 | Cc1ccc(-n2cc(CNCCCn3ccnc3)c(-c3cccc(C)c3)n2)cc1 | 0 |
| KKL-0663 | COc1ccc2cc(C(=O)C3CCCN(Cc4cnc(SC)nc4)C3)ccc2c1 | 2 |
| KKL-0664 | CSc1ccc(C(=O)C2CCCN(Cc3cc4ccccc4[nH]3)C2)cc1 | 0 |
| KKL-0665 | CCn1ccnc1CN(C)Cc1cn(-c2ccc(C)cc2)nc1-c1cccc(C)c1 | 0 |
| KKL-0666 | CCOC(=O)C1CCN(Cc2nc(-c3ccc(Cl)cc3)no2)CC1 | 6 |
| KKL-0667 | C1CCC(c2ccc(-c3cnnc(N4CCN(c5cnccn5)CC4)n3)cc2)CC1 | 0 |

| | | |
|---|---|---|
| KKL-0668 | CC(C)Cc1cc(C(=O)N2CCc3c([nH]c4ccccc34)C2c2ccccc2F)nc(N)n1 | 0 |
| KKL-0669 | Cc1c(CN2CCC(CO)(CCOc3ccccc3)CC2)[nH]c2c(Cl)cccc12 | 0 |
| KKL-0670 | CCc1sc(NC(=O)c2cnc(SC)nc2)nc1-c1ccc(F)cc1 | 5 |
| KKL-0671 | CSc1ncc(C(=O)Nc2ccccc2OCC=C)cn1 | 0 |
| KKL-0672 | CSc1ncc(C(=O)N2CCC(C3CCCCC3)C2)cn1 | 0 |
| KKL-0673 | CC(C)(C)c1ncc(C(=O)N2CCc3[nH]c4c(Cl)cccc4c3C2)cn1 | 1 |
| KKL-0674 | Cc1cc2c(cc1C)C(O)CC1(O2)CCN(CC(O)c2ccc3ccccc3c2)CC1 | 0 |
| KKL-0675 | Cc1ccc2c(C)nc(Nc3nc(O)c4ccccc4n3)nc2c1 | 1 |
| KKL-0676 | CC(C)Oc1cccc(C(=O)C2CCCN(C3CSCCSC3)C2)c1 | 0 |
| KKL-0677 | COC(=O)[C@@H](NC(=O)c1cc(COc2ccccc2)on1)c1ccccc1 | 2 |
| KKL-0678 | COc1cccc(C2(NC(=O)c3cnc(SC)nc3)CCCC2)c1 | 4 |
| KKL-0679 | CSc1ncc(C(=O)Nc2cccc(-c3csnn3)c2)cn1 | 0 |
| KKL-0680 | CN(Cc1ccnc(N2CCCC2)n1)Cc1cccc2ncccc12 | 0 |
| KKL-0681 | Cc1ccc(C(O)CN2CCC3(CC2)CC(O)c2cc(Cl)ccc2O3)cc1C | 0 |
| KKL-0682 | O=C(CN1N=C2C(=CC1=O)CCc1nc(N3CCCCC3)ncc12)OC1CCCCC1 | 0 |
| KKL-0683 | COCc1nnc(NC(=O)c2cc(-c3ccc(F)cc3)n(C)n2)s1 | 0 |
| KKL-0684 | Cc1cccc(-c2cc3ncccc3c(NCc3ccccn3)n2)c1 | 0 |
| KKL-0685 | Cc1c(C(=O)NCCCc2ccc(F)cc2)[nH]nc1-c1ccccc1 | 0 |
| KKL-0686 | CSc1ncc(C(=O)NC(C(C)C)c2cccs2)cn1 | 0 |
| KKL-0687 | COc1ccc(-c2cc(NC(=O)c3cnc(SC)nc3)n(C)n2)cc1 | 0 |
| KKL-0688 | COc1ccccc1C1(O)CCN(CC(O)c2ccc3ccccc3c2)CC1 | 3 |
| KKL-0689 | CCCOc1ccc(N2CC(C(=O)Nc3cc(C)ccn3)CC2=O)cc1 | 0 |
| KKL-0690 | Cc1ccc2cc3c(nc2c1C)OC(=O)N(C1CCCC1)C3 | 1 |
| KKL-0691 | CC(NC(=O)c1cc2cc(F)ccc2[nH]1)c1cc(O)n2nc(C3CCOCC3)cc2n1 | 4 |
| KKL-0692 | Cc1ccc(C(NC(=O)C2(c3ccc(Cl)cc3)CCOCC2)c2cccnc2)cc1 | 19 |
| KKL-0693 | Fc1ccc(CCCNC(=O)c2cc(-c3ccccn3)[nH]n2)cc1 | 0 |
| KKL-0694 | CSc1ncc(C(=O)NC(C)c2ccc3c(c2)CCC3)cn1 | 5 |
| KKL-0695 | CCCC(NC(=O)c1cnc(SC)nc1)c1cccs1 | 2 |
| KKL-0696 | O=C(Nc1ccc(Oc2cccnc2)cc1)C1Oc2ccccc2O1 | 0 |

| | | |
|---|---|---|
| KKL-0697 | Cc1ccc(NC(=O)C2OCOc3ccccc32)cc1N1CCCCC1 | 0 |
| KKL-0698 | Cc1ccc2cc(C(N3CCN(c4cccc(C)c4C)CC3)c3nnnn3C(C)(C)C)c(O)nc2c1 | 0 |
| KKL-0699 | Cc1cc2c(cc1C)C(O)CC1(O2)CCN(CC(O)c2ccc3c(c2)CCC3)CC1 | 0 |
| KKL-0700 | C(N1CCCC2(C1)NCCc1c2[nH]c2ccccc12)c1ccccc1 | 0 |
| KKL-0701 | CN(C)CCN(Cc1ccc2c(c1)OCO2)C(=O)c1cc2cc(Cl)ccc2n1C | 0 |
| KKL-0702 | CC1CC1C(=O)N1CCC(c2nc(Nc3cccc(F)c3)cc(-c3cccnc3)n2)CC1 | 6 |
| KKL-0703 | COc1ccccc1CC(C)NC(=O)c1cnc(SC)nc1 | 0 |
| KKL-0704 | CCC(NC(=O)c1cnc(SC)nc1)c1ccc(F)cc1 | 0 |
| KKL-0705 | Cc1nn(C)c(C)c1-c1cc(C(=O)NCCCc2ccc(F)cc2)[nH]n1 | 2 |
| KKL-0706 | OC(CN1CCC2(CC1)CC(O)c1cc(Cl)ccc1O2)c1ccc2c(c1)CCC2 | 1 |
| KKL-0707 | COc1ccccc1C1(O)CCN(Cc2[nH]c3ccc(C)cc3c2C)CC1 | 7 |
| KKL-0708 | Fc1ccc(CN(CCCn2ccnc2)Cc2ccccc2F)cc1F | 7 |
| KKL-0709 | O=C(CN1c2c(cnn2-c2ccccc2)C2=NCCN2C1=O)N1CCc2ccccc21 | 0 |
| KKL-0710 | CSc1ncc(C(=O)Nc2ccc3[nH]c4c(c3c2)CCCC4)cn1 | 0 |
| KKL-0711 | CCC(C)c1ccc(O)c(NC(=S)NC(=O)c2cncc(Br)c2)c1 | 3 |
| KKL-0712 | [O-][N+](=O)c1ccc(C(=O)Nc2ccc(Cl)cc2C(F)(F)F)o1 | 0 |
| KKL-0713 | Oc1cccc(-n2c(/C=C/c3ccc([N+]([O-])=O)o3)nc3ccccc3c2=O)c1 | 0 |
| KKL-0714 | COc1ccc(NC(=O)CC23CC4CC(CC(C4)C2)C3)cc1[N+]([O-])=O | 5 |
| KKL-0715 | Fc1ccc(Nc2nc(-c3ccccn3)cs2)cc1 | 18 |
| KKL-0716 | COc1ccc(Cn2c3nncn3c3c(c2=O)C2(Cc4c-3cccc4)CCCCC2)cc1 | 13 |
| KKL-0717 | Cc1cc(OC(=O)C2CC2)c2c3c(c(=O)oc2c1)CCCC3 | 10 |
| KKL-0718 | CCCCc1cc(=O)oc2cc(OCC(=O)OCc3ccccc3)c(Cl)cc12 | 0 |
| KKL-0719 | FC(F)(F)c1cccc(NS(=O)(=O)c2ccc(NC(=O)Nc3ccccc3)cc2)c1 | 0 |
| KKL-0720 | [O-][N+](=O)c1ccc(NC(=O)CC2Sc3ccc(Cl)cc3NC2=O)cc1 | 15 |
| KKL-0721 | Cc1ccccc1OC(=O)c1sc2ccccc2c1Cl | 0 |
| KKL-0722 | Clc1ccc(SCC(=O)Nc2ccc(Br)cn2)cc1 | 8 |
| KKL-0723 | COc1ccccc1-c1nc2cccc3cccc([nH]1)c23 | 0 |
| KKL-0724 | CC1CCN(CCN=C(S)Nc2ccc(F)cc2)CC1 | 0 |
| KKL-0725 | OC(COc1ccccc1Cl)CN1CCN(CCN2C(=O)c3cccc4cccc(c34)C2=O)CC1 | 6 |

| | | |
|---|---|---|
| KKL-0726 | CCOC(=O)c1cccc(NC(=O)C(Cl)Cl)c1 | 0 |
| KKL-0727 | CCOC(=O)c1ccc(NC(=O)/C=C/c2ccc([N+]([O-])=O)s2)cc1 | 0 |
| KKL-0728 | Clc1cccc(Nc2nc(NCc3ccco3)c3ccccc3n2)c1 | 0 |
| KKL-0729 | Cc1ccc(-n2c(=O)oc3ccc(Cl)cc3c2=O)cc1Cl | 0 |
| KKL-0730 | CCOC(=O)c1ccc2nc(C)cc(Nc3ccc(N4CCCCC4)cc3)c2c1 | 0 |
| KKL-0731 | CC(C(=O)OC1CCCCC1)n1cnc2scc(-c3ccc(C)c(C)c3)c2c1=O | 0 |
| KKL-0732 | COC(=O)CSc1nnc(NC(=O)c2ccc(C(C)(C)C)cc2)s1 | 0 |
| KKL-0733 | CCn1nnc(NC(=O)COc2ccc(C3CCCCC3)cc2)n1 | 8 |
| KKL-0734 | COc1ccc(Cc2nn3c(-c4ccccc4)nnc3s2)cc1 | 15 |
| KKL-0735 | Cc1c(C(=O)NC(=S)Nc2ccc(Oc3ccc(Cl)cc3)cc2)cccc1[N+]([O-])=O | 17 |
| KKL-0736 | ClC(Cl)(Cl)C(=O)Nc1ccccc1-c1ccccc1 | 0 |
| KKL-0737 | Fc1ccc(CC(=O)Nc2ccc(Cl)c(C(F)(F)F)c2)cc1 | 14 |
| KKL-0738 | [O-][N+](=O)c1cc(C#N)c(C#N)cc1Oc1ccc(OCc2ccccc2)cc1 | 5 |
| KKL-0739 | CCc1ccc(Oc2coc3cc(OC(C)C(=O)OC)ccc3c2=O)cc1 | 3 |
| KKL-0740 | CC(C(=O)OCc1ccccc1)n1cnc2sc3c(c2c1=O)CCC(C)C3 | 7 |
| KKL-0741 | Clc1cc(-c2ccccc2)oc(=O)c1Cl | 0 |
| KKL-0742 | CCCOc1ccc(Nc2cc(C)nc3ccc(Cl)cc23)cc1 | 0 |
| KKL-0743 | CC(C)c1ccc(OCC(=O)NNC(=S)NC(=O)c2ccco2)c(Br)c1 | 24 |
| KKL-0744 | Fc1ccc(NC(=O)CC23CC4CC(CC(C4)C2)C3)cc1Cl | 0 |
| KKL-0745 | CCC(C)c1ccc(O)c(NC(=S)NC(=O)c2ccc(-c3ccc(Cl)cc3)o2)c1 | 0 |
| KKL-0746 | CC1CCN(C(=S)NC(=O)c2ccc(Cl)cc2)CC1 | 13 |
| KKL-0747 | Cc1nc2ccccc2n1CC(O)COc1ccc(C2CCC=C2)cc1 | 0 |
| KKL-0748 | COC(=O)COc1ccc2c(c1)oc(C)c(-c1ccccc1Cl)c2=O | 0 |
| KKL-0749 | CCOC(=O)COc1ccc2c(oc(C)c(-c3ccccc3Cl)c2=O)c1C | 2 |
| KKL-0750 | [O-][N+](=O)c1ccc(-c2[nH]c(-c3cccc(F)c3)nc2-c2ccccc2)cc1 | 0 |
| KKL-0751 | CN(C(C)=O)c1ccc(OCC(O)Cn2c3ccc(Cl)cc3c3cc(Cl)ccc23)cc1 | 7 |
| KKL-0752 | CCc1ccc(OCC(=O)Nc2ccc(N3CCN(C(=O)c4cccs4)CC3)c(Cl)c2)cc1 | 0 |
| KKL-0753 | COc1ccc(NC(=O)CC23CC4CC(CC(C4)C2)C3)cc1Cl | 3 |
| KKL-0754 | [O-][N+](=O)c1ccc(C(=O)Oc2c(Cl)cc(Cl)c3cccnc23)cc1 | 96 |

| | | |
|---|---|---|
| KKL-0755 | COc1cc(CNC23CC4CC(CC(C4)C2)C3)ccc1OCc1ccccc1 | 4 |
| KKL-0756 | O=C(Oc1cccc2cccnc12)c1ccco1 | 0 |
| KKL-0757 | COc1cc(NC(=O)C(Cl)Cl)c(OC)cc1Cl | 0 |
| KKL-0758 | CCC(OC(C)=O)c1ccc(OC(C)=O)c(OC)c1 | 7 |
| KKL-0759 | COC(=O)C(C)Sc1nnc(-c2ccc(Br)cc2)o1 | 0 |
| KKL-0760 | CCc1nc2ccccc2n1CC(O)COc1ccc(I)cc1 | 0 |
| KKL-0761 | CCCOC(=O)c1ccc(NC(=S)NC(=O)c2ccc(Cl)cc2)cc1 | 3 |
| KKL-0762 | COC(=O)COc1ccc2c(-c3cccc([N+]([O-])=O)c3)cc(=O)oc2c1 | 0 |
| KKL-0763 | CCOC(=O)C(C)Oc1ccc2c(-c3ccc(OC)cc3)cc(=O)oc2c1 | 0 |
| KKL-0764 | CN(C)c1ccc(CNC23CC4CC(CC(C4)C2)C3)cc1 | 0 |
| KKL-0765 | Oc1nc(/C=C/c2cccc3ccccc23)nc(O)c1[N+]([O-])=O | 0 |
| KKL-0766 | NC(CC(=O)c1ccc(Cl)cc1Cl)C(Cl)(Cl)Cl | 0 |
| KKL-0767 | Cc1nn(-c2ccccc2)c(S)c1C=Nc1ccccc1O | 0 |
| KKL-0768 | C=CCN1C(=N)S/C(=C2\C(=O)Nc3ccccc32)C1=O | 1 |
| KKL-0769 | Cc1cc(C)c(N2C(=O)NC(=O)/C(=C\c3ccc[nH]3)C2=O)c(C)c1 | 0 |
| KKL-0770 | CN(C)c1ccc(Nc2nc(-c3ccccn3)cs2)cc1 | 0 |
| KKL-0771 | Cc1ccc2[nH]c3c(c2c1)CCCC3[NH2+]CCCOc1ccc(F)cc1 | 84 |
| KKL-0772 | CCOC(=O)COc1ccc2cc(-c3nc4ccccc4s3)c(=O)oc2c1 | 2 |
| KKL-0773 | Cc1nc2cc(C)ccn2c1-c1csc(Nc2ccccn2)n1 | 0 |
| KKL-0774 | CCOC(=O)COc1ccc(S(=O)(=O)Nc2cccc(Cl)c2C)cc1 | 3 |
| KKL-0775 | CN(C)CCC(=O)c1ccc(Br)cc1 | 0 |
| KKL-0776 | CC(C)C(=O)Nc1ccc(C(=O)Oc2ccccc2)c(O)c1 | 0 |
| KKL-0777 | CCCCOC(=O)COc1ccc2c(c1)oc(C)c(Oc1ccc(OCC)cc1)c2=O | 11 |
| KKL-0778 | Cc1ccc(-c2ccc(/C=C/C(=O)c3ccco3)o2)cc1 | 0 |
| KKL-0779 | Cc1ccc(S(=O)(=O)NC(=O)Oc2cccc3cccnc23)cc1 | 13 |
| KKL-0780 | CC(NCC(O)COc1ccccc1C(C)(C)C)c1ccccc1 | 17 |
| KKL-0781 | CC(=O)Nc1nc2ccccc2n1CCOc1ccccc1 | 1 |
| KKL-0782 | Cc1ccc(-c2csc(-c3ccc(O)cc3)n2)cc1 | 14 |
| KKL-0783 | CCOC(=O)COc1ccc(C(=O)Nc2ccc(F)cc2)cc1 | 0 |

| | | |
|---|---|---|
| KKL-0784 | Cc1cnc(NC(=O)CSc2ccc(Cl)cc2)s1 | 0 |
| KKL-0785 | CC1(C)Cc2nc3ccccc3c(NCc3ccccc3)c2C=C1 | 0 |
| KKL-0786 | CC1(C)Oc2ccc3ccc(=O)oc3c2C(C=O)=C1Cl | 0 |
| KKL-0787 | COc1ccc(OCC(=O)Nc2cccc(Cl)c2)cc1 | 0 |
| KKL-0788 | CCOC(=O)/C(=C/c1ccc(-c2ccc(F)cc2)o1)C#N | 0 |
| KKL-0789 | COC(=O)COc1ccc2c(c1)occ(-c1ccc(Cl)cc1)c2=O | 0 |
| KKL-0790 | OC(COCCC12CC3CC(CC(C3)C1)C2)CN1CCc2ccccc2C1 | 0 |
| KKL-0791 | O=C(Nc1nc(-c2ccccn2)cs1)C1CCCC1 | 0 |
| KKL-0792 | C(CNc1nc(-c2ccccc2)nc2ccccc12)Cn1ccnc1 | 0 |
| KKL-0793 | CCc1ccc(C=c2sc(=CC(=O)C(C)(C)C)[nH]c2=O)cc1 | 0 |
| KKL-0794 | OC(CNCCc1ccc(C(F)(F)F)cc1)COc1cccc2ccccc12 | 0 |
| KKL-0795 | Cc1cc(NC(=O)C2CCCCC2)n(-c2nc3ccccc3[nH]2)n1 | 9 |
| KKL-0796 | CCc1cccc2c(C(=O)COC(=O)c3ccc([N+]([O-])=O)s3)c[nH]c12 | 40 |
| KKL-0797 | C(N1CCC(Nc2nc(-c3cccs3)nc3ccccc23)CC1)c1ccccc1 | 0 |
| KKL-0798 | COCCCNC1=NN=C(c2cc(C)n(-c3ccccc3F)c2C)CS1 | 0 |
| KKL-0799 | [O-][N+](=O)c1ccc(Cl)cc1C(=O)Nc1nc(-c2ccccn2)cs1 | 16 |
| KKL-0800 | O=C(CCCSc1ccccc1)Nc1nc(-c2ccccn2)cs1 | 1 |
| KKL-0801 | CN(C)CCCn1c(SCC(=O)Nc2nc(-c3ccc(C)cc3)cs2)nc2ccccc2c1=O | 0 |
| KKL-0802 | Cc1sc2nc(-c3ccccn3)nc(O)c2c1-c1ccccc1 | 26 |
| KKL-0803 | Nc1ccccc1Nc1ccccc1 | 4 |
| KKL-0804 | CCCCNC(=O)Nc1nnc(-c2ccc(Cl)cc2)o1 | 1 |
| KKL-0805 | Cc1nn(Cc2ccccc2Cl)c2sc(C(=O)NCc3ccc4c(c3)OCO4)cc12 | 0 |
| KKL-0806 | CN(C)CCCn1c(SCC(=O)Nc2nc(-c3cccc(Br)c3)cs2)nc2ccccc2c1=O | 0 |
| KKL-0807 | COc1ccc(-c2nnc(SCC(=O)Nc3cc(C)on3)o2)cc1 | 0 |
| KKL-0808 | CC(C)(c1ccccc1)c1ccc(OCC(O)CNC2CCN(Cc3ccccc3)CC2)cc1 | 0 |
| KKL-0809 | CCSc1nnc(NC(=O)c2ccc3ccccc3n2)s1 | 5 |
| KKL-0810 | CCOc1ccc2nc(NC(=O)c3ccccn3)sc2c1 | 0 |
| KKL-0811 | CCC(CC)c1nnc(NC(=O)c2ccc3ccccc3n2)s1 | 0 |
| KKL-0812 | Fc1ccc(CSc2nnc(NC(=O)c3ccccn3)s2)cc1 | 14 |

| | | |
|---|---|---|
| KKL-0813 | Fc1ccc2nc(NC(=O)c3ccccn3)sc2c1 | 0 |
| KKL-0814 | O=C(Nc1nc(-c2ccccn2)cs1)/C=C/c1ccco1 | 0 |
| KKL-0815 | [O-][N+](=O)c1ccc(C(=O)OCC(=O)c2c[nH]c3ccccc23)s1 | 0 |
| KKL-0816 | CC1CCCCN1C(=O)COC(=O)/C=C/c1ccc2ccccc2n1 | 0 |
| KKL-0817 | Oc1nc(-c2ccccn2)nc2sc(-c3ccccc3)cc12 | 20 |
| KKL-0818 | NC(=O)c1ccc(NC(=O)COC(=O)c2c3c(nc4ccccc24)/C(=C\c2cccs2)CCC3)cc1 | 0 |
| KKL-0819 | CC1CCN(C(=O)COC(=O)c2cc3ccccc3cc2O)CC1 | 0 |
| KKL-0820 | COc1cc(/C=C/C)ccc1OCC(=O)OCc1csc(Nc2ccc(C)cc2)n1 | 0 |
| KKL-0821 | CCOC(=O)C1=C(NC(=O)Nc2ccc(Cl)c(Cl)c2)Nc2ccccc2N=C1CC | 0 |
| KKL-0822 | CC1CCCC(NCC(O)COC(c2ccc(F)cc2)c2ccc(F)cc2)C1C | 0 |
| KKL-0823 | CCC1CCCCN1C(=O)COC(=O)c1cc2ccccc2cc1O | 0 |
| KKL-0824 | Cn1c(-c2ccccc2)c(C(=O)CCl)c2ccccc12 | 0 |
| KKL-0825 | c1cnc2c(c1)ccc1ccc[nH+]c21 | 8 |
| KKL-0826 | Cc1ccc(Nc2nnc(SCC(=O)Nc3cccnc3Cl)s2)cc1C | 0 |
| KKL-0827 | Cc1ccc(C)c(SCC(=O)OC/C(O)=C(/C#N)c2nc3ccccc3[nH]2)c1 | 4 |
| KKL-0828 | CCOC(=O)c1sc2nc(CSc3ccccc3)nc(N3CCCN(C)CC3)c2c1C | 0 |
| KKL-0829 | CCCn1c(NC(=O)c2cccc(C)c2C)nc2ccccc12 | 3 |
| KKL-0830 | [O-][N+](=O)c1ccccc1/C=C1\COc2ccccc2C1=O | 0 |
| KKL-0831 | CCn1cc(C(O)=O)c(=O)c2cc(F)c(N3CCNCC3)cc12 | 56 |
| KKL-0832 | CN(Cc1nc(O)c2sccc2n1)Cc1nc(O)c2ccccc2n1 | 0 |
| KKL-0833 | COc1ccc(Nc2ccccc2N)cc1 | 0 |
| KKL-0834 | Cc1ccc(Nc2ccccc2N)cc1 | 0 |
| KKL-0835 | Cc1cc(Nc2ccccc2)n(-c2nc3ccccc3[nH]2)n1 | 21 |
| KKL-0836 | CCN(Cc1ccccc1)c1ccc(C=NN=C(N)N)cc1 | 18 |
| KKL-0837 | Clc1ccc2oc(SCC(=O)NCCc3ccccc3)nc2c1 | 0 |
| KKL-0838 | CSCC[C@H](NC(=O)c1cc([N+]([O-])=O)cc([N+]([O-])=O)c1)c1nc2ccccc2[nH]1 | 0 |
| KKL-0839 | CCOC(=O)c1sc(NC(=O)CNCc2ccccn2)c(C(=O)OCC)c1C | 0 |
| KKL-0840 | Cc1nn(-c2ccccn2)c2c1C(c1c(C)cc(C)cc1C)C1=C(O)c3ccccc3C1=N2 | 0 |
| KKL-0841 | O=C(NCCc1c[nH]c2ccccc12)c1cc2ccccc2oc1=O | 0 |

| | | |
|---|---|---|
| KKL-0842 | CC1CC(C)CN(C(=O)COC(=O)c2cc3ccccc3cc2O)C1 | 0 |
| KKL-0843 | O=C(Nc1ccc2c(c1)OCCO2)c1cccc2nc3ccccc3nc12 | 0 |
| KKL-0844 | Fc1c(F)c(F)c(-c2cc(=O)c3ccccc3o2)c(F)c1F | 0 |
| KKL-0845 | CC(C)C1CCC(C)CC1OC(=O)Cn1c(COc2ccccc2C)[n+](C)c2ccccc12 | 0 |
| KKL-0846 | COc1ccc(N=c2cc(-c3ccc(O)cc3)oc3ccccc23)cc1 | 0 |
| KKL-0847 | CCC(CO)N1C(C)=CC(=Cc2sc3ccccc3[n+]2Cc2ccccc2)C=C1C | 0 |
| KKL-0848 | FC1(F)C(c2ccccc2)N(c2ccc3c(c2)OCCO3)C1=O | 0 |
| KKL-0849 | [O-][N+](=O)c1cccnc1Sc1nnc(NCC=C)s1 | 0 |
| KKL-0850 | COC(=O)CN1C(c2ccccc2)c2cc(Cl)ccc2N=C1c1ccc(OC)cc1 | 0 |
| KKL-0851 | O=C1Nc2ccccc2C1=NN=CC1=C(N2CCOCC2)/C(=C/c2ccccc2)CC1 | 0 |
| KKL-0852 | CC(C)c1ccc(Nc2nnc(SCC(=O)Nc3ncc(S(=O)(=O)c4ccc([N+]([O-])=O)cc4)s3)s2)cc1 | 1 |
| KKL-0853 | Cc1cccc(Nc2cc(Cl)c3nonc3c2[N+]([O-])=O)c1C | 0 |
| KKL-0854 | CCOC(=O)C1=C(C)N(c2ccc(Br)cc2)C(=O)C1 | 0 |
| KKL-0855 | CCOC(=O)COc1cc2oc(=O)cc(CC)c2cc1Cl | 0 |
| KKL-0856 | Cc1ccc2oc(-c3cc(NC(=O)Cc4ccccc4)ccc3Cl)nc2c1 | 0 |
| KKL-0857 | CC1CCCc2sc(NC(=S)NC(=O)c3ccccc3)c(C#N)c21 | 0 |
| KKL-0858 | Cc1ccc(C(=O)NCCC23CC4CC(CC(C4)C2)C3)cc1 | 0 |
| KKL-0859 | CC(=O)c1cccc(Nc2cc(C)nc3c2ccc2ccccc32)c1 | 0 |
| KKL-0860 | O=C(COc1ccc2c(-c3ccccc3)cc(=O)oc2c1)OC1CCCCC1 | 0 |
| KKL-0861 | Cc1cc(Sc2ncnc(N)c2[N+]([O-])=O)ncn1 | 0 |
| KKL-0862 | CC(C)(C)c1ccc(OCCN2C(=O)C(=O)c3cc(Br)ccc32)cc1 | 0 |
| KKL-0863 | Clc1ccc(OCC(=O)Nc2ccc(N3CCN(C(=O)c4ccco4)CC3)c(Cl)c2)cc1 | 0 |
| KKL-0864 | COc1ccc(-c2csc(-c3ccc(OC)c(OC)c3)n2)cc1C | 1 |
| KKL-0865 | FC(F)(F)c1cc(NC(=O)CSc2ccc(Cl)cc2)ccc1Cl | 0 |
| KKL-0866 | Clc1cccc(NC(=O)c2cc(Br)ccc2Cl)c1 | 0 |
| KKL-0867 | CC(C)(C)c1ccc(C(=O)Nc2cccc(NC(=O)c3cccs3)c2)cc1 | 0 |
| KKL-0868 | CC1CCc2sc3ncnc(NCCc4ccccc4)c3c2C1 | 2 |
| KKL-0869 | OCCN1CCN(c2ccc([N+]([O-])=O)c(NCCOc3cccc4ccccc34)c2)CC1 | 0 |
| KKL-0870 | CCn1nnc(NC(=O)c2sc3ccccc3c2Cl)n1 | 0 |

| | | |
|---|---|---|
| KKL-0871 | COc1ccc(CCNc2cc(C)nc3c(-c4ccccc4)c(C)nn23)cc1OC | 0 |
| KKL-0872 | CC(C)(C)c1cc(NCCCn2ccnc2)n2ncc(-c3ccccc3)c2n1 | 0 |
| KKL-0873 | Clc1ccc(C(=O)NCc2nnc3c4ccccc4c(-c4ccccc4)nn23)cc1 | 0 |
| KKL-0874 | Oc1ccccc1N=Cc1ccc[nH]1 | 4 |
| KKL-0875 | [O-][n+]1onc2c1ccc1nnc(-c3ccccc3)cc21 | 8 |
| KKL-0876 | CCn1cc(C(=O)NCc2cccc(F)c2)c(=O)c2cc(F)c(N3CCN(C(=O)c4ccco4)CC3)cc12 | 8 |
| KKL-0877 | C=CCOC(=O)C(C#N)c1nc2ccccc2nc1N1CCN(Cc2ccc3c(c2)OCO3)CC1 | 0 |
| KKL-0878 | Oc1nc(-c2ccccn2)nc2ccccc12 | 0 |
| KKL-0879 | CCOC(=O)c1cnc(-n2nc(C)cc2C)nc1Nc1ccc(OC)cc1 | 0 |
| KKL-0880 | COc1ccc(C(CCNCc2ccc(F)cc2)c2ccc(F)cc2)cc1 | 0 |
| KKL-0881 | CN1CCN(c2cc(C(C)(C)C)nc3c(-c4ccccc4)c(C)nn23)CC1 | 0 |
| KKL-0882 | CC(C)c1cc(Nc2ccc(NC(C)=O)cc2)n2nc(C)c(-c3ccccc3)c2n1 | 0 |
| KKL-0883 | Cc1nn2c(O)c(Cc3ccccc3)c(C)nc2c1-c1ccc(Cl)cc1 | 0 |
| KKL-0884 | CCCCCCCCCCn1c2c(c(=N)c3c1CCC3)CCC2 | 12 |
| KKL-0885 | COC(=O)CSc1nc2c(c(-c3ccco3)c1C#N)CCCC2 | 8 |
| KKL-0886 | CC(=O)/C=C/c1ccc2ccccc2n1 | 0 |
| KKL-0887 | N=c1n(Cc2ccccc2)cnc2n(Cc3ccco3)c(-c3ccccc3)c(-c3ccccc3)c12 | 0 |
| KKL-0888 | Fc1ccc(COc2cccc(-c3nn(-c4ccccc4)cc3/C=C(/C#N)C(=O)NCCCn3ccnc3)c2)cc1 | 0 |
| KKL-0889 | CCOC(=O)c1cnc(-n2nc(C)cc2C)nc1Nc1ccc(N(C)C)cc1 | 0 |
| KKL-0890 | CCC(C)NC(=O)c1c(N)n(N=Cc2ccccn2)c2nc3ccccc3nc12 | 0 |
| KKL-0891 | Cc1nn2c(O)cc(C(C)(C)C)nc2c1-c1ccccc1 | 17 |
| KKL-0892 | CCCc1cc(NCc2ccccn2)n2nc(C)c(-c3ccccc3)c2n1 | 0 |
| KKL-0893 | CCc1nc2sc3ccccc3c(=O)c2c(=O)n1Cc1ccc(OC)cc1 | 0 |
| KKL-0894 | CCCCCCCCCn1c2c(c(=N)c3c1CCCC3)CCC2 | 7 |
| KKL-0895 | COC(=O)CSc1nc2c(c(-c3ccc(C)o3)c1C#N)CCCC2 | 0 |
| KKL-0896 | Oc1cc2[nH]c3ccccc3c2cc1C(=O)Nc1ccc(Cl)cc1 | 102 |
| KKL-0897 | CN1C(=O)N(C)C(=O)C(=C(Nc2ccccc2)SCc2ccccc2Cl)C1=O | 0 |
| KKL-0898 | C(Nc1nc(-c2ccccn2)nc2ccccc12)C1CCCO1 | 0 |
| KKL-0899 | CCOC(=O)c1cnc(-n2nc(C)cc2C)nc1NC1CCCC1 | 4 |

| | | |
|---|---|---|
| KKL-0900 | COc1ccccc1N1CCN(c2nc(C)nc3c4ccccc4oc23)CC1 | 4 |
| KKL-0901 | COc1ccc(CCNc2cc(C)nc3c(-c4ccc(Cl)cc4)c(C)nn23)cc1OC | 0 |
| KKL-0902 | CCCc1cc(NCCCn2ccnc2)n2nc(C)c(-c3ccccc3)c2n1 | 17 |
| KKL-0903 | CCN1CCN(c2cc3n(C4CC4)cc(C(O)=O)c(=O)c3cc2F)CC1 | 6 |
| KKL-0904 | CN1CCN(c2ccc(C(=O)Nc3ccc(C)c(Cl)c3)cc2[N+]([O-])=O)CC1 | 2 |
| KKL-0905 | CC(C)(C)c1cc(Cl)c(O)c(NC(=S)NC(=O)c2cccc3ccccc23)c1 | 0 |
| KKL-0906 | C(Cc1ccccc1)N(Cc1ccccc1)C1CCN(CC2CC3CC2C=C3)CC1 | 0 |
| KKL-0907 | C(Cc1ccccc1)N(Cc1ccccc1)C1CCN(Cc2c[nH]c3ccccc23)CC1 | 0 |
| KKL-0908 | COc1ccccc1OC(=O)c1sc2ccccc2c1Cl | 0 |
| KKL-0909 | CCc1cc2c(cc1OCC(=O)OC)occ(-c1ccc(OC)cc1)c2=O | 0 |
| KKL-0910 | CCCCOc1ccc(S(=O)(=O)Nc2ccc(I)cc2)cc1 | 10 |
| KKL-0911 | CCCCCn1c(NCc2ccc(N(C)C)cc2)nc2ccccc12 | 0 |
| KKL-0912 | CCCC(=O)NC(=S)Nc1ccc(N2CCN(C(=O)c3cccs3)CC2)c(Cl)c1 | 0 |
| KKL-0913 | COc1ccccc1-c1coc2cc(OCC(=O)OCc3ccccc3)ccc2c1=O | 0 |
| KKL-0914 | Cc1cccc(N2CCN(C3=NC(=O)C(=C4CCCCC4)S3)CC2)c1 | 0 |
| KKL-0915 | CCC1CCCCN1C(=S)NC(=O)c1ccc(F)cc1 | 0 |
| KKL-0916 | [O-][N+](=O)c1ccc(Oc2ccc(C(=O)Oc3ccc(F)cc3)cc2)cc1 | 0 |
| KKL-0917 | CCc1ccccc1-n1c(=O)oc2ccc(Cl)cc2c1=O | 1 |
| KKL-0918 | CC(C)OC(=O)c1ccc(NC(=O)COc2c(C)ccc(C)c2C)cc1 | 0 |
| KKL-0919 | Brc1ccc(C(=O)NCCC23CC4CC(CC(C4)C2)C3)cc1 | 0 |
| KKL-0920 | OC(COc1ccccc1Cl)CN1CCN(C(c2ccccc2)c2ccccc2)CC1 | 0 |
| KKL-0921 | COc1ccc(Nc2nc(-c3ccccn3)cs2)c(OC)c1 | 0 |
| KKL-0922 | CSc1nc2ccc(NC(=O)C3CN(Cc4ccccc4)C(=O)C3)cc2s1 | 0 |
| KKL-0923 | COc1ccc(-c2nc(NC(=O)c3cnc(SC)nc3)sc2C)cc1F | 0 |
| KKL-0924 | Fc1cc(F)c2nc(NC(=O)c3ccccn3)sc2c1 | 0 |
| KKL-0925 | OC(CN1CCC2(CC1)CC(O)c1cc(F)ccc1O2)c1ccc2c(c1)CCCC2 | 0 |
| KKL-0926 | COc1ccc2[nH]c(CN3CCC(O)(c4cccc(C)c4)CC3)c(C)c2c1 | 5 |
| KKL-0927 | CSc1ncc(CN2CCC(O)(c3ccc(C(F)(F)F)cc3)CC2)cn1 | 0 |
| KKL-0928 | COc1ccc(-c2csc(NC(=O)C3CC4C=CC3C43CC3)n2)cc1OC | 0 |

| | | |
|---|---|---|
| KKL-0929 | COCc1nnc(NC(=O)c2nnn(-c3ccc(F)cc3)c2C)s1 | 7 |
| KKL-0930 | COc1ccc(-c2cc(C(=O)N3C[C@H](C)N(Cc4ccco4)C[C@@H]3C)no2)cc1 | 0 |
| KKL-0931 | CSc1ncc(C(=O)Nc2ccc(Oc3ccc4c(c3)OCO4)cc2)cn1 | 7 |
| KKL-0932 | CCC(NC(=O)c1cnc(C(C)(C)C)nc1)c1ccc(F)cc1 | 5 |
| KKL-0933 | OC(CN1CCC(O)(c2ccc(F)cc2)CC1)c1ccc(-c2ccccc2)cc1 | 0 |
| KKL-0934 | OC(CN1CCC2(CC1)CC(O)c1cc(Cl)ccc1O2)c1ccc(Cl)cc1 | 0 |
| KKL-0935 | CC(C)(C)c1ccc(C(O)CN2CCC3(CC2)CC(O)c2cc(F)ccc2O3)cc1 | 0 |
| KKL-0936 | Cc1cc2c(cc1Cl)C(O)CC1(O2)CCN(CC(O)c2ccc(Cl)cc2)CC1 | 2 |
| KKL-0937 | Cc1c(CN2CCC(O)(c3ccc(F)cc3)CC2)[nH]c2ccc(F)cc12 | 0 |
| KKL-0938 | Cc1oc(C(=O)Nc2ncc(Cc3ccccc3F)s2)cc1CN1CCOCC1 | 0 |
| KKL-0939 | COc1cc(/C=C/c2ccc3c([N+]([O-])=O)cccc3n2)ccc1OCC(O)=O | 0 |
| KKL-0940 | Fc1ccccc1Cc1cnc(NC(=O)c2ccccc2Cn2ccnc2)s1 | 0 |
| KKL-0941 | Clc1cccc(Cc2cnc(NC(=O)CCCn3ccnc3)s2)c1Cl | 0 |
| KKL-0942 | [O-][N+](=O)c1cc(/C=C/C(=O)c2cccnc2)ccc1Cl | 0 |
| KKL-0943 | [O-][N+](=O)c1cc(C(=O)Oc2ccc(Cl)cc2)ccc1N1CCOCC1 | 0 |
| KKL-0944 | COc1cccc(C(=O)Oc2ccccc2/C=C/C(=O)c2cc3ccccc3o2)c1 | 0 |
| KKL-0945 | O=C(Nc1nnc(C2CC2)s1)c1ccccn1 | 0 |
| KKL-0946 | Cc1nnc(NC(=O)c2ccc3ccccc3c2)o1 | 0 |
| KKL-0947 | Cc1cc(NCc2ccccn2)n2ncc(-c3ccccc3)c2n1 | 0 |
| KKL-0948 | CCSc1nnc(NC(=O)c2ccccn2)s1 | 0 |
| KKL-0949 | CC(C)(C)c1ccc(-c2c[n+](-c3ccc(Br)cc3)c3n2CCCS3)cc1 | 0 |
| KKL-0950 | COc1ccc(C=c2sc(=CC(=O)C(C)(C)C)[nH]c2=O)cc1O | 0 |
| KKL-0951 | Cc1ccc(Nc2cc(C)nc(NCc3ccccc3)n2)cc1 | 4 |
| KKL-0952 | Cc1c(C(=O)Oc2cccc3cccnc23)cccc1[N+]([O-])=O | 8 |
| KKL-0953 | Cc1nn(-c2ccccc2)c(O)c1N=Nc1nccs1 | 27 |
| KKL-0954 | OCCN1CCN(c2nc(-c3ccccc3)nc3c4ccccc4oc23)CC1 | 12 |
| KKL-0955 | CCOC(=O)CSc1nc(C#N)c(SCC(=O)OCC)s1 | 12 |
| KKL-0956 | Cc1ccc(-c2csc(-c3ccncc3)n2)cc1C | 0 |
| KKL-0957 | CC(=O)Nc1ccc2cccc(O)c2n1 | 55 |

| | | |
|---|---|---|
| KKL-0958 | CCOC(=O)COc1ccc2c(c1)oc(C(=O)OCC)c(-c1ccc(OC)c(OC)c1)c2=O | 16 |
| KKL-0959 | CCOC(=O)c1ccc(NC(=O)COc2ccccc2CNCCc2ccccc2)cc1 | 0 |
| KKL-0960 | [O-][N+](=O)c1cc(S(=O)(=O)C(F)(F)F)ccc1Sc1nc2ccccc2s1 | 0 |
| KKL-0961 | Cc1nc2c(cc1C(=O)/C=C/c1ccc(Br)cc1)CCCC2 | 0 |
| KKL-0962 | FC(F)(F)C1CC(c2ccc(Br)cc2)Nc2c(C(=O)NCc3ccco3)cnn21 | 14 |
| KKL-0963 | Cc1ccc(S/C=C\C(=O)c2ccccc2)cc1 | 0 |
| KKL-0964 | Cc1c(CNC(=O)c2cnn3c2NC(c2ccc(Br)cc2)CC3C(F)(F)F)cnn1C | 10 |
| KKL-0965 | CCCCC(=O)NC(=S)Nc1ccc(N2CCN(C(=O)c3ccco3)CC2)c(Cl)c1 | 8 |
| KKL-0966 | FC(F)(F)C1CC(c2ccc3c(c2)OCO3)Nc2cc(C(=O)NCc3cccs3)nn21 | 22 |
| KKL-0967 | CC(=O)c1ccc(S(=O)(=O)c2ccc(C)cc2)c([N+]([O-])=O)c1 | 8 |
| KKL-0968 | COC(=O)COc1cccc(NC(=O)COc2ccccc2)c1 | 0 |
| KKL-0969 | CN(C)c1ccc(/C=C/C=C2/C(=O)NC(=O)N(c3ccccc3)C2=O)cc1 | 0 |
| KKL-0970 | [O-][N+](=O)c1ccc(-c2nnc(SCC#CCOC(=O)c3cccs3)o2)cc1 | 6 |
| KKL-0971 | Cc1ccc(C)c(N=Nc2nc3ccccc3n2C)c1 | 0 |
| KKL-0972 | CCn1c(/C=C/c2ccc([N+]([O-])=O)o2)nc2ccccc2c1=O | 2 |
| KKL-0973 | Cc1ccc(C(=O)/C=C/c2ccccc2[N+]([O-])=O)cc1 | 0 |
| KKL-0974 | CC1CCCN(CCC(=O)c2ccc(Br)cc2)C1 | 7 |
| KKL-0975 | COc1cccc(OCC(=O)c2ccc(Br)cc2)c1 | 0 |
| KKL-0976 | CCOC(=O)c1c(C)c(C(N)=O)sc1NC(=O)c1c(F)c(F)c(F)c(F)c1F | 5 |
| KKL-0977 | Brc1ccc(C(=O)N/C(=C/c2ccccc2)C(=O)NCC(=O)OCc2ccccc2)cc1 | 0 |
| KKL-0978 | CCCN=C1C(=O)c2ccccc2C(O)=C1N(C(C)=O)c1cccc(F)c1 | 0 |
| KKL-0979 | CC(C)(C)/C(O)=C\C=Nc1ccccc1O | 0 |
| KKL-0980 | CC(=O)Nc1cccc(C(=O)/C=C/c2ccc(Br)cc2)c1 | 0 |
| KKL-0981 | Clc1ccc(OC(=O)C23CC4CC(CC(C4)C2)C3)c2ncccc12 | 8 |
| KKL-0982 | Cc1ccc(/C=C/C(=O)C2C(=O)c3ccccc3C2=O)cc1 | 0 |
| KKL-0983 | COC(=O)C(CC(C)C)NC(=O)/C=C/c1ccc(Cl)cc1 | 6 |
| KKL-0984 | CN1Cc2c(c3c(N)c(C(=O)c4ccc(Br)cc4)sc3nc2N2CCOCC2)CC1(C)C | 0 |
| KKL-0985 | CCCCCOc1ccc(C(=O)NCC(=O)NCC(=O)OCC)cc1 | 7 |
| KKL-0986 | CCCCCCn1c2ccccc2c(O)c(C(=O)Nc2nc(CC(=O)OCC)cs2)c1=O | 0 |

| | | |
|---|---|---|
| KKL-0987 | CCOC(=O)C1=Nc2ccccc2C1=[N+]=[N-] | 0 |
| KKL-0988 | CN1C(=O)N(C)C(=O)C(=C(Nc2ccccc2)SCc2ccc(Cl)cc2)C1=O | 10 |
| KKL-0989 | FC(F)(F)c1cccc(N2CCN(C3=C(NS(=O)(=O)c4cccs4)C(=O)c4ccccc4C3=O)CC2)c1 | 2 |
| KKL-0990 | CCOC(=O)c1cnc(-n2nc(C)cc2C)nc1NC(C)c1ccccc1 | 5 |
| KKL-0991 | OCc1nc2ccccc2n1N=Cc1ccc([N+]([O-])=O)s1 | 0 |
| KKL-0992 | Cc1nn2c(NCCCn3ccnc3)cc(C)nc2c1-c1ccc(Cl)cc1 | 4 |
| KKL-0993 | CN(C)CCCNc1cc(C(C)(C)C)nc2c(-c3ccc(Cl)cc3)cnn12 | 17 |
| KKL-0994 | COC(=O)CSc1nc(N)c2c3c(sc2n1)CCCC3 | 9 |
| KKL-0995 | ClCC(=O)N(c1c[nH]c2ccccc12)c1ccccc1 | 0 |
| KKL-0996 | Cc1nn(-c2ccccc2)c2c1C(c1ccc(O)c(O)c1)N1C(=N2)C(Nc2ccccc2)=Nc2c1[illegible] | 0 |
| KKL-0997 | O=Cc1ccc2oc3nc4n(c(=O)c3c(=O)c2c1)CCCS4 | 8 |
| KKL-0998 | CCOC(=O)c1cnc(-n2nc(C)cc2C)nc1NCCc1ccccc1 | 10 |
| KKL-0999 | C(CNc1nc(-c2ccccc2)nc2c3ccccc3oc12)Cn1ccnc1 | 6 |
| KKL-1000 | C1CCCN(c2nc(-c3ccccn3)nc3ccccc23)CC1 | 1 |
| KKL-1001 | CC(C)(C)c1cc(NCc2ccccn2)n2ncc(-c3ccc(Cl)cc3)c2n1 | 4 |
| KKL-1002 | Cc1ccc2nc(/N=C(\N)NC(=O)NC3CCCCC3)nc(C)c2c1 | 0 |
| KKL-1003 | CC(NC(=O)Nc1nnc(C(F)(F)F)s1)(C(F)(F)F)C(F)(F)F | 0 |
| KKL-1004 | CCCCCn1c2ccccc2c(O)c(C(=O)Nc2nc3ccccc3[nH]2)c1=O | 7 |
| KKL-1005 | O=C(Nc1nc(NC(=O)c2ccccc2)nn1-c1ccccc1)c1ccco1 | 44 |
| KKL-1006 | COc1cccc(C(=O)Nc2nc(-c3ccccc3)c(-c3ccccc3)o2)c1 | 9 |
| KKL-1007 | Clc1ccc2oc3nc4n(c(=O)c3c(=O)c2c1)CCCS4 | 8 |
| KKL-1008 | NC1CCN(c2c(F)cc3c(c2Cl)n(C2CC2)cc(C(O)=O)c3=O)C1 | 38 |
| KKL-1009 | N=c1c2c(ncn1Cc1ccco1)Oc1c(ccc3ccccc31)C2c1ccccc1 | 0 |
| KKL-1010 | CCN1CCN(c2cc(C(C)(C)C)nc3c(-c4ccccc4)c(C)nn23)CC1 | 3 |
| KKL-1011 | CC(C)(C)c1cc(NCCCn2ccnc2)n2ncc(-c3ccc(Cl)cc3)c2n1 | 9 |
| KKL-1012 | [O-][N+](=O)c1ccc(-c2nc(C(Cl)Cl)c3ccccc3n2)cc1 | 5 |
| KKL-1013 | CCCCCCc1cc2cc(C(=O)NC)c(=N)oc2cc1O | 12 |
| KKL-1014 | CCCCCCCn1c2ccccc2c(O)c(C(=O)Nc2cnccn2)c1=O | 0 |
| KKL-1015 | OC(COCC(F)(F)C(F)(F)C(F)(F)C(F)F)Cn1c2ccc(Cl)cc2c2cc(Cl)ccc12 | 0 |

| | | |
|---|---|---|
| KKL-1016 | N#Cc1nc(-c2cccs2)oc1NCCc1ccccc1 | 3 |
| KKL-1017 | COc1cc(OC)c(-c2nn3c(C4CCCCC4)nnc3s2)cc1OC | 1 |
| KKL-1018 | CN(C)CCSc1nc(-n2nc(C)cc2C)nc2sc3c(c12)CCCC3 | 7 |
| KKL-1019 | CCCCN1C(=O)NC(=O)/C(=C(O)\C=C\c2ccc3c(c2)OCO3)C1=O | 13 |
| KKL-1020 | COc1cccc(Nc2c3c(nc4c(-c5ccccc5)c(C)nn24)CCC3)c1 | 4 |
| KKL-1021 | Clc1ccc(NCc2n[n+](CC(=O)c3ccc(Br)cc3)c3n2CCCCC3)cc1 | 9 |
| KKL-1022 | CCCCNC(=O)c1cc2n(n1)C(C(F)(F)F)CC(c1ccccc1)N2 | 3 |
| KKL-1023 | CCc1nnc(NC(=O)c2cccc(C(=O)Nc3nnc(CC)s3)n2)s1 | 6 |
| KKL-1024 | CCCCCCCCCC(=NO)c1ccccn1 | 15 |
| KKL-1025 | COc1cc(OC)cc(C(=O)Nc2nc(-c3ccccc3)c(-c3ccccc3)o2)c1 | 15 |
| KKL-1026 | Cc1nc2ccccc2n1N=Cc1ccc([N+]([O-])=O)s1 | 0 |
| KKL-1027 | C(CC1=CCCCC1)Nc1[nH]cnc2c1nc1ccccc21 | 0 |
| KKL-1028 | C(Oc1ccccc1)c1nn2c(-c3[nH]nc4c3CCC4)nnc2s1 | 7 |
| KKL-1029 | OC(/C=C/c1ccc2c(c1)OCO2)=C1/C(=O)NC(=O)N(CCc2ccccc2)C<br>1=O | 0 |
| KKL-1030 | COc1ccc(N2CCN(Cc3nc(N)nc(Nc4ccc(C)c(Cl)c4)n3)CC2)cc1 | 7 |
| KKL-1031 | CN(C)C(CNC(=O)c1oc2c(C)cccc2c1C)c1ccc2c(c1)OCO2 | 0 |
| KKL-1032 | Cc1cc(C)c(-n2cccn2)c(CN(CCCn2ccnc2)Cc2ccc(F)cc2)c1 | 8 |
| KKL-1033 | CSc1nc2nc3c(c(O)n2n1)CCCC3 | 0 |
| KKL-1034 | CC1Cc2cc(CNCc3cn(C)c4cc(F)ccc34)ccc2O1 | 0 |
| KKL-1035 | COc1ccccc1CNCc1cc2c(C)nn(-c3ccc(F)cc3)c2nc1O | 8 |
| KKL-1036 | COc1ccc(CN(CCCn2ccnc2)Cc2[nH]c3ccc(F)cc3c2C)cc1 | 17 |
| KKL-1037 | CC(C)Oc1ccccc1CNCc1cc2c(C)nn(-c3ccccc3)c2nc1O | 0 |
| KKL-1038 | Cc1ccc2c(c1)C(=O)C(CN(CCCn1ccnc1)Cc1ccccc1Cl)=CO2 | 1 |
| KKL-1039 | Cc1ccc(CN(CCCn2ccnc2)Cc2ccc(N3CCCC3)c(F)c2)o1 | 0 |
| KKL-1040 | CCOc1ccccc1CNc1ccc(C2N(C(C)=O)CCc3sccc32)cc1 | 0 |
| KKL-1041 | COc1ccc2[nH]c(CN(CCCn3ccnc3)Cc3ccc(F)cc3)c(C)c2c1 | 16 |
| KKL-1042 | OC(CN1CCC(O)(c2ccc(C(F)(F)F)cc2)CC1)c1ccc(Cl)cc1 | 9 |
| KKL-1043 | CC12CCC(C(=O)Oc3cccc4cccnc34)(CC1=O)C2(C)C | 6 |
| KKL-1044 | CN(C)c1ccc(Nc2ncnc3ccc(Br)cc23)cc1 | 0 |

| | | |
|---|---|---|
| KKL-1045 | O=C(OCc1nnc(-c2ccccc2)o1)c1cc(-c2cccs2)nc2ccccc12 | 9 |
| KKL-1046 | CC1C(c2ccccc2)N(C)C(c2ccccc2)CC1(O)c1ccccc1 | 0 |
| KKL-1047 | O=c1oc2ccc3ccccc3c2cc1-c1nc2ccccc2o1 | 2 |
| KKL-1048 | Cc1ccc(-c2nnc3sc(CCc4ccccc4)nn23)cc1 | 0 |
| KKL-1049 | NS(=O)(=O)c1ccc(CCNc2ncnc3n(-c4ccc(F)cc4)cc(-c4ccccc4)c23)cc1 | 1 |
| KKL-1050 | Cc1ccc(C(=O)/C=C\C(=O)c2ccc(C)cc2)cc1 | 0 |
| KKL-1051 | C(Cc1ccccc1)Nc1ccnc2cc3ccccc3cc12 | 0 |
| KKL-1052 | COc1cc(CNC(C)c2ccccc2)cc(Cl)c1OCC(=O)Nc1ccccc1 | 0 |
| KKL-1053 | O=C(NC1CCCCC1)N1CCN(Cc2ccc3c(c2)OCO3)CC1 | 3 |
| KKL-1054 | COc1cc(NC(=O)c2ccc3c(c2)OCCO3)ccc1NC(=O)c1ccccc1Cl | 3 |
| KKL-1055 | Cc1cccc(Nc2nc(N)nc(CN3CCN(c4cccc(Cl)c4)CC3)n2)c1 | 0 |
| KKL-1056 | CN(C)c1ccc(N=Nc2nc3ccccc3n2C)cc1 | 0 |
| KKL-1057 | CCN(CC)c1ccc(/C=C(/C#N)c2nc3ccccc3[nH]2)c(OC(C)C)c1 | 0 |
| KKL-1058 | Cc1c(C(=O)Oc2cccc3cccnc23)oc2ccccc12 | 10 |
| KKL-1059 | CC(=O)C1=C(C)NC(SCC(=O)c2ccc(Cl)c(Cl)c2)=C(C#N)C1c1ccco1 | 2 |
| KKL-1060 | CC(=O)Nc1nc(NC(=O)/C=C/c2ccccc2)n(-c2ccccc2)n1 | 18 |
| KKL-1061 | CCOC(=O)CSCC(=O)Nc1c(C(=O)C2CC2)oc2ccccc12 | 5 |
| KKL-1062 | [O-][N+](=O)c1ccc(-c2nnc(SCC=C)o2)cc1 | 0 |
| KKL-1063 | Cc1cc(OCCCn2ccnc2)ccc1Cl | 5 |
| KKL-1064 | COC(=O)COc1ccc(CSCC(=O)Nc2ccc3ccccc3c2)cc1 | 6 |
| KKL-1065 | CCOC(=O)N1CCN(CCC(=O)c2ccc(Cl)cc2)CC1 | 0 |
| KKL-1066 | CCOc1cc(CNCc2ccccc2OCc2ccccc2)cc(Cl)c1OCC | 14 |
| KKL-1067 | [O-][N+](=O)c1ccc(-n2nnnc2-c2ccccc2)c([N+]([O-])=O)c1 | 0 |
| KKL-1068 | CCCOC(=O)c1ccc(NC(=O)c2ccc(Br)o2)cc1 | 0 |
| KKL-1069 | Oc1cccc2ccc(/C=C/c3cccs3)nc12 | 45 |
| KKL-1070 | Clc1ccc(OC(=O)c2ccc(COC(=O)c3cnccn3)cc2)cc1 | 3 |
| KKL-1071 | Cc1cc(C)n(-c2nc(C)cc(O)n2)n1 | 6 |
| KKL-1072 | CCc1ccc(/C=C/C(=O)C2C(=O)c3ccccc3C2=O)cc1 | 0 |
| KKL-1073 | CC(C)(C)C(=O)NC(=S)NNC(=O)C(c1ccccc1)c1ccccc1 | 16 |

| | | |
|---|---|---|
| KKL-1074 | O=C(CNCCN1CCOCC1)Nc1sc2c(c1C(=O)Nc1ccccc1)CCC2 | 2 |
| KKL-1075 | Fc1ccc(CCNC(=O)NC23CC4CC(CC(C4)C2)C3)cc1 | 0 |
| KKL-1076 | CCC(=Nc1ccccc1O)C1=C(O)CC(c2ccccc2)CC1=O | 1 |
| KKL-1077 | CCOC(=O)COc1ccc2c(c1)occ(-c1ccccc1OC)c2=O | 1 |
| KKL-1078 | CC1CCN(c2nc(-c3ccncc3)nc3ccccc23)CC1 | 1 |
| KKL-1079 | CC(C)=CCN1CCN(CCC2=C(C)CCCC2(C)C)CC1CCO | 4 |
| KKL-1080 | Cc1cccc(NC(=O)c2ccc([N+]([O-])=O)o2)c1 | 0 |
| KKL-1081 | CC(C)(C)c1ccc(C(=O)CC(=O)c2cc(Cl)ccc2O)cc1 | 3 |
| KKL-1082 | Clc1ccc(C(=O)Oc2cccc3cccnc23)cc1S(=O)(=O)N1CCOCC1 | 4 |
| KKL-1083 | CC(C)c1ccc(C)cc1OCCCCn1ccnc1 | 11 |
| KKL-1084 | Cc1cccc(C(=O)Nc2ccc(C(=O)/C=C/c3cccnc3)cc2)c1 | 5 |
| KKL-1085 | CCOC(=O)CSc1nc2c(c(-c3ccc(C)o3)c1C#N)CCCC2 | 0 |
| KKL-1086 | CCOC(=O)c1c(NC(=O)c2cc3ccccc3oc2=O)scc1-c1ccccc1 | 10 |
| KKL-1087 | [O-<br>][N+](=O)c1cc2c3c(cccc3c1N1CCOCC1)C(=O)N(c1cccc3ccccc13<br>[illegible] | 0 |
| KKL-1088 | COc1ccc(N2CCN(c3nc(NC(C)(C)C)nc(N4CCCCC4)n3)CC2)cc1 | 1 |
| KKL-1089 | Clc1ccccc1CN1C(=O)NC(=O)/C(=C\c2ccc[nH]2)C1=O | 5 |
| KKL-1090 | COc1cc(/C=C/C(=O)c2ccc(F)cc2)cc(OC)c1OC | 0 |
| KKL-1091 | Cc1ccc2[nH]c(NCc3ccccc3)nc2c1 | 0 |
| KKL-1092 | COc1ccc(Cc2nnc(SCC(=O)NCCc3ccccc3)o2)cc1 | 6 |
| KKL-1093 | COc1ccc(C(=O)CCN2CCCC(C)C2)cc1 | 9 |
| KKL-1094 | Fc1ccc(C(=O)CCCSc2nnc(SCCCC(=O)c3ccc(F)cc3)s2)cc1 | 2 |
| KKL-1095 | CCCOC(=O)CSc1nnc(-c2cc(OC)c(OC)c(OC)c2)o1 | 0 |
| KKL-1096 | C(Nc1nc(N2CCCCC2)nc2ccccc12)c1ccco1 | 0 |
| KKL-1097 | Cc1cc(C)cc(Nc2c([N+]([O-])=O)nn(-c3cccc(Cl)c3)[n+]2[O-])c1 | 4 |
| KKL-1098 | CCOc1ccc(CN2CCN(CC(O)Cn3c4ccccc4c4ccccc34)CC2)cc1 | 0 |
| KKL-1099 | Cc1n[nH]c(C)c1CCCCOc1cccc(C)c1C | 1 |
| KKL-1100 | Cc1ccc2[nH]c(/C(=C\c3ccc([N+]([O-])=O)o3)C#N)nc2c1 | 0 |
| KKL-1101 | CCOc1ccc(Nc2nc(-c3ccccn3)cs2)cc1 | 0 |
| KKL-1102 | COC(=O)c1ccc2nc(C)cc(Nc3ccc(OCc4ccccc4)cc3)c2c1 | 0 |

| | | |
|---|---|---|
| KKL-1103 | C(Cc1ccccc1)Nc1nc2ccccc2[nH]1 | 0 |
| KKL-1104 | [O-][N+](=O)c1ccc(Sc2nc3ccccc3o2)c([N+]([O-])=O)c1 | 0 |
| KKL-1105 | COC(=O)C(CC(C)C)NC(=O)/C=C/c1ccccc1Cl | 0 |
| KKL-1106 | OC(CNCc1ccco1)Cn1c2ccc(Br)cc2c2cc(Br)ccc12 | 0 |
| KKL-1107 | Cc1nn(-c2nc(-c3ccc(Cl)cc3)cs2)c(O)c1N=Nc1ccc(O)cc1 | 4 |
| KKL-1108 | Oc1cccc2ccc(/C=C/c3c(F)cccc3Cl)nc12 | 26 |
| KKL-1109 | CCOCCOC(=O)COc1ccc2c(-c3ccccc3)cc(=O)oc2c1 | 3 |
| KKL-1110 | CSc1nsc(SCC(=O)Nc2ccc(OC(C)=O)cc2)n1 | 0 |
| KKL-1111 | Cc1nc2cc(OCc3cc(C(=O)N4CCCC(c5ccccc5)C4)no3)ccc2s1 | 2 |
| KKL-1112 | O=C(/C=C/c1cccs1)N1CCc2ccc(NC(=O)c3ccccn3)cc2C1 | 0 |
| KKL-1113 | CCOC(=O)CCNC(=O)c1cc(COc2cc(C)c(Cl)c(C)c2)on1 | 0 |
| KKL-1114 | OC(=O)c1cn(-c2ccc(N3CCCC3)c(F)c2)c2c(F)c(F)c(F)cc2c1=O | 3 |
| KKL-1115 | Cc1ccc(Cl)cc1N=C1C=C(NS(=O)(=O)c2cccs2)c2ccccc2C1=O | 0 |
| KKL-1116 | Cc1nc([N+]([O-])=O)c(Br)n1Cc1ccc(Cl)c(Cl)c1 | 0 |
| KKL-1117 | Clc1ccc2nc(-c3ccccn3)c(Nc3ccc4c(c3)OCO4)n2c1 | 2 |
| KKL-1118 | COc1ccc2cc(-c3[nH]ncc3CNCc3ccc(C(F)(F)F)cc3)ccc2c1 | 0 |
| KKL-1119 | COCC1CCCN(c2nncc(-c3ccc(C4CCCCC4)cc3)n2)C1 | 0 |
| KKL-1120 | C1CCC(c2ccc(-c3cnnc(N4CCOCC4)n3)cc2)CC1 | 0 |
| KKL-1121 | COc1ccc2cc(C(=O)C3CCCN(Cc4ccc(C#CC(C)(C)O)cc4)C3)ccc2c1 | 10 |
| KKL-1122 | COC(=O)C(CCSC)NC(=O)c1cc(COc2ccc3sc(C)nc3c2)on1 | 0 |
| KKL-1123 | CCOC(=O)N1CCN(C(=O)CSc2nnc(-c3cc(OC)cc(OC)c3)o2)CC1 | 0 |
| KKL-1124 | Cc1cc(N2CCCCC2)nc2ccc(NC(=O)c3ccc(Br)o3)cc12 | 0 |
| KKL-1125 | Cc1ccc(C2CC(C(F)(F)F)n3nc(C(=O)NCc4ccco4)cc3N2)cc1 | 0 |
| KKL-1126 | Cc1nn2c(O)c3c(nc2c1-c1ccccc1)CCCC3 | 0 |
| KKL-1127 | COc1cccc(/C=C2\Sc3ccc(C(=O)NCc4cccnc4)cc3N(C)C2=O)c1 | 0 |
| KKL-1128 | Cc1ccccc1-n1ncc2c([N+]([O-])=O)cc([N+]([O-])=O)cc12 | 0 |
| KKL-1129 | COc1ccc(-n2cc(CNCCCn3ccnc3)c(-c3cccc(F)c3)n2)cc1 | 2 |
| KKL-1130 | Cc1cccc(-c2cccc(NC(=O)C3CCCN(Cc4ccc(C#CCO)s4)C3)c2)c1 | 4 |
| KKL-1131 | COc1cccc(-c2cc(CC3(O)CCN(Cc4ccc(SC)cc4)CC3)on2)c1 | 4 |

| | | |
|---|---|---|
| KKL-1132 | Cc1cc(C)n(CCNCc2cn(-c3ccc(C)cc3)nc2-c2ccc(F)cc2)n1 | 2 |
| KKL-1133 | Cc1ccc(C2CC(C(F)(F)F)n3ncc(C(=O)NCc4ccco4)c3N2)cc1 | 0 |
| KKL-1134 | COC(=O)COc1ccc2c(c1)occ(-c1ccccc1OC)c2=O | 3 |
| KKL-1135 | Clc1ccc(CCNC(=O)c2ccc(CNc3c(N4CCOCC4)c(=O)c3=O)cc2)cc1 | 0 |
| KKL-1136 | Cc1ccc(Nc2c(-c3ccccn3)nc3cccc(C)n23)cc1 | 0 |
| KKL-1137 | COc1cccc(OCCNC(=O)c2cc([N+]([O-])=O)cc([N+]([O-])=O)c2)c1 | 10 |
| KKL-1138 | COc1cccc(-c2ccccc2NC(=O)C2CCN(Cc3cc(C)ccc3C)CC2)c1 | 0 |
| KKL-1139 | Cc1cccc(-c2ccc(NC(=O)C3CCN(Cc4ccc(C#CCO)s4)CC3)cc2)c1 | 8 |
| KKL-1140 | CC(NC(=O)c1ccc2[n+]([O-])onc2c1)c1cnn(-c2ccccc2C)c1C | 0 |
| KKL-1141 | CSCCC(=O)N1CCC(C(=O)Nc2cccc(-c3cccc(Cl)c3)c2)CC1 | 0 |
| KKL-1142 | CCCC(=O)OCCNc1cc(SCc2ccco2)c2nonc2c1[N+]([O-])=O | 0 |
| KKL-1143 | Cc1[nH]c2ccccc2c1/C=C1\SC(=N)N(c2nccs2)C1=O | 6 |
| KKL-1144 | Cc1ccc(N=C2C=C(NS(=O)(=O)c3cccs3)c3ccccc3C2=O)c(C)c1 | 0 |
| KKL-1145 | COC(=O)[C@@H](NC(=O)c1nc(N2CCCC2)nc2c1CCCC2)c1ccccc1 | 0 |
| KKL-1146 | FC(F)(F)c1cccc(N2CCN(C(=O)C3CCN(c4nnc(-n5cccc5)s4)CC3)CC2)c1 | 5 |
| KKL-1147 | Cc1cccc(-c2cccc(NC(=O)C3CCCN(Cc4cccn4-c4ncccn4)C3)c2)c1 | 0 |
| KKL-1148 | CSCCC(=O)N1CCC(C(=O)Nc2ccc(-c3cccc(C)c3)cc2)CC1 | 1 |
| KKL-1149 | Cc1cc(C)n2nc(C(=O)N3CCc4c([nH]c5ccccc45)C3c3ccc(F)cc3)cc2n1 | 3 |
| KKL-1150 | COc1cccc(-c2ccccc2NC(=O)C2CCN(Cc3cc4ccccc4[nH]3)CC2)c1 | 2 |
| KKL-1151 | COc1cccc(-c2ccccc2NC(=O)C2CCN(Cc3ccc(OC)c(C)c3)CC2)c1 | 0 |
| KKL-1152 | CC(=O)OCCNc1cc(SCc2ccco2)c2nonc2c1[N+]([O-])=O | 0 |
| KKL-1153 | COc1ccc(CN2CCCCC2C(=O)Nc2ccc(-n3cccn3)cc2)c(F)c1 | 0 |
| KKL-1154 | Cc1ccccc1C1N(C(=O)c2cccc(C#CC(C)(C)O)c2)CCc2c1[nH]c1ccccc21 | 0 |
| KKL-1155 | Clc1ccc(CNCc2c[nH]nc2-c2ccc(-c3ccccc3)cc2)s1 | 0 |
| KKL-1156 | COC(=O)c1ccc(CN2CCC(C(=O)Nc3cccc(-c4ccco4)c3)CC2)cc1 | 0 |
| KKL-1157 | CSc1ccc(C(=O)C2CCCN(Cc3ccc(C#CCCO)cc3)C2)cc1 | 2 |
| KKL-1158 | COc1cccc(-c2ccccc2NC(=O)C2CCN(CCC(C)c3ccccc3)CC2)c1 | 0 |
| KKL-1159 | Cc1cccc(-c2cccc(NC(=O)C3CCCN(C4CCSCC4)C3)c2)c1 | 2 |
| KKL-1160 | COC(=O)CCCC(=O)N1CCC(C(=O)Nc2ccc(-c3nc4ccccc4o3)cc2)CC1 | 0 |

| | | |
|---|---|---|
| KKL-1161 | CC(C)CC(O)C1CCN(c2nncc(-c3cccc(F)c3)n2)CC1 | 0 |
| KKL-1162 | Cc1ccc(-n2cc(CN3CCN(C(=O)c4ccco4)CC3)c(-c3ccc(F)cc3)n2)cc1 | 0 |
| KKL-1163 | COc1ccc(-c2ncc(CN3CCCC(C(=O)c4ccc(Cl)cc4)C3)cn2)cc1 | 0 |
| KKL-1164 | CSCCC(=O)NC1CCCc2c1cnn2-c1ccc(C(C)(C)C)cc1 | 2 |
| KKL-1165 | CCC(NCc1cn(-c2ccc(C)cc2)nc1-c1cccc(C)c1)c1ccncc1 | 0 |
| KKL-1166 | CC(C)c1ccc(NC2CCCN(C(=O)c3ccc4[n+]([O-])onc4c3)C2)cc1 | 1 |
| KKL-1167 | COC(=O)c1ccc(-c2nc(CN3CCC4(CC3)C=Cc3ccccc34)c(C)o2)cc1 | 5 |
| KKL-1168 | COc1cccc(-c2ccccc2NC(=O)C2CCN(Cc3ccc4ccccc4c3)CC2)c1 | 0 |
| KKL-1169 | C(NC1CCCc2c1cnn2-c1ccccn1)c1cn(-c2ccccc2)nc1-c1ccccc1 | 0 |
| KKL-1170 | Cc1cccc(-c2ccc(NC(=O)C3CCN(C(=O)CCc4cnn(C)c4)CC3)cc2)c1 | 0 |
| KKL-1171 | C(CNc1nncc(-c2ccc(C3CCCCC3)cc2)n1)Cn1ccnc1 | 8 |
| KKL-1172 | COc1cccc(-c2ccccc2NC(=O)C2CCN(Cc3ccc(F)c(OC)c3)CC2)c1 | 0 |
| KKL-1173 | COC(=O)c1ccc(C(=O)NC2CC(C)(C)Cc3c2cnn3-c2ccc(F)cc2)nc1 | 6 |
| KKL-1174 | COC(=O)c1ccc(C(=O)NC2CCCc3c2cnn3-c2cc(F)cc(F)c2)nc1 | 2 |
| KKL-1175 | COc1cccc(-c2ccccc2NC(=O)C2CCN(Cc3cccc(SC)c3)CC2)c1 | 0 |
| KKL-1176 | CCOC(=O)C1(CC2CC2)CCN(Cc2ccc(OC)c(C)c2C)CC1 | 0 |
| KKL-1177 | Cc1ccc(-n2cc(CNCCc3ccccn3)c(-c3cccc(C)c3)n2)cc1 | 0 |
| KKL-1178 | CN(CCC12CC3CC(CC(C3)C1)C2)Cc1nc(COc2ccccc2)no1 | 0 |
| KKL-1179 | CCOC(=O)C1(CCc2ccccc2)CCN(C(=O)Nc2ccc(SC)cc2)CC1 | 2 |
| KKL-1180 | Cc1ccc(-n2cc(CNCCc3nc(O)c4ccccc4n3)c(-c3cccc(C)c3)n2)cc1 | 0 |
| KKL-1181 | Cc1c(CN2CCC(C(=O)Nc3ccc(-n4cnnn4)cc3)CC2)[nH]c2ccc(F)cc12 | 2 |
| KKL-1182 | COc1ccc(Cc2nc3ccc(C(=O)NCCCSC)cc3o2)cc1 | 0 |
| KKL-1183 | CCn1nc(C)cc1C(=O)N1CCc2c([nH]c3ccccc23)C1c1cc(F)ccc1F | 0 |
| KKL-1184 | O=C(N1CCCCO1)c1cc(COc2ccc3c(c2)CCCC3)on1 | 0 |
| KKL-1185 | CCCc1ccc2oc(C(=O)NCCc3ccc(O)cc3)c(C)c2c1 | 2 |
| KKL-1186 | CN(CC1Cc2ccccc2CN1C)C(=O)c1cc(COc2cncc(Cl)c2)on1 | 0 |
| KKL-1187 | O=Cc1ccc(C(=O)N2CCCC(C(=O)c3ccc(Oc4ccccc4)cc3)C2)cc1 | 2 |
| KKL-1188 | COc1cc(C(=O)N2CCc3c([nH]c4ccccc34)C2c2ccccn2)c2ccccc2n1 | 0 |
| KKL-1189 | COC(=O)[C@@H](NC(=O)c1cc(COc2cccc(C(F)(F)F)c2)on1)c1ccccc1 | 2 |

| | | |
|---|---|---|
| KKL-1190 | O=C(N[C@H]1CC[C@]2(CC1)CC(=O)c1ccccc1O2)c1cc2ccccc2o1 | 0 |
| KKL-1191 | Cc1ccc(NCCc2c([N+]([O-])=O)cc([N+]([O-])=O)cc2[N+]([O-])=O)cc1 | 0 |
| KKL-1192 | CCOC(=O)CSc1nnc(-c2ccc(Cl)cc2Cl)o1 | 0 |
| KKL-1193 | COc1cccc(-n2nnnc2S(=O)(=O)Cc2ccccc2)c1 | 0 |
| KKL-1194 | COc1ccc(C)cc1NC(=O)c1ccc(Cc2ccc(C)cc2)o1 | 7 |
| KKL-1195 | Cc1ccc(CN(CCCn2ccnc2)Cc2cccc(Cl)c2Cl)o1 | 17 |
| KKL-1196 | CCCC(=O)N1C(C)Cc2cc(C(O)CN3CCc4[nH]c5c(C)cccc5c4C3)ccc21 | 0 |
| KKL-1197 | COc1ccc(NCc2cc3ccc(SC)cc3nc2O)c(OC)c1 | 0 |
| KKL-1198 | OCC#CCSc1nnc(-c2cccc3ccccc23)o1 | 1 |
| KKL-1199 | Cc1ccc(Nc2c(-c3ccccn3)nc3ccc(C)cn23)cc1 | 4 |
| KKL-1200 | COc1ccc(OCC(O)Cn2c3ccccc3n(CCN3CCCC3)c2=N)cc1 | 4 |
| KKL-1201 | CCOC(=O)COc1ccc2c(c1)oc(C)c(Oc1cnn(-c3ccccc3)c1)c2=O | 12 |
| KKL-1202 | Cc1ccc(CN(CCCn2ccnc2)Cc2[nH]c3ccc(F)cc3c2C)s1 | 20 |
| KKL-1203 | Cc1ccc(C2=Nc3ccccc3SC(c3cccc([N+]([O-])=O)c3)C2)cc1 | 6 |
| KKL-1204 | [O-][N+](=O)c1ccc(/C=C/C(=O)Nc2ccc(CC#N)cc2)o1 | 0 |
| KKL-1205 | Brc1ccc(C2C=C(c3ccccc3)Nc3ncnn32)cc1 | 1 |
| KKL-1206 | Cc1c(C(=O)N2CCn3cnnc3C2)oc2cc(C)ccc12 | 0 |
| KKL-1207 | COc1ccccc1N1CCN(Cc2nc(-c3ccc(C)cc3)no2)CC1 | 5 |
| KKL-1208 | COc1ccc(NC(=O)Cn2nnc(-c3ccc(OCC(C)C)cc3)n2)c(OC)c1 | 5 |
| KKL-1209 | C(N1CCC2(CC1)Nc1ccccc1-n1cccc12)C1=Cc2ccccc2OC1 | 7 |
| KKL-1210 | OC(CN1CCCCC1)Cn1cc(/C=C/C(=O)c2ccccc2)c2ccccc12 | 0 |
| KKL-1211 | Cc1ccccc1NC(=O)c1ccccc1NC(=S)NC(=O)c1cccnc1 | 9 |
| KKL-1212 | Clc1ccc(C(=O)c2ccc(CSc3nnc(-c4ccccc4)o3)o2)cc1 | 0 |
| KKL-1213 | COc1ccc(NCc2cc3c(C)nn(-c4ccccc4)c3nc2O)c(OC)c1 | 0 |
| KKL-1214 | Cc1cc(N2CCCCC2)nc(N2CCC(O)(Cc3ccccc3)CC2)n1 | 9 |
| KKL-1215 | COc1ccc2[nH]c(CNCCc3nc4cc(Cl)ccc4[nH]3)cc2c1 | 0 |
| KKL-1216 | CCOC(=O)CCc1c(C)c(C#N)c2nc3ccccc3n2c1Cl | 0 |
| KKL-1217 | CCCOC(=O)CSc1nnc(-c2ccc(N(C)C)cc2)o1 | 9 |
| KKL-1218 | CC(C)C(=O)N1C[C@H](c2ccc(F)cc2)[C@H]2CN(Cc3ccccc3C)CC[C@@H]12 | 2 |

| | | |
|---|---|---|
| KKL-1219 | CCOc1ccc2n(C)c(C(=O)N3CCCC(c4cc5[nH]cccc5n4)C3)cc2c1 | 0 |
| KKL-1220 | O=C(NC(=S)N1CCCC1)c1ccc(-c2ccccc2)cc1 | 9 |
| KKL-1221 | Clc1ccc(OCc2ccccc2Cl)c(CNC2CCCCC2)c1 | 0 |
| KKL-1222 | Cc1ccc(CN(CCCn2ccnc2)Cc2ccc(Cl)c(Cl)c2)o1 | 14 |
| KKL-1223 | COc1ccc2c(c1)NC1(COC3(C1)CCN(Cc1cccc(F)c1)CC3)c1cccn1-2 | 0 |
| KKL-1224 | O=C(Nc1nc(-c2ccc3c(c2)CCO3)cs1)c1cccc(Cn2cncn2)c1 | 0 |
| KKL-1225 | COc1ccc2c(c1)NC1(COC3(C1)CCN(C(=O)C1CCCC1)CC3)c1cccn1-2 | 8 |
| KKL-1226 | CC(C)Oc1ccc2c(-c3ccccc3)cc(=O)oc2c1C | 1 |
| KKL-1227 | CN(C)c1ccc(Oc2cc(C)nc(C3CCCN(C(=O)c4cc(C5CC5)on4)C3)c2)cc1 | 0 |
| KKL-1228 | CCCOC(=O)CSc1nnc(-c2ccc(OC)cc2)o1 | 7 |
| KKL-1229 | CC1CC(C)CN(C(=O)c2ccc(CN=c3c(N4CCCC4)c(O)c3=O)cc2)C1 | 6 |
| KKL-1230 | CCCN(CCC)CC(O)Cn1cc(/C=C/C(=O)c2ccccc2)c2ccccc12 | 0 |
| KKL-1231 | CC(C)(C)n1cnc2cc(NC(=O)c3ccc(Cl)cc3)ccc12 | 0 |
| KKL-1232 | COc1cccc2c(OC)c3ccoc3nc12 | 0 |
| KKL-1233 | Cn1cccc1CN1CCC2(CC3(CO2)Nc2cc(Cl)ccc2-n2cccc23)CC1 | 1 |
| KKL-1234 | CC(=O)N1CC2(CCN(C(=O)CCC3=Nc4ccccc4OC3=O)CC2)Nc2ccccc21 | 2 |
| KKL-1235 | CCn1c(CN2CCC3(CC2)Nc2cc(OC)ccc2-n2cccc23)nc2ccccc12 | 6 |
| KKL-1236 | CCC(=O)N1CCc2c([nH]c3ccccc23)C12CCN(Cc1ccccc1)CC2 | 1 |
| KKL-1237 | Cc1cc(C(F)(F)F)c2c(NCc3cccc4cccnc34)[nH]nc2n1 | 6 |
| KKL-1238 | Cn1nc(-c2cccnc2)cc1CN1CCN(C(=O)c2cc3ccc(Cl)cc3[nH]2)CC1 | 3 |
| KKL-1239 | COc1cccc(N2CCN(Cc3nc(N)nc(Nc4ccc(C)c(Cl)c4)n3)CC2)c1 | 0 |
| KKL-1240 | CCCn1nnnc1NC(=O)c1cccs1 | 13 |
| KKL-1241 | CC(C)(C)c1ccc(C(=O)Nc2ccc(S(=O)(=O)Nc3nccs3)cc2)cc1Br | 2 |
| KKL-1242 | Cc1nc(N2CCN(Cc3ccc4c(c3)OCO4)CC2)c2cc(-c3ccccc3)sc2n1 | 9 |
| KKL-1243 | CC(NC(=O)C1(c2ccc(Cl)cc2)CCC1)c1n[nH]c(-c2cccnc2)n1 | 8 |
| KKL-1244 | CN(C)c1ccc(/C=C/c2nc3ccccc3c(=O)n2-c2ccccc2)cc1 | 4 |
| KKL-1245 | COc1cccc(N2CCN(CC(=O)Nc3cc(Cl)ccc3Cl)CC2)c1 | 0 |
| KKL-1246 | CCOC(=O)c1ccc(NC(=O)C2C3CC(C)(Oc4ccccc43)N(C)C2=O)cc1 | 0 |
| KKL-1247 | Cc1ccc(NC(=O)CN2CCC(c3nc4cc(Cl)ccc4[nH]3)CC2)cc1 | 0 |

| | | |
|---|---|---|
| KKL-1248 | CCc1ccc(CCC(=O)Nc2ccc(C)cc2)o1 | 0 |
| KKL-1249 | Cc1cc(OCCCCCn2ccnc2)cc(C)c1Cl | 21 |
| KKL-1250 | Cc1cc(=O)oc2cc(OCC(=O)OC3CCCCC3)ccc12 | 7 |
| KKL-1251 | CC(C)COc1ccc(-c2nnn(CC(=O)Nc3ccc4c(c3)OCCO4)n2)cc1 | 9 |
| KKL-1252 | CCN(CC)C(=S)NC(=O)c1ccc(F)cc1 | 9 |
| KKL-1253 | COc1ccc2c(c1)NC1(CCN(CC3=Cc4ccccc4OC3)CC1)c1cccn1-2 | 0 |
| KKL-1254 | COc1ccc2nc(C)cc(SCC(=O)Nc3cc(Cl)ccc3Cl)c2c1 | 2 |
| KKL-1255 | COc1ccc(-c2cc3ncccc3c(NCCNC(=O)c3scnc3C)n2)cc1 | 0 |
| KKL-1256 | CCN(CC)CCn1c2ccccc2n(CC(O)COc2ccc(Cl)cc2Cl)c1=N | 0 |
| KKL-1257 | Cc1cccc(OCCCn2cnc3cc(C)c(C)cc23)c1 | 10 |
| KKL-1258 | C(Cc1ccccc1)N1CCC2(CC1)Nc1ccccc1-n1cccc12 | 0 |
| KKL-1259 | CN(C)c1ccc(-c2nnc(SCC(=O)OCc3ccccc3)o2)cc1 | 11 |
| KKL-1260 | [O-][N+](=O)c1c(-n2cccn2)nn(-c2ccccc2)c1C(Cl)Cl | 0 |
| KKL-1261 | Cc1cc2ncn(CC(O)COc3ccc(F)cc3)c2cc1C | 3 |
| KKL-1262 | Clc1nc(NC2CC2)c2ccccc2n1 | 0 |
| KKL-1263 | O=C(N1CCCCC1c1nc(Nc2ccccc2)cc(-c2ccncc2)n1)c1ccccn1 | 10 |
| KKL-1264 | CCOC(=O)Cn1nnc(-c2ccc(OCc3ccccc3F)cc2)n1 | 0 |
| KKL-1265 | CCOc1ccccc1NC(=O)CN1CCC(C(=O)c2ccc(F)cc2)CC1 | 3 |
| KKL-1266 | [O-][N+](=O)c1ccc(C(=O)NC(=S)N2CCCCCC2)cc1 | 18 |
| KKL-1267 | CCOc1ccccc1NC(=O)CSc1nc2ccc(NC(=O)c3ccccc3Cl)cc2s1 | 0 |
| KKL-1268 | CC(C)[C@H]1CN(Cc2ccccc2F)CCN1C(=O)C1=CC(=O)c2cc(C)ccc2O1 | 0 |
| KKL-1269 | C1CCN(c2nc(/C=C/c3ccccc3)nc3ccccc23)C1 | 0 |
| KKL-1270 | CC(C)(C)c1cc(Cl)c(O)c(NC(=S)NC(=O)c2cccnc2)c1 | 0 |
| KKL-1271 | COC(=O)COc1ccc(NC(=O)COc2ccc(C(C)(C)C)cc2)cc1 | 0 |
| KKL-1272 | Clc1ccc(OCCCn2ccnc2)cc1Cl | 0 |
| KKL-1273 | Cc1ccc(C(=O)Nc2ccc([N+]([O-])=O)cc2C)cc1I | 0 |
| KKL-1274 | Cc1ccc(OCCCn2ccnc2)c(CC=C)c1 | 0 |
| KKL-1275 | CC(C)(C)NCCCCOc1ccc(Cl)cc1C(C)(C)C | 0 |
| KKL-1276 | COc1cc(/C=C/c2ccc3cccc(O)c3n2)cc([N+]([O-])=O)c1O | 0 |

| | | |
|---|---|---|
| KKL-1277 | COC(=O)COc1ccc(/C=C(/C#N)c2nc3ccccc3[nH]2)cc1 | 0 |
| KKL-1278 | CCc1c(C)c2ccc(OCC(=O)OC)c(C)c2oc1=O | 0 |
| KKL-1279 | Clc1ccc(-c2ccc(C(=O)Nc3cccc(Br)c3)o2)cc1 | 0 |
| KKL-1280 | CCc1ccc(OCC(=O)NNC(=O)c2cnccn2)c(Br)c1 | 3 |
| KKL-1281 | Oc1ccc(Br)cc1-c1nc(-c2cccs2)c(-c2cccs2)[nH]1 | 6 |
| KKL-1282 | FC(F)(F)c1cccc(NC(=O)CC23CC4CC(CC(C4)C2)C3)c1 | 0 |
| KKL-1283 | COC(=O)COc1ccc(-c2ccccc2)cc1 | 4 |
| KKL-1284 | CC(=O)Oc1cccc2ccc(/C=C/c3ccc(Cl)cc3)nc12 | 0 |
| KKL-1285 | [O-][N+](=O)c1cc(C(F)(F)F)ccc1Sc1nnc(-c2ccccc2Cl)n1CC=C | 5 |
| KKL-1286 | Clc1cc(Br)ccc1OCCCCn1ccnc1 | 0 |
| KKL-1287 | CCOC(=O)COc1ccc(-c2ccc(C)cc2)cc1 | 0 |
| KKL-1288 | COc1cccc(NC(=O)CSc2nnc(-c3ccc(Br)cc3)o2)c1 | 6 |
| KKL-1289 | CCCOc1ccccc1-c1n[nH]c(SCC(=O)OCC)n1 | 5 |
| KKL-1290 | Cc1cccc(N=C(S)NCCN2CCN(c3ccc(F)c(Cl)c3)CC2)c1C | 0 |
| KKL-1291 | CCOc1ccc(C(=O)Nc2ccc(N3CCN(C(=O)c4cccs4)CC3)c(Cl)c2)cc<br>1Br | 1 |
| KKL-1292 | OC(=O)c1ccc(NC(=S)NC(=O)c2ccc(-c3ccccc3)cc2)cc1 | 0 |
| KKL-1293 | Brc1ccc2ncnc(Nc3ccc(N4CCCC4)cc3)c2c1 | 0 |
| KKL-1294 | COc1cc(Br)cc2cc(C(=O)Nc3cccc(C(F)(F)F)c3)c(=O)oc12 | 0 |
| KKL-1295 | Cc1cc(OCC(=O)NC2C3CC4CC(C3)CC2C4)cc(C)c1Br | 0 |
| KKL-1296 | COc1ccccc1/C=C/c1ccc2cccc(OC(C)=O)c2n1 | 16 |
| KKL-1297 | CC(=O)Oc1cccc2ccc(/C=C/c3cccs3)nc12 | 33 |
| KKL-1298 | COC(=O)COc1ccc(NC(=O)c2ccc3c(c2)C(=O)N(C(C)(C)C)C3=O)<br>cc1 | 0 |
| KKL-1299 | Cc1nc2ccc(C(=O)OCc3nnc(-c4ccccc4)o3)cc2nc1C | 0 |
| KKL-1300 | CC(C)C(=O)N1CCN(c2ccccc2NC(=O)c2ccc([N+]([O-<br>])=O)o2)CC1 | 0 |
| KKL-1301 | O=C(CC12CC3CC(CC(C3)C1)C2)NC(=S)Nc1ccc(-<br>c2nc3ccccc3[nH]2)cc1 | 0 |
| KKL-1302 | [O-<br>][N+](=O)c1ccc(C(=O)Nc2ccc(N3CCN(Cc4ccccc4F)CC3)cc2)o1 | 0 |
| KKL-1303 | COC(=O)COc1ccc(C2CCCCC2)cc1 | 0 |
| KKL-1304 | CC(C)(C)c1cc(Cl)ccc1OCCn1ccnc1 | 11 |
| KKL-1305 | CC(C)(C)c1cc(Cl)ccc1OCCCCn1ccnc1 | 5 |

| | | |
|---|---|---|
| KKL-1306 | CCOc1ccc(NC(=O)c2c(NC(=O)CNCCN3CCOCC3)sc3c2CCC3)cc1 | 1 |
| KKL-1307 | CCC(C)c1ccc(OCCn2ccnc2)cc1 | 13 |
| KKL-1308 | Cc1nc2ccc(C(=O)OCc3cccc(F)c3)cc2nc1C | 0 |
| KKL-1309 | FC(F)(F)c1cccc(N2CCN(C(=O)NC34CC5CC(CC(C5)C3)C4)CC2)c1 | 3 |
| KKL-1310 | COc1ccc(NC(=O)c2c(NC(=O)c3ccc([N+]([O-])=O)o3)sc3c2CCC3)cc1 | 0 |
| KKL-1311 | Cc1cccc(NC(=S)NC(=O)CC23CC4CC(CC(C4)C2)C3)c1 | 0 |
| KKL-1312 | [O-][N+](=O)c1ccc(C(=O)Nc2nc3c(Cl)cccc3s2)o1 | 0 |
| KKL-1313 | CC(C(=O)OCc1ccccc1)n1cnc2scc(-c3ccc(Br)cc3)c2c1=O | 0 |
| KKL-1314 | CC(C)(C)c1ccc(C(=O)NC(=S)Nc2ccc(C(O)=O)cc2)cc1 | 0 |
| KKL-1315 | CCC(C)(C)c1ccc(OCCCn2ccnc2)cc1 | 9 |
| KKL-1316 | Clc1ccc(SCCCn2ccnc2)cc1 | 1 |
| KKL-1317 | CCOC(=O)c1ccc(NC(S)=Nc2ccc(Oc3ccccc3)cc2)cc1 | 0 |
| KKL-1318 | [O-][N+](=O)c1ccc(N2CCN(C(S)=NCc3ccc(Cl)cc3)CC2)cc1 | 0 |
| KKL-1319 | [O-][N+](=O)c1cccc(NC(S)=Nc2ccc(Cl)c(C(F)(F)F)c2)c1 | 0 |
| KKL-1320 | CC(C)(C)NC(=O)c1cc2cc([N+]([O-])=O)ccc2oc1=O | 0 |
| KKL-1321 | COc1cccc(C(=O)NC(=S)Nc2ccc(N3CCN(C(=O)c4cccs4)CC3)c(Cl)c2)c1 | 0 |
| KKL-1322 | Clc1ccc(OCCCCn2ccnc2)c(Br)c1 | 0 |
| KKL-1323 | FC(F)C(F)(F)Oc1ccc(NC(=O)Nc2cccc(Cl)c2)cc1 | 0 |
| KKL-1324 | CCCOc1ccc(C(=O)NC(=S)Nc2ccc(C(O)=O)cc2)cc1 | 0 |
| KKL-1325 | Cc1ccc(OCCCCn2ccnc2)c(CC=C)c1 | 1 |
| KKL-1326 | Cc1cc(C(C)(C)C)ccc1OCCCCn1ccnc1 | 12 |
| KKL-1327 | CCc1c(C)sc(N=C(S)NC2CCN(Cc3ccccc3)CC2)c1C(=O)OC | 0 |
| KKL-1328 | Cc1ccc2nsnc2c1S(=O)(=O)Nc1cccc2cccnc12 | 3 |
| KKL-1329 | Cc1ccc(S(=O)(=O)Nc2ccc(NC(=O)Nc3cccc(C(F)(F)F)c3)cc2)cc1 | 0 |
| KKL-1330 | CCN(CC)c1ccc(NC(=O)c2ccc([N+]([O-])=O)o2)c(C)c1 | 0 |
| KKL-1331 | Clc1ccc(CNC(=S)N2CCN(C/C=C/c3ccccc3)CC2)cc1 | 0 |
| KKL-1332 | CC(=O)c1ccc(NC(=O)Nc2cc(OCC(F)(F)F)cc(OCC(F)(F)C(F)F)c2)cc1 | 0 |
| KKL-1333 | CC(=O)c1ccc(NC(S)=Nc2ccc(Br)c(Cl)c2)cc1 | 0 |
| KKL-1334 | COC(=O)c1ccc(OCC(=O)Nc2ccccc2F)cc1 | 0 |

| | | |
|---|---|---|
| KKL-1335 | Fc1ccc(NC(=O)NC2CCCCCCC2)cc1 | 0 |
| KKL-1336 | [O-][N+](=O)c1cccc(NC(=O)Nc2ccc(Cl)c(C(F)(F)F)c2)c1 | 0 |
| KKL-1337 | NC(=O)COc1ccc(-c2ccc(C#N)cc2)cc1 | 0 |
| KKL-1338 | [O-][N+](=O)c1cccc(NC(=O)Nc2ccc(Oc3ccccc3)cc2)c1 | 0 |
| KKL-1339 | CCOC(=O)c1cc(-c2ccccc2)sc1N=C(S)NC1CCN(Cc2ccccc2)CC1 | 0 |
| KKL-1340 | Cc1ccc(CSCC(=O)Nc2ccc(Cl)c(Cl)c2)cc1 | 0 |
| KKL-1341 | CCOc1cc2c3c(c1)C(C)=CC(C)(C)N3C(=O)C21C(C#N)=C(N)Oc2 cc(O)ccc21 | 0 |
| KKL-1342 | Cc1cc(NC(=O)c2ccc([N+]([O-])=O)o2)c(C)cc1I | 0 |
| KKL-1343 | CC12CC3CC(C)(C1)CC(C(=O)Nc1nc4ccccc4s1)(C3)C2 | 2 |
| KKL-1344 | CCOC(=O)CSc1nnc2c3sc4nc(N5CCOCC5)c5c(c4c3ncn12)CC(C )(C)OC5 | 0 |
| KKL-1345 | Cn1nc([N+]([O-])=O)nc1Sc1nnc(-c2cccnc2)o1 | 0 |
| KKL-1346 | CC(=O)N1CC2(CCN(CC(=O)Nc3ccc(Cl)cc3)CC2)Nc2ccccc21 | 0 |
| KKL-1347 | Cc1cc2cc(CN(Cc3cccnc3)C(=O)Nc3ccc(F)cc3)c(O)nc2cc1C | 0 |
| KKL-1348 | CC(C)(C)C(=O)N1CCc2nc(N3CCCC3)nc(Oc3ccc(F)cc3)c2C1 | 0 |
| KKL-1349 | CN(C(=O)c1c[nH]nc1-c1ccc(CNC(=O)c2cccc(Cl)c2)cc1)c1ccccc1 | 4 |
| KKL-1350 | Fc1ccc(NC(=O)Nc2ccc(Cl)cc2)cc1F | 0 |
| KKL-1351 | COC(=O)c1cc(NC(=O)Nc2cccc(Cl)c2)cc(C(=O)OC)c1 | 6 |
| KKL-1352 | CN(C)c1ccc(Nc2nc(-c3ccc(O)cc3O)cs2)cc1 | 0 |
| KKL-1353 | Clc1ccc(C(=O)COc2cccc3ccccc23)cc1 | 0 |
| KKL-1354 | CCCN1C(=O)c2cccc3cc([N+]([O-])=O)cc(c23)C1=O | 0 |
| KKL-1355 | CC1CN(c2ccc([N+]([O-])=O)cc2)CCN1C(=O)c1cccc(Br)c1 | 0 |
| KKL-1356 | CCC(Sc1nc2ccccc2o1)C(=O)Nc1nnc(C)s1 | 2 |
| KKL-1357 | CCn1c(NCc2c(O)ccc3ccccc23)nc2ccccc12 | 0 |
| KKL-1358 | CC(=NNC1=NC(=O)C(Cc2ccc(Cl)cc2)S1)c1ccc(F)cc1 | 16 |
| KKL-1359 | COc1ccccc1NC(=O)CSc1nc(-c2ccccc2)cc(C(F)(F)F)c1C#N | 0 |
| KKL-1360 | Cc1ccc(C)c(N2C(=O)c3cccc4c([N+]([O-])=O)ccc(c34)C2=O)c1 | 0 |
| KKL-1361 | [O- ][N+](=O)c1cccc(NC(=O)CN(c2ccc(Cl)c(Cl)c2)S(=O)(=O)c2ccccc [illegible] | 2 |
| KKL-1362 | Cc1ccc([N+]([O-])=O)cc1NC(=O)Nc1cccc(Cl)c1 | 4 |
| KKL-1363 | CCc1ccccc1N1C(=O)c2cccc3c([N+]([O-])=O)ccc(c23)C1=O | 0 |

| | | |
|---|---|---|
| KKL-1364 | O=C(Oc1cccc2cccnc12)C(Cc1ccccc1)N1C(=O)C2C3CCC(C3)C2C1=O | 9 |
| KKL-1365 | CN(C)c1ccc(NC(=O)c2ccncc2)cc1 | 0 |
| KKL-1366 | OC1=C2C(c3ccc(C(F)(F)F)cc3)Nc3ccccc3N=C2CC(c2ccc(Cl)cc2)C1 | 0 |
| KKL-1367 | Brc1ccc(Nc2nc(-c3ccccn3)cs2)cc1 | 0 |
| KKL-1368 | Fc1ccc(C(=O)Nc2ncnc3sc4c(c23)CCCC4)cc1 | 0 |
| KKL-1369 | FC(F)(F)c1cccc(NC(=O)/C=C/c2cccc(C(F)(F)F)c2)c1 | 0 |
| KKL-1370 | Clc1ccc(OCCCCn2ccnc2)c(CC=C)c1 | 0 |
| KKL-1371 | CCC(C)c1ccc(OCCCCn2ccnc2)cc1 | 14 |
| KKL-1372 | CCOC(=O)c1cc(CC)sc1NC(=O)c1cc2ccccc2oc1=O | 0 |
| KKL-1373 | Cc1ccc(NC(=O)C(Cl)Cl)cc1[N+]([O-])=O | 4 |
| KKL-1374 | CCN1C(=O)c2cccc3cc([N+]([O-])=O)cc(c23)C1=O | 0 |
| KKL-1375 | Cc1cc(C)c2[nH]c3ccccc3c2n1 | 0 |
| KKL-1376 | CC(=O)N1CCN(c2ccc(NC(=S)NC(=O)c3cc4ccccc4o3)cc2)CC1 | 0 |
| KKL-1377 | Cn1c(NC(=O)c2ccccc2)nc2ccccc12 | 0 |
| KKL-1378 | CCOC(=O)/C=C/C(=O)Nc1nc(-c2ccccc2)cs1 | 0 |
| KKL-1379 | CC(C)CC(=O)NC(c1ccc(Cl)cc1Cl)c1c(O)ccc2ccccc12 | 13 |
| KKL-1380 | Cc1cc(NS(=O)(=O)c2ccc(NC(=O)c3ccc(C(C)(C)C)cc3)cc2)no1 | 0 |
| KKL-1381 | N=c1n(Cc2ccco2)cnc2n(Cc3ccco3)c(-c3ccccc3)c(-c3ccccc3)c12 | 0 |
| KKL-1382 | Clc1ccc(OCCCCn2ccnc2)cc1Cl | 7 |
| KKL-1383 | Cc1ccc(C(=O)Oc2ccc(Cl)c3cccnc23)cc1[N+]([O-])=O | 0 |
| KKL-1384 | CCOc1ccc(NCCC(=O)c2cccs2)cc1 | 0 |
| KKL-1385 | Brc1ccccc1C(=O)Oc1cccc2cccnc12 | 0 |
| KKL-1386 | O=c1oc2ccc3ccccc3c2cc1C#N | 0 |
| KKL-1387 | COC(=O)COc1ccc(NC(=O)c2ccccc2F)cc1 | 0 |
| KKL-1388 | CC(C)(C)c1cnc(NC(=O)c2ccccc2O)s1 | 0 |
| KKL-1389 | COc1ccc(/C=C/c2ccc3cccc(O)c3n2)cc1 | 0 |
| KKL-1390 | Clc1cccc(N2CCN(CCCn3c(=S)[nH]c4ccccc4c3=O)CC2)c1 | 0 |
| KKL-1391 | CC(=O)Oc1cccc2ccc(/C=C/c3ccc(F)cc3)nc12 | 9 |
| KKL-1392 | Cc1ccc(-c2nnc(SCC#CCOC(=O)c3ccco3)o2)cc1 | 0 |

| | | |
|---|---|---|
| KKL-1393 | Cc1ccc(C(=O)NCCC2=CCCCC2)cc1Br | 0 |
| KKL-1394 | Cc1nn(-c2ccccc2)c(C)c1N1C(=O)c2cccc3cc([N+]([O-])=O)cc(c23)C1=O | 1 |
| KKL-1395 | Clc1cccc(NCCC(=O)c2ccco2)c1 | 0 |
| KKL-1396 | CCOC(=O)COc1ccc(S(=O)(=O)N2CCc3ccccc3C2)cc1 | 2 |
| KKL-1397 | CCCn1nnc(NC(=O)COc2ccc(C3CCCCC3)cc2)n1 | 0 |
| KKL-1398 | COC(=O)c1ccc(N2C(=O)CC(N(CCc3ccccc3)C(S)=Nc3ccccc3)C2=O)cc1 | 3 |
| KKL-1399 | Nc1nnc(Sc2ccc([N+]([O-])=O)c3ncccc23)s1 | 0 |
| KKL-1400 | COc1ccccc1-c1coc2cc(OCC(=O)OC3CCCCC3)ccc2c1=O | 0 |
| KKL-1401 | Cc1ccc(NC(=O)C23CC4CC(C2)CCC(C4)C3)cc1 | 0 |
| KKL-1402 | [O-][N+](=O)c1ccc(C(=O)Nc2ccc(F)c(Cl)c2)o1 | 0 |
| KKL-1403 | Cc1ccnc(N2C(=O)c3cccc4c([N+]([O-])=O)ccc(c34)C2=O)c1 | 0 |
| KKL-1404 | CCOC(=O)C1=C(c2ccc(C)cc2)C(=O)c2ccccc2C1=O | 6 |
| KKL-1405 | CCOC(=O)c1ccc(NC(=O)CSc2nnc(-c3ccc(OC)cc3)o2)cc1 | 4 |
| KKL-1406 | Cc1ccc2n(CC(O)CN3CCN(CCO)CC3)c(-c3ccccc3)c(-c3ccccc3)c2c1 | 4 |
| KKL-1407 | CCOC(=O)COc1ccc(S(=O)(=O)Nc2ccc(C)c(C)c2)cc1 | 0 |
| KKL-1408 | CC(=O)c1ccc2[nH]c3ccc(C(C)=O)cc3c2c1 | 0 |
| KKL-1409 | Nc1c2c(nc3sc4c(c13)CCCC4)CCCC2 | 0 |
| KKL-1410 | Cc1ccc(C(=O)NCCC2=CCCCC2)cc1I | 2 |
| KKL-1411 | Cc1ccc(-c2csc3nc(SCC(N)=O)n(-c4ccccc4)c(=O)c23)cc1C | 0 |
| KKL-1412 | COc1ccc2c(c1)CCC1C2=Nc2ccccc2S1 | 0 |
| KKL-1413 | CCC(C)Oc1ccc(C(=O)NC(=S)Nc2ccc(NC(=O)c3ccco3)cc2)cc1 | 4 |
| KKL-1414 | CCOc1cc(C(=O)ON=C(N)Cc2ccc(Cl)cc2Cl)cc(OCC)c1OCC | 0 |
| KKL-1415 | Cc1ccc(OCCN2CCN(c3nc(NCc4ccccc4)c4ccccc4n3)CC2)cc1 | 0 |
| KKL-1416 | [O-][N+](=O)c1ccc2c3c(cccc13)C(=O)N(c1cccc(C(=O)NC3CCCCC3)[illegible] | 0 |
| KKL-1417 | CCOc1ccc(NC(=O)CSc2nc3ccc([N+]([O-])=O)cc3s2)cc1 | 0 |
| KKL-1418 | [O-][N+](=O)c1ccccc1C(=O)Oc1c(Cl)cc(Cl)c2cccnc12 | 90 |
| KKL-1419 | CCCc1cc2c(cc1OCC(=O)OCC)occ(-c1ccc3c(c1)OCCCO3)c2=O | 0 |
| KKL-1420 | CCOC(=O)CCc1c(C)c(C#N)c2nc3ccccc3n2c1O | 0 |
| KKL-1421 | COC(=O)C(C)Oc1ccc2c(c1)occ(Oc1ccccc1Br)c2=O | 0 |

| | | |
|---|---|---|
| KKL-1422 | [O-][N+](=O)c1cc(C(F)(F)F)ccc1SCC(=O)OC1CCCCC1 | 1 |
| KKL-1423 | CCC(=O)Oc1ccc(C(=O)Nc2cc3oc4ccccc4c3cc2OC)cc1 | 0 |
| KKL-1424 | CC1CCc2nc3sc4c(c3c(N)c2C1)CCCC4 | 0 |
| KKL-1425 | Cc1cc(C(=O)NCC23CC4CC(CC(C4)C2)C3)ccc1[N+]([O-])=O | 0 |
| KKL-1426 | CN1CCN(c2ccc([N+]([O-])=O)c(-n3nc(-c4cc(C)ccc4C)c4ccccc4c3=O)c2)CC1 | 0 |
| KKL-1427 | CC1CCCN(CCCOc2ccc(Cl)c(C)c2C)C1 | 0 |
| KKL-1428 | CCC(C)NCCCOc1ccc(Cl)c2ccccc12 | 0 |
| KKL-1429 | CCC(C)Oc1ccc(C(=O)NC(=S)Nc2ccc3c(c2)COC3=O)cc1 | 0 |
| KKL-1430 | CC1CCN(C(=S)NC(=O)c2cccc(C)c2)CC1 | 0 |
| KKL-1431 | O=C(Nc1ccc(C(=O)NCC2COc3ccccc3O2)cc1)C1COc2ccccc2O1 | 0 |
| KKL-1432 | CCOC(=O)c1ccc(NC(=S)NC(=O)CCc2ccccc2)cc1 | 0 |
| KKL-1433 | Cc1ccc(-n2cnc3cc(NCc4cccs4)ccc23)cc1 | 3 |
| KKL-1434 | CCCOc1ccc(C2Oc3ccc(Br)cc3C=C2[N+]([O-])=O)cc1OC | 2 |
| KKL-1435 | CC(C)(C)c1ccc(C(=O)NC(=S)Nc2ccc(F)cc2)cc1Br | 13 |
| KKL-1436 | CC(C)(C)c1ccc(C(=O)NC(=S)N2CCCC2)cc1 | 12 |
| KKL-1437 | COc1ccc(C(=O)NC(=S)Nc2ccc([N+]([O-])=O)cc2O)cc1Br | 5 |
| KKL-1438 | C=CCn1c(-c2cccs2)nc2ccccc12 | 0 |
| KKL-1439 | CCc1ccc(O)c(NC(=S)NC(=O)c2cccc3ccccc23)c1 | 1 |
| KKL-1440 | CN(Cc1ccccc1)C(=S)NC(=O)c1cccs1 | 17 |
| KKL-1441 | CCOC(=O)COc1ccc2c(C)c(CC(=O)OC)c(=O)oc2c1 | 1 |
| KKL-1442 | COc1cc([N+]([O-])=O)ccc1NC(=O)c1cc(Cl)ccc1O | 0 |
| KKL-1443 | S=C(NCCc1c[nH]c2ccccc12)Nc1ccc(Nc2ccccc2)cc1 | 6 |
| KKL-1444 | Clc1ccc(CCNC(=S)Nc2ccc(Nc3ccccc3)cc2)cc1 | 4 |
| KKL-1445 | O=C(NC(=S)N1CCc2ccccc21)c1ccccc1 | 17 |
| KKL-1446 | CCCCc1ccc(NC(=O)c2cc3cc(Br)cc(OC)c3oc2=O)cc1 | 8 |
| KKL-1447 | CCN(CC)C(=S)NC(=O)c1ccc(Cl)cc1 | 20 |
| KKL-1448 | FC(F)(F)c1cc(NC(=O)c2csc3c2CCCC3)ccc1Cl | 2 |
| KKL-1449 | [O-][N+](=O)c1ccc(C(=O)CCNc2cccc(Cl)c2)cc1 | 0 |
| KKL-1450 | CCOc1ccc(OCC(=O)Nc2cc(C)cc(C)c2)cc1 | 7 |

| | | |
|---|---|---|
| KKL-1451 | CC(C)CNS(=O)(=O)c1ccc(CCC(=O)Nc2ccc(Cl)cc2C)cc1 | 2 |
| KKL-1452 | CN(Cc1ccccc1)C(=S)NC(=O)c1ccc(C)cc1 | 2 |
| KKL-1453 | OC(COc1ccc(C(=O)c2ccc(Cl)cc2)cc1)CN1CCN(Cc2ccccc2)CC1 | 0 |
| KKL-1454 | Fc1ccccc1NC(=O)CSc1nnc(SCc2cccc3ccccc23)s1 | 7 |
| KKL-1455 | CSc1ccc(C(CC(=O)c2ccccc2)Nc2ccc(F)cc2)cc1 | 3 |
| KKL-1456 | CCOC(=O)COc1ccc2n(CC)c(C)c(C(=O)OCC)c2c1 | 0 |
| KKL-1457 | CCOC(=O)COc1ccc(S(=O)(=O)N2CCC(C)CC2)cc1 | 0 |
| KKL-1458 | COC(=O)COc1ccc(NC(=O)COc2ccc(C)cc2C)cc1 | 4 |
| KKL-1459 | Fc1ccccc1NC(=O)CSc1ccc(Br)cc1 | 0 |
| KKL-1460 | CCOC(=O)COc1ccc(S(=O)(=O)Nc2c(C)cccc2CC)cc1 | 0 |
| KKL-1461 | CCCC(=O)N1CCN(c2ccc(NC(=O)c3ccc(-c4ccc(Cl)cc4)o3)cc2Cl)CC1 | 0 |
| KKL-1462 | CCCCSc1ccc(NCCC(=O)c2ccco2)cc1 | 3 |
| KKL-1463 | CCC(=O)Oc1cccc2ccc(/C=C/c3ccccc3F)nc12 | 14 |
| KKL-1464 | Oc1cccc2ccc(/C=C/c3cccc(F)c3)nc12 | 18 |
| KKL-1465 | COC(=O)c1ccc(COC(=O)c2ccccc2Cl)cc1 | 0 |
| KKL-1466 | COc1cc(NC(=O)c2cc3ccccc3o2)ccc1NC(=O)c1ccccc1Cl | 0 |
| KKL-1467 | CN(c1ccccc1)c1ccc([N+]([O-])=O)c2nonc12 | 0 |
| KKL-1468 | CCOC(=O)COc1ccc(C(=O)NC2CCCCC2)cc1 | 1 |
| KKL-1469 | Cc1ccc(NC(=O)c2ccc(OCC(=O)OC3CCCCC3)cc2)cc1 | 11 |
| KKL-1470 | CC(=O)c1ccc(NCCC(=O)c2ccco2)cc1 | 4 |
| KKL-1471 | Cc1cccc(C)c1OCC(=O)NN=Cc1ccc(O)c([N+]([O-])=O)c1 | 8 |
| KKL-1472 | COc1ccc(C(=O)NC(=S)Nc2ccc(N3CCCCC3)cc2)cc1OC | 0 |
| KKL-1473 | COc1ccc(-c2cc(C(=O)NN=C/C=C/c3ccccc3OC)[nH]n2)cc1 | 0 |
| KKL-1474 | CCCn1c(CNc2ccccc2O)nc2ccccc12 | 0 |
| KKL-1475 | COC(=O)c1c(NC(=O)c2c(F)c(F)c(F)c(F)c2F)sc2c1CCCCC2 | 8 |
| KKL-1476 | CCn1cc(C(O)=O)c(=O)c2cc(F)c(N3CCN(C(S)=Nc4cc(C(F)(F)F)ccc4Cl)CC3)cc12 | 5 |
| KKL-1477 | CC(N1C(=O)C2C(C3C=CC2C2CC32)C1=O)C(=O)Oc1cccc2cccnc12 | 5 |
| KKL-1478 | CC(C)OC(=O)COc1ccc2c(-c3cccc([N+]([O-])=O)c3)cc(=O)oc2c1 | 0 |
| KKL-1479 | COC(=O)c1sccc1NC(=O)c1c(F)c(F)c(F)c(F)c1F | 0 |

| | | |
|---|---|---|
| KKL-1480 | [O-<br>][N+](=O)c1ccc(NC(=O)CN2CCN(Cc3ccccc3)CC2)c(C(=O)c2ccc[illegible] | 0 |
| KKL-1481 | CCN(CC)S(=O)(=O)c1ccc(Cl)c(C(=O)Oc2cccc3cccnc23)c1 | 4 |
| KKL-1482 | Oc1c(Br)cc(NS(=O)(=O)c2ccc(F)cc2)c2ccccc12 | 0 |
| KKL-1483 | Cc1ccc(Sc2ccc(C=O)o2)cc1 | 0 |
| KKL-1484 | CC12CC3CC(C)(C1)CC(CNCc1ccccc1O)(C3)C2 | 3 |
| KKL-1485 | CCOc1ccc2nc3cc(N)ccc3c(N)c2c1 | 1 |
| KKL-1486 | Cc1ccc(OCC(O)CN2CCN(CC(O)COc3ccc(C)cc3C)CC2)c(C)c1 | 2 |
| KKL-1487 | CCC(=O)NC(=S)Nc1ccc(NC(=O)c2cc(I)ccc2Cl)cc1 | 7 |
| KKL-1488 | O=C(NC(=S)Nc1ccccc1N1CCCC1)c1cccc2ccccc12 | 12 |
| KKL-1489 | CCC(C)NCCOc1ccc(C(C)(C)c2ccccc2)cc1 | 1 |
| KKL-1490 | Cc1cccc2sc(NC(=O)c3ccc(C(C)(C)C)cc3)nc12 | 2 |
| KKL-1491 | COC(=O)COc1cccc(NC(=O)c2cccc(C)c2)c1 | 1 |
| KKL-1492 | Cc1oc2c(c1C(=O)Nc1cccc3ccccc13)CCCC2 | 0 |
| KKL-1493 | COC(=O)CSc1nc(N2CCCCC2)c2c(c1C#N)CCCC2 | 0 |
| KKL-1494 | FC(F)(F)c1cccc(/C=C/C(=O)Nc2cccc(Br)c2)c1 | 0 |
| KKL-1495 | CN(C)c1ccc(C(O)(c2ccc(N(C)C)cc2)c2ccc(N(C)C)cc2)cc1 | 0 |
| KKL-1496 | CCOC(=O)COc1ccc2c3c(c(=O)oc2c1)CCCCC3 | 0 |
| KKL-1497 | CCOC(=O)c1ccc(COC(=O)c2cccc([N+]([O-])=O)c2)cc1 | 3 |
| KKL-1498 | CCOc1ccc(NC(=O)CSc2nnc(-c3ccccc3Cl)o2)cc1 | 2 |
| KKL-1499 | CC(=O)Oc1cccc2ccc(/C=C/c3c(F)cccc3Cl)nc12 | 0 |
| KKL-1500 | Cc1ccc(NC(=O)C(=O)NNC(=O)c2ccco2)cc1Cl | 0 |
| KKL-1501 | Ic1ccccc1C(=O)Nc1ccc(-c2nc3ccccc3c(=O)o2)cc1 | 2 |
| KKL-1502 | CC1=NN(C(=O)c2ccccc2)C(O)(c2ccncc2)C1 | 0 |
| KKL-1503 | CCOc1cccc(N2C(=O)CC(SC(=N)NN=C(C)c3ccc(Cl)s3)C2=O)c1 | 1 |
| KKL-1504 | COC(=O)COc1ccc(NC(=O)COc2cccc(C)c2)cc1 | 0 |
| KKL-1505 | Cc1ccc(NC(=O)c2cc3cc(Cl)cc(Cl)c3oc2=O)c(C)c1 | 7 |
| KKL-1506 | COC(=O)C(Oc1ccc2c(c1)oc(=O)c1cc(OC)ccc21)c1ccccc1 | 0 |
| KKL-1507 | Clc1c(C(=O)Nc2nnc(C3CCCCC3)s2)sc2ccccc12 | 1 |
| KKL-1508 | COc1ccccc1N1C(=O)c2cccc3cc([N+]([O-])=O)cc(c23)C1=O | 0 |

| | | |
|---|---|---|
| KKL-1509 | CC(Oc1cccc(C)c1C)C(=O)Nc1ccc([N+]([O-])=O)cc1 | 4 |
| KKL-1510 | COc1ccc(OCc2n(CC(=O)OC3CC(C)CCC3C(C)C)c3ccccc3[n+]2C)cc1 | 0 |
| KKL-1511 | Cc1ccc2nc(-c3ccccn3)c(Nc3ccccc3C)n2c1 | 0 |
| KKL-1512 | COC(=O)CSc1ccc2c(N)c(C(=O)OC)sc2c1 | 0 |
| KKL-1513 | CCOC(=O)C1CCCN(c2ccc([N+]([O-])=O)c3nonc23)C1 | 0 |
| KKL-1514 | CCOC(=O)CSc1nnc(-c2cc(OC)c(OC)c(OC)c2)o1 | 0 |
| KKL-1515 | Cc1cccc(OCC(O)Cn2c3ccccc3n(CCN3CCCCC3)c2=N)c1 | 16 |
| KKL-1516 | CCOC(=O)COc1ccc2c(c1)occ(-c1ccccc1)c2=O | 4 |
| KKL-1517 | [O-][N+](=O)c1nnn([C@@H]2[C@@H](c3ccccc3)CC(c3cccs3)=CC2[illegible] | 0 |
| KKL-1518 | CCOC(=O)COc1ccc2ccccc2c1/C=C/[N+]([O-])=O | 0 |
| KKL-1519 | CN(c1ccccc1)c1nc(N2CCCC2)nc(-n2ccnc2)n1 | 0 |
| KKL-1520 | COc1ccc(CN2CCN(CC(O)Cn3c4ccccc4c4ccccc34)CC2)cc1 | 0 |
| KKL-1521 | Cc1ccn2c(Nc3ccccc3)c(-c3ccccn3)nc2c1 | 0 |
| KKL-1522 | C1CCCN(c2nc(-c3ccccc3)nc(-n3ccnc3)n2)CC1 | 1 |
| KKL-1523 | Cc1ccc(Nc2c(-c3ccccn3)nc3cc(C)ccn23)cc1 | 0 |
| KKL-1524 | Fc1ccc(NC(=O)CSc2n[nH]c(-c3ccc(Cl)cc3)n2)cc1Cl | 0 |
| KKL-1525 | [O-][N+](=O)c1cccc(NC(=O)Nc2ccc(OC(F)(F)F)cc2)c1 | 0 |
| KKL-1526 | COc1ccc(-c2c(C)nn3c(O)c(Cc4ccccc4)c(C)nc23)cc1 | 4 |
| KKL-1527 | OC(COc1ccc2ccccc2c1)CN1CCCN(CC(O)COc2ccc3ccccc3c2)CC1 | 0 |
| KKL-1528 | COc1ccc(CCN=C(S)N2CCN(C/C=C/c3ccccc3)CC2)cc1OC | 2 |
| KKL-1529 | CC(Nc1nc(N2CCN(C(=O)c3ccc(Cl)cc3)CC2)nc2ccccc12)c1ccccc1 | 4 |
| KKL-1530 | CCOC(=O)c1c(NC(=O)COc2ccc(C(C)C)cc2)sc2c1CCC2 | 0 |
| KKL-1531 | Cc1cc([N+]([O-])=O)ccc1NC(=O)Nc1cccc(C(F)(F)F)c1 | 0 |
| KKL-1532 | CCOC(=O)COc1ccc2c(occ(-c3ccccc3F)c2=O)c1C | 0 |
| KKL-1533 | CC(NC(=O)Nc1ccc(C(C)=O)cc1)C12CC3CC(CC(C3)C1)C2 | 0 |
| KKL-1534 | CCOC(=O)c1cc(-c2ccccc2)sc1N=C(S)NC(C)c1ccncc1 | 0 |
| KKL-1535 | COc1ccc(C(C)(C)C)cc1N=C(S)Nc1ccc(Cl)c(Cl)c1 | 0 |
| KKL-1536 | COc1cc([N+]([O-])=O)ccc1N=C(S)Nc1ccc(F)c(Cl)c1 | 0 |
| KKL-1537 | CCOC(=O)c1ccc(NC(S)=Nc2ccc(F)c(Cl)c2)cc1 | 0 |

| | | |
|---|---|---|
| KKL-1538 | CCC(C)c1ccc(OCC(=O)Nc2ccc(OC)c([N+]([O-])=O)c2)cc1 | 4 |
| KKL-1539 | Oc1c(Sc2cccc3cccnc23)cc(NS(=O)(=O)c2ccccc2)c2ccccc12 | 0 |
| KKL-1540 | [O-][N+](=O)c1ccc(NC(S)=NCc2ccc(Cl)c(Cl)c2)cc1 | 0 |
| KKL-1541 | COc1ccc(C(=O)Oc2ccc(C(=O)OCc3ccccc3)cc2)cc1OC | 0 |
| KKL-1542 | CCOC(=O)Cn1nnc(-c2ccc(OCc3ccccc3C)cc2)n1 | 0 |
| KKL-1543 | CCN(CC)c1ccc(NC(S)=NCc2ccc(Cl)cc2)cc1 | 2 |
| KKL-1544 | COc1ccc(OC)c(NC(=O)Nc2ccc(Cl)c(C(F)(F)F)c2)c1 | 0 |
| KKL-1545 | CC(NC(=O)Nc1cccc(C(F)(F)F)c1)C1CC2CCC1C2 | 0 |
| KKL-1546 | Cc1cc(C)cc(COc2ccc3c(C)c(CCC(O)=O)c(=O)oc3c2C)c1 | 0 |
| KKL-1547 | CCc1cc(C(=O)OC)c(N=C(S)NCc2ccc(Cl)c(Cl)c2)s1 | 0 |
| KKL-1548 | Cn1nc([N+]([O-])=O)nc1Sc1ncnn1-c1ccccc1 | 0 |
| KKL-1549 | CCOC(=O)c1c(N=C(S)NC2CCN(Cc3ccccc3)CC2)sc(C)c1-c1ccccc1 | 0 |
| KKL-1550 | CCN(CC)c1ccc(NC(S)=NCCc2ccc(OC)c(OC)c2)cc1 | 0 |
| KKL-1551 | COC(=O)Cc1c(C)c2ccc(OCC(=O)OCc3ccccc3)c(C)c2oc1=O | 4 |
| KKL-1552 | CC(NS(=O)(=O)c1ccc(Cl)cc1)C(=O)Nc1ccc(C)c(Cl)c1 | 0 |
| KKL-1553 | CCc1cc(I)ccc1NC(=O)c1ccc([N+]([O-])=O)o1 | 0 |
| KKL-1554 | Oc1ccccc1-c1n[nH]c(/C=C/c2ccc(Cl)cc2)n1 | 0 |
| KKL-1555 | COc1ccc(OC(C)C(=O)Nc2ccc(Cl)cc2C)cc1 | 4 |
| KKL-1556 | CC(C)(C)c1ccc(Cn2cnc([N+]([O-])=O)n2)cc1 | 0 |
| KKL-1557 | COc1ccc(OC(C)C(=O)Nc2ccc(I)cc2)cc1 | 0 |
| KKL-1558 | Cc1c(OC(=O)c2ccccc2Cl)ccc2c1CCCC2 | 0 |
| KKL-1559 | COc1cccc(OC)c1C(=O)Nc1ccc(Cl)c(-c2nc3ccccc3s2)c1 | 0 |
| KKL-1560 | CC1CCN(c2ccc(NC(=O)c3ccc([N+]([O-])=O)o3)cc2)CC1 | 0 |
| KKL-1561 | Cc1c(OC(=O)c2cccc([N+]([O-])=O)c2)ccc2c1CCCC2 | 1 |
| KKL-1562 | COC(=O)COc1ccc(NC(=O)c2ccc(-c3ccccc3)cc2)cc1 | 4 |
| KKL-1563 | COC(=O)COc1cccc(NC(=O)c2ccc(Cl)cc2)c1 | 7 |
| KKL-1564 | COC(=O)COc1ccc(NC(=O)c2ccccc2C)cc1 | 0 |
| KKL-1565 | CCOc1cc(CNCCO)cc(Br)c1OCc1ccc(Cl)cc1Cl | 0 |
| KKL-1566 | Cc1nn(-c2nc3ccccc3[nH]2)c(N)c1-c1ccccc1 | 0 |

| | | |
|---|---|---|
| KKL-1567 | Cc1ccc(-n2cc(-c3ccccc3)c3c(NCc4cccnc4)ncnc23)cc1 | 0 |
| KKL-1568 | COC(=O)COc1ccc(NC(=O)c2cccc(C)c2)cc1 | 0 |
| KKL-1569 | COC(=O)COc1ccc(NC(=O)c2ccccc2I)cc1 | 0 |
| KKL-1570 | NC(=O)c1c(NC(=O)c2c(F)c(F)c(F)c(F)c2F)sc2c1CCC2 | 0 |
| KKL-1571 | Clc1ccc(SCC(=O)Nc2ccc(C#N)cc2)cc1 | 0 |
| KKL-1572 | COC(=O)COc1ccc(NC(=O)c2ccccc2Br)cc1 | 0 |
| KKL-1573 | FC(F)(F)c1cccc(-n2cc(-c3ccccc3)c3c(NCc4cccnc4)ncnc23)c1 | 0 |
| KKL-1574 | Fc1cccc(NC(=O)Nc2ccc(C(F)(F)F)cc2)c1 | 0 |
| KKL-1575 | FC(F)(F)c1ccc(NC(=O)Nc2cccc(Cl)c2)cc1 | 2 |
| KKL-1576 | COc1ccc(C2(C(=O)Nc3ccc(Br)cc3)CCCC2)cc1 | 0 |
| KKL-1577 | Brc1ccc(SCC(=O)Nc2ccc(C#N)cc2)cc1 | 0 |
| KKL-1578 | CCOC(=O)c1c(C)c(C(=O)OC)sc1NC(=O)c1c(F)c(F)c(F)c(F)c1F | 0 |
| KKL-1579 | CC(C)C(=O)N1CCN(c2ccc(NC(=O)c3ccc([N+]([O-<br>])=O)o3)cc2)CC1 | 0 |
| KKL-1580 | CCOC(=O)Cn1cc(C#N)c2cccc(CC)c12 | 3 |
| KKL-1581 | CC(C)NS(=O)(=O)c1ccc(CCC(=O)Nc2ccc(Cl)cc2C)cc1 | 0 |
| KKL-1582 | CCOc1cc(CNC(C)C23CC4CC(CC(C4)C2)C3)cc(Cl)c1OCC(=O)N<br>C(C)(C)C | 0 |
| KKL-1583 | CCOC(=O)c1ccc(NC(S)=NCCN2CCc3ccccc3C2)cc1 | 0 |
| KKL-1584 | CC(C)c1ccc(Cn2cnc3cc(C)c(C)cc23)cc1 | 0 |
| KKL-1585 | O=C(OCc1ccc(C(=O)Oc2ccccc2)cc1)c1ccco1 | 0 |
| KKL-1586 | Cc1ccc(NC(=O)CC2COc3ccccc3O2)cc1Cl | 0 |
| KKL-1587 | OC1(c2c[nH]c3ccccc23)C(=O)N(Cc2ccccc2Cl)c2ccc(Br)cc21 | 0 |
| KKL-1588 | [O-][N+](=O)c1cccc(NC(S)=Nc2ccc(Cl)c(Cl)c2)c1 | 0 |
| KKL-1589 | Cn1nc(-<br>c2ccco2)cc1[C@H]1CN2CC[C@H]1C[C@@H]2COC(=O)Nc1ccc<br>[illegible] | 0 |
| KKL-1590 | Cc1ccn2c(Nc3ccccc3C)c(-c3ccccn3)nc2c1 | 0 |
| KKL-1591 | CC(C)=CCCC1(C)Oc2c(C)cc3c([nH]c4ccccc34)c2C=C1 | 0 |
| KKL-1592 | CCC(CC)c1cc([C@H]2CN3CC[C@H]2C[C@@H]3CNCc2ccc(CO<br>)o2)nc(-c2ccncc2)n1 | 0 |
| KKL-1593 | Cc1c(Cc2ccccc2)c(NCCCO)n2c(nc3ccccc23)c1C#N | 0 |
| KKL-1594 | COC(=O)[C@@H]1Cc2c([nH]c3ccccc23)[C@H]2C[C@@H](NC(<br>C)C)C[C@@H](c3cccc(Cl)c3)N12 | 1 |
| KKL-1595 | COCCN[C@H]1C[C@H]2N([C@H](C(=O)OC)Cc3c2[nH]c2ccccc<br>32)[C@H](c2ccc(C(F)(F)F)cc2)C1 | 0 |

| | | |
|---|---|---|
| KKL-1596 | COC(=O)c1cccc(NC(=S)NC[C@H]2C[C@@H]3CCN2C[C@@H]3c2cc(-c3ccc4ccccc4c3)nn2C)c1 | 71 |
| KKL-1597 | CCc1ccc(NC(=O)c2cnc(N3CCN(c4ccccn4)CC3)c3ccccc23)cc1 | 4 |
| KKL-1598 | Cc1nc([C@H]2CN3CC[C@H]2C[C@@H]3CNC(=O)c2ccc(C(C)(C)C)cc2)cc(-c2ccccc2Br)n1 | 0 |
| KKL-1599 | CCC(CC)c1cc([C@H]2CN3CC[C@H]2C[C@@H]3CNCc2ccc(OC)cc2)nc(-c2ccncc2)n1 | 0 |
| KKL-1600 | COC(=O)[C@@H]1Cc2c([nH]c3ccccc23)[C@H]2C[C@H](NC3CC3)C[C@@H](c3ccc(C(F)(F)F)cc3)N12 | 0 |
| KKL-1601 | OC1=C(c2ccccc2)C2CCCN2C1=O | 0 |
| KKL-1602 | COC(=O)[C@@H]1Cc2c([nH]c3ccccc23)[C@H]2C[C@@H](NCCO)C[C@@H](c3ccc(-c4ccccc4)cc3)N12 | 0 |
| KKL-1603 | CC(C)CCCCCCCCCC/C=C/C(=O)N[C@@H]1[C@@H](O)[C@@H](O)C(CC(O)[C@H]2O[C@@H](N3C=CC(=O)NC3=O)[C@H] [illegible] | 0 |
| KKL-1604 | O=C(/C=C/c1ccccc1)/C=C/c1ccccc1 | 0 |
| KKL-1605 | COC(=O)[C@@H]1Cc2c([nH]c3ccccc23)[C@@H]2C[C@H](NCCCn3ccnc3)C[C@@H](c3ccc(F)c(F)c3)N12 | 1 |
| KKL-1606 | COC(=O)[C@@H]1Cc2c([nH]c3ccccc23)[C@H]2C[C@H](NCCNC(C)=O)C[C@@H](c3ccc(C(C)C)cc3)N12 | 0 |
| KKL-1607 | C(C1CCN(c2nc3ccccc3n3cnnc23)CC1)c1ccccc1 | 0 |
| KKL-1608 | COC(=O)COc1ccc(-c2ccc(F)cc2)cc1C=O | 0 |
| KKL-1609 | CS(=O)(=O)Cc1nc2ccccc2oc1=O | 0 |
| KKL-1610 | COC(=O)[C@@H]1Cc2c([nH]c3ccccc23)[C@H]2C[C@H](NCc3cccnc3)C[C@@H](c3ccc(OC(F)(F)F)cc3)N12 | 0 |
| KKL-1611 | COc1cc(SC)ccc1C(=O)N[C@H]1CCN2[C@@H](C1)C(=O)N(c1ccc(Cl)cc1)C2=O | 0 |
| KKL-1612 | CC(C)CC(=O)OCC1=COC(OC(=O)CC(C)C)C2C1CC(OC(C)=O)C2(O)CCl | 0 |
| KKL-1613 | COC(=O)[C@@H]1Cc2c([nH]c3ccccc23)[C@H]2C[C@H](NCc3cccnc3)C[C@@H](c3ccc(C(F)(F)F)cc3)N12 | 0 |
| KKL-1614 | CC(C)CC(=O)OCC1=COC(OC(=O)CC(C)C)C2C1CC(OC(C)=O)C2(O)COC(=O)CC(C)C | 0 |
| KKL-1615 | CCOC(=O)Cc1csc(NC(=O)c2c(O)c3ccccc3n(CCC(C)C)c2=O)n1 | 75 |
| KKL-1616 | Cc1ccc(NC(=O)c2ccc([N+]([O-])=O)o2)cc1C | 4 |
| KKL-1617 | Cc1ccc(/N=C(\N)Nc2nc(C)cc(N3CCCCC3)n2)c(C)c1 | 0 |
| KKL-1618 | Cc1ccc(NC(=O)N2CCC(c3nc(O)c4nnn(Cc5cccc(Cl)c5)c4n3)CC2)cc1 | 0 |
| KKL-1619 | Cc1cc([N+]([O-])=O)ccc1NC(=O)CN1C2CCC1CC(NC(=O)NC13CC4CC(CC(C4) [illegible] | 0 |
| KKL-1620 | CC(C)OC(=O)CSc1nnc(-c2ccc(Cl)cc2Cl)o1 | 0 |
| KKL-1621 | Cn1cc(C(=O)CCl)c2ccccc12 | 1 |
| KKL-1622 | N(c1nc(-c2ccccn2)cs1)c1ccccc1 | 0 |
| KKL-1623 | Cn1c(COc2cccc3ccccc23)nnc1SCC(=O)OC1CCCCC1 | 1 |
| KKL-1624 | CN1C(=O)C(CC(C)=O)(OC(C)=O)c2ccccc21 | 0 |

| | | |
|---|---|---|
| KKL-1625 | CC1CC(C)(C)Nc2ccc(N=O)cc21 | 0 |
| KKL-1626 | [O-][N+](=O)c1cc(C(=O)NCc2ccco2)cc([N+]([O-])=O)c1 | 0 |
| KKL-1627 | COc1ccc(C(=O)Nc2cc(OC)c(NC(=O)c3ccco3)cc2OC)cc1 | 0 |
| KKL-1628 | Cc1ccc(Cc2cnc(NC(=O)c3cccs3)s2)cc1 | 0 |
| KKL-1629 | CCOC(=O)CSc1nc(N)c(C#N)c(CC(C)C)c1C#N | 0 |
| KKL-1630 | Cc1cccc(-c2cc3nc(C)c(C)c(NCCCn4ccnc4)n3n2)c1 | 0 |
| KKL-1631 | O=C(OCC#CCSc1nnc(-c2ccccc2)o1)c1ccco1 | 4 |
| KKL-1632 | Cc1ccc(NCc2nnc(SCC(=O)OC3CCCCC3)o2)c(C)c1 | 0 |
| KKL-1633 | CCOC(=O)CSc1nnc(-c2ccc([N+]([O-])=O)cc2)o1 | 0 |
| KKL-1634 | O=C(Oc1cccc2cccnc12)/C=C\c1ccccc1 | 0 |
| KKL-1635 | CCCC(=O)N1CCN(c2ccc(NC(=O)c3ccc([N+]([O-<br>])=O)o3)cc2Cl)CC1 | 0 |
| KKL-1636 | C[n+]1c2n(c3ccccc13)N=C(c1cc(C(C)(C)C)c(O)c(C(C)(C)C)c1)C<br>S2 | 0 |
| KKL-1637 | CCCCCCCCn1c2c(c(=N)c3ccccc13)CCC2 | 1 |
| KKL-1638 | CC(C)Nc1nc(NC(C)C)nc(-n2nc(C)cc2C)n1 | 0 |
| KKL-1639 | Clc1cc(Cl)c2cccnc2c1OC(=O)c1ccco1 | 0 |
| KKL-1640 | Cc1ccc([N+]([O-<br>])=O)cc1NC(=O)CN(C1CCCCC1)C(=O)c1ccccc1Cl | 0 |
| KKL-1641 | Cc1ccc(CNC(=O)c2ccc([N+]([O-])=O)o2)cc1 | 0 |
| KKL-1642 | Clc1cccc(NCc2nnc(SCC(=O)OC3CCCCC3)o2)c1 | 3 |
| KKL-1643 | CCN1/C(=C/c2sc3ccccc3[n+]2C)C=Cc2cccc(C)c21 | 0 |
| KKL-1644 | COc1ccc(CNC(=O)c2c(C)nc3ccccn23)cc1 | 0 |
| KKL-1645 | Clc1cccc(Cc2cnc(N3C(=N)SCC3=O)s2)c1 | 2 |
| KKL-1646 | CCOC(=O)CSc1ccc(-c2nc3ccccc3[nH]2)cn1 | 0 |
| KKL-1647 | Cc1nc2ccccn2c1C(=O)NCc1ccc2c(c1)OCO2 | 1 |
| KKL-1648 | COc1ccc(C(=O)Oc2cccc3cccnc23)cc1[N+]([O-])=O | 0 |
| KKL-1649 | OC(CN1CCN(c2ccccn2)CC1)Cn1c2ccc(Cl)cc2c2cc(Cl)ccc12 | 0 |
| KKL-1650 | COC(=O)CSc1nnc(-c2ccc(OC)cc2OC)o1 | 2 |
| KKL-1651 | [O-][N+](=O)c1ccc(C(=O)NCCN2CCOCC2)c([N+]([O-])=O)c1 | 0 |
| KKL-1652 | [O-][N+](=O)c1cc([N+]([O-])=O)c(Nc2ccccn2)c([N+]([O-])=O)c1 | 0 |
| KKL-1653 | COc1ccc(-c2nnc(SCC(=O)OC3CCCCC3)o2)cc1OC | 0 |

| | | |
|---|---|---|
| KKL-1654 | Cc1cc(C(=O)CCl)c(C)n1Cc1ccco1 | 0 |
| KKL-1655 | O=C(CSc1nnc(-c2cccs2)o1)OC1CCCCC1 | 0 |
| KKL-1656 | CCCC(=O)N1CCN(c2ccc(NC(=O)c3ccc([N+]([O-])=O)o3)cc2)CC1 | 0 |
| KKL-1657 | CC(C)(C)NC(=O)COC(=O)c1snc(C(=O)NC2CCCCC2)c1N | 1 |
| KKL-1658 | CCc1nnc(NC(=O)c2cc3ccccc3oc2=O)s1 | 0 |
| KKL-1659 | Cc1cccc(OCc2nnc(SCC(=O)OC3CCCCC3)o2)c1 | 0 |
| KKL-1660 | CSc1nc(C(Cl)Cl)nc(-c2ccccc2)n1 | 4 |
| KKL-1661 | COc1ccc(C2(CNC(=O)c3cc([N+]([O-])=O)cc([N+]([O-])=O)c3)CCCC2)cc1OC | 0 |
| KKL-1662 | COc1ccc(-c2n(C)c3ccccc3[n+]2CC(=O)OC2CC(C)CCC2C(C)C)cc1 | 0 |
| KKL-1663 | COc1cc(CNCc2ccccn2)ccc1OCc1ccc(Cl)cc1Cl | 0 |
| KKL-1664 | CCOC(=O)CSc1ccc(-c2nc3cc(C)c(C)cc3[nH]2)cn1 | 0 |
| KKL-1665 | COC(=O)CSc1nnc(-c2ccc(OC)c(OC)c2)o1 | 0 |
| KKL-1666 | CCOC(=O)CSc1nnc(-c2ccc(OC)cc2OC)o1 | 0 |
| KKL-1667 | CC(C)COC(=O)c1ccc2c(c1O)C(=O)c1c(cccc1)C2=O | 0 |
| KKL-1668 | Cc1nc(NC(=O)c2ccc(C)cc2)sc1C(=O)Nc1ccc2c(c1)CCC2 | 0 |
| KKL-1669 | C(SC1=CC=CC=C1)C1=NN2C(=NN=C2C2=CC=CC=C2)S1 | 2 |
| KKL-1670 | FC1=CC=CC(OC2=CC(C3CCCNC3)=NC(C3=CC=NC=C3)=N2)=C1 | 3 |
| KKL-1671 | CC(C)(C)C1=CC=C(C2=NN=C(SCC(=O)OC3CCCCC3)O2)C=C1 | 0 |
| KKL-1672 | BrC1=C(C2=CC=C3OCOC3=C2)N=C2N=CC=CN12 | 2 |
| KKL-1673 | CC1=CC=CC(CN2C=CC(NC(=O)C3=CC=C4OCOC4=C3)=N2)=C1 | 3 |
| KKL-1674 | C(C1=CC=CS1)C1=NN2C(=NN=C2C2=NNC3=C2CCC3)S1 | 0 |
| KKL-1675 | CC1=CC(OCCCON2C(=N)N=C(N)NC2(C)C)=CC(C)=C1 | 13 |
| KKL-1676 | CN1C=C(NC(=O)C2=CC(NC(=O)CNC(N)=N)=CN2C)C=C1C(=O)NCCC(N)=N | 0 |
| KKL-1677 | C1CC1C1=CC(NC2CCCCC2)=NC(C2=CC=CC=N2)=N1 | 11 |
| KKL-1678 | C12=C(N=C(NC3=CC=C(S(N)(=O)=O)C=C3)C=N1)C=CC=C2 | 4 |
| KKL-1679 | FC1=CC=C2N=C(NC3=NC(C4=CC=CC=N4)=CS3)SC2=C1 | 2 |
| KKL-1680 | COC1=C2OCOC2=CC(CN2CCC3(CC2)C=CC2=C3C=CC=C2)=C1 | 4 |
| KKL-1681 | COC1=CC=C2SC(NC3=NC(C4=NC=CC=C4)=CS3)=NC2=C1 | 0 |
| KKL-1682 | COC1=CC=C(C2=CN=CC(C3=CC(N4CCN(CCO)CC4)=NC(C4=CC=CC=C4)=N3)=C2)C=C1 | 0 |

| | | |
|---|---|---|
| KKL-1683 | CSC1=CC=CC(C(=O)NC2=NC(C3=CC=CC=N3)=CS2)=C1 | 10 |
| KKL-1684 | CC1=C2C=CC=CC2=C(NCC2=CC=CC(Cl)=C2)N=N1 | 0 |
| KKL-1685 | OC1=CC=C(F)C=C1C(=O)C1=CN(C(=O)C2=CC=CO2)N=C1 | 3 |
| KKL-1686 | CC1=CC=C(C(=O)NC2=NC(C3=CC=CC=N3)=CS2)S1 | 12 |
| KKL-1687 | FC1=C(C2=NOC(N3CCN(CC4=CC=C5OCOC5=C4)CC3)=N2)C=CC=C1 | 0 |
| KKL-1688 | FC1=CC=C(CC(=O)NC2=NC(C3=NC=CC=C3)=CS2)C=C1 | 13 |
| KKL-1689 | C(CC1=CC=CC=C1)NC1=CC(C2=CC=CC=C2)=NC2=NN=CN12 | 2 |
| KKL-1690 | CC(OC1=CC=CC=C1)C(=O)NC1=NC(C2=CC=CC=N2)=CS1 | 15 |
| KKL-1691 | CCN1C(=O)C(=O)NC2=C1C=CC(C(=O)NC1=C(NC3=CC=C(OC)C=C3)C=CC=C1)=C2 | 2 |
| KKL-1692 | CCC1=CC=C([C@@H]2C[C@H](C(F)(F)F)N3N=CC(C(=O)NCC4=CC=C(OC)C=C4)=C3N2)C=C1 | 0 |
| KKL-1693 | C1(N)=C(NC2=CC=C(OC)C=C2)N=CC=C1 | 4 |
| KKL-1694 | C12=C(C=C(OC3CCNCC3)C(C3=CC=C(OC(F)(F)F)C=C3)=C1)C=NC=C2 | 8 |
| KKL-1695 | CC1=CC=CC(C2=NC(NC3=CC=NC=N3)=C3SC=CC3=N2)=N1 | 0 |
| KKL-1696 | O=C(N1CCN(CC2=CC=C3OCOC3=C2)CC1)C1=CC2=CC=CC=C2O1 | 0 |
| KKL-1697 | C1OC2=C(O1)C=C1N=C(NC3=NC(C4=CC=CC=N4)=CS3)SC1=C2 | 0 |
| KKL-1698 | CC1=CC=CC=C1OCCCON1C(=N)N=C(N)NC1(C)C | 11 |
| KKL-1699 | O=C(C=CC1=CC=CC=C1)N1CCN(CC2=CC=C3OCOC3=C2)CC1 | 1 |
| KKL-1700 | CCC1=CC=CC=C1OCCCON1C(=N)N=C(N)NC1(C)C | 3 |
| KKL-1701 | O=C(CCC1CCCCC1)NC1=CC=CC(C2=NN=C(C3=CC=CO3)O2)=C1 | 3 |
| KKL-1702 | COC1=CC=C(NC2=CC=CC=C2NC(=O)C2=CC=C(O)N=C2)C=C1 | 2 |
| KKL-1703 | FC1=CC=C2C(Cl)=C(C(=O)NC3=NCCS3)SC2=C1 | 0 |
| KKL-1704 | O=C(N1CCN(CC2=CC=C3OCOC3=C2)CC1)C1=C2CCCCC2=NN1 | 0 |
| KKL-1705 | ClC1=CC=C(S(=O)(=O)C2=C(C#N)N=C(C3=CC=CC=C3)O2)C=C1 | 0 |
| KKL-1706 | CSC1=CC=C(C2=NN=C(NC(=O)C3=CC=C(Cl)S3)O2)C=C1 | 50 |
| KKL-1707 | ClC1=C2N=C(N(CCCN3C=CN=C3)C(=O)C3=CC=CO3)SC2=CC=C1 | 0 |
| KKL-1708 | CCC1=C(C)N=C(N2N=C(C)C=C2NC(=O)C2=CC=CC=C2SC)NC1=O | 2 |
| KKL-1709 | O=C(CCCSC1=CC=CC=C1)NNC(=O)C1=CC=CS1 | 3 |
| KKL-1710 | NC1=NC(N)=C2C(SC3=CC=C4C=CC=CC4=C3)=CC=CC2=N1 | 0 |
| KKL-1711 | NC1=CC=CN=C1NC1=CC=C(C(F)(F)F)C=C1 | 4 |

| | | |
|---|---|---|
| KKL-1712 | CC1=CSC2=NC(C3=CC=CC(C)=N3)=NC(NC3=CC=NC=N3)=C12 | 6 |
| KKL-1713 | COC1=CC(NC2=CC=CC=C2)=CC=C1NS(=O)(=O)C1=C(C)ON=C1C | 6 |
| KKL-1714 | CCN(CC)C1=CC=C(C2=NN3C(=NN=C3C3=NNC4=C3CCC4)S2)C=C1 | 1 |
| KKL-1715 | CNC1=C2SC=CC2=NC(SC)=N1 | 5 |
| KKL-1716 | C[C@@H](NCCC1=NC(C2=CN(C)C3=CC=CC=C23)=NO1)C1CCCCC1 | 3 |
| KKL-1717 | CC1(C)N=C(N)N=C(N)N1OCCC1=CC=C2C=CC=CC2=C1 | 20 |
| KKL-1718 | NC1=CC(OCC2=CC=CC=C2)=CC=C1NC1=CC=CC=C1 | 0 |
| KKL-1719 | C1=C(F)C(S(N(C2CC2)CC2=CC=C(C(N)=O)C=C2)(=O)=O)=CC=C1 | 0 |
| KKL-1720 | N1(C(NCC2=COC=C2)=O)CCN(CC2=CC=C3C(=C2)OCO3)CC1 | 3 |
| KKL-1721 | C1=CC=CC(S(C(C2=CC=CS2)CN)(=O)=O)=C1 | 0 |
| KKL-1722 | COC1=C(OCCCCN2C(C3CC3)=NC3=CC=CC=C23)C=CC=C1 | 5 |
| KKL-1723 | CC(C)OC(=O)COC1=CC=C(C2=CN=C(N(C)CCC3=CC=NC=C3)N=C2)C=C1 | 3 |
| KKL-1724 | CC1=CN=C(N2CCN[C@@H](CC3=CC=CC=C3)C2)N=C1OC1=CC=C(N2C=CN=C2)C=C1 | 6 |
| KKL-1725 | CC1=C(C)C=C(CN2CCC3(CC2)OCCC2=C3C=CS2)O1 | 0 |
| KKL-1726 | N1(C(NCC2=CC=CO2)=O)CCN(CC2=CC=C3C(=C2)CCO3)CC1 | 0 |
| KKL-1727 | C1N(C(NCC2=CC=CO2)=O)CCN(CC2=CC=C(Cl)S2)C1 | 3 |
| KKL-1728 | C12=C(N(CCNC(=O)C3=CC=CC=N3)C(C(F)(F)F)=CC1=O)C=CC(C)=C2 | 0 |
| KKL-1729 | O=C(NC1=CC=C(NC2=NCCN2)C=C1)C1=CC=C(C(=O)NC2=CC=C(NC3=NCCN3)C=C2)C=C1 | 0 |
| KKL-1730 | C(CN1CCOCC1)N1C(CC2=CC=CC=C2)=NN=C1SC1=CC=NC(N2CCN(C3=CC=NC=C3)CC2)=N1 | 0 |
| KKL-1731 | C(CN1C(SC2=CC=NC(N3CCN(C4=CC=NC=C4)CC3)=N2)=NN=C1C1=CC=CC=N1)CN1CCOCC1 | 4 |
| KKL-1732 | C12=C(C(=O)C(CNCCCC)CO1)C=C(OC)C=C2 | 0 |
| KKL-1733 | C12=C(C)C(C(=O)NCCOC3=CC=CC=C3)N=C1CCCC2=O | 0 |
| KKL-1734 | C12=C(C=C(C(=O)NC)C(OCC)=C1)C=CC=C2 | 0 |
| KKL-1735 | CC1=C(C(=O)NCC2=CC=CC=C2)N2C=CC=CC2=N1 | 0 |
| KKL-1736 | C1=CC=CC(C(NC2=NC3=C(S2)CCCC2=C3N=CC=C2)=O)=C1C(F)(F)F | 26 |
| KKL-1737 | C1(C2=CC(F)=C(OC3=NN=C(C(OC)=O)C=C3)C=C2)=CSC(C)=N1 | 3 |
| KKL-1738 | C12=C(C(Cl)=NN=C1N1CCCC1)C=CC=C2 | 0 |
| KKL-1739 | C([C@H]1CN(C2=NC=CC(OC3=CC=C(N4C=CN=C4)C=C3)=N2)CCN1)C1=CC=CC=C1 | 4 |
| KKL-1740 | CC(C)(C)C(=O)CC1(O)OC2=CC=CC=C2N=C1C=C(O)C(C)(C)C | 6 |

| | | |
|---|---|---|
| KKL-1741 | CC1=CC=C2N=C(Cl)C(CNCC3=CC=CS3)=CC2=C1 | 0 |
| KKL-1742 | ClC1=CC=C(SCCC(=O)NC2=NC(C3=CC=CC=N3)=CS2)C=C1 | 10 |
| KKL-1743 | CC1(C)NC(N)=NC(=N)N1OCCCOC1=CC=C(Cl)C=C1Cl | 18 |
| KKL-1744 | CSC1=NC(NCC=C)=C2C=CC=CC2=N1 | 5 |
| KKL-1745 | CSC1=CC=C(C(=O)NC2=NC(C3=NC=CC=C3)=CS2)C=C1 | 5 |
| KKL-1746 | COC1=CC=C(CCNC(=O)C2=C(C)N=C3C=CC=CN23)C=C1 | 14 |
| KKL-1747 | CN(C)S(=O)(=O)C1=CC=C(C(=O)NC2=NN=C(C3=CC=C(Cl)S3)O2)C=C1 | 1 |
| KKL-1748 | CC1=C(C(=O)NCC2=CC=C(Cl)C=C2)N2C=CC=CC2=N1 | 13 |
| KKL-1749 | C12=C(OC(=O)C(CCC(N(CC3=CC=C(Cl)C=C3)C)=O)=N1)C=CC=C2 | 0 |
| KKL-1750 | COC1=CC=C(C2=CC=C(NC3=CC=CC(S(=O)(=O)CCNCCC4=CC=CS4)=C3)N=C2)C=N1 | 0 |
| KKL-1751 | ClC1=CC=C(CCNC2=CC(C3=CC=CC=C3)=NC3=NC=NN23)C=C1 | 0 |
| KKL-1752 | O=C(NC1=CC=CC=C1C1=NC(C2CCC2)=CS1)OCC1CCNCC1 | 2 |
| KKL-1753 | COC1=CC(CC2=CN=C(N)N=C2N)=C2C=NC=CC2=C1N(C)C | 0 |
| KKL-1754 | C1=C(C2=NN=C(SC3=NN4C(=NN=C4C(F)(F)F)C=C3)O2)OC=C1 | 1 |
| KKL-1755 | COC1=CC=CC(C2=NC3=C(N4CCCC4)C=CC=C3N2CC2CCCN2)=N1 | 0 |
| KKL-1756 | CC1=C(C(=O)NCC2=CC=CO2)N2C=C(C)C=CC2=N1 | 0 |
| KKL-1757 | CC(C)CNC(=O)C1=C(C)N=C2C=CC(C)=CN12 | 3 |
| KKL-1758 | C1(Cl)=C(Cl)C(C2=C(NC)N=C(SC)N=N2)=CC=C1 | 0 |
| KKL-1759 | O=C(C=CC1=CC=CS1)N1CCN(CC2=CC3=C(OCO3)C=C2)CC1 | 0 |
| KKL-1760 | CCOC(=O)C1=NOC(C2=CSC(NC3=C(C)C=C(C)C=C3C)=N2)=C1 | 0 |
| KKL-1761 | CC1=CC=CC=C1CN1CCC(O)(C2=CC=C(Cl)C(C(F)(F)F)=C2)CC1 | 3 |
| KKL-1762 | ClC1=CC=C(C(=O)NC2=NN=C(C3=CC=C(Cl)S3)O2)S1 | 0 |
| KKL-1763 | CCCCC1=CC=C(C(=O)NC2=C3C=CC=CC3=CC=N2)N=C1 | 50 |
| KKL-1764 | NC(=NC1=C2C=CC=CC2=CC(C2=CC=CC=N2)=N1)C1=CC=CC(Cl)=C1 | 0 |
| KKL-1765 | COC1=CC=C2N=C(C)C=C(OCC(=O)NCC3=CC=CC=C3)C2=C1 | 39 |
| KKL-1766 | CC1=CC=C(CNC(=O)C2=C3NC(C4=CC=C(C)C=C4)CC(C(F)F)N3N=C2)C=C1 | 0 |
| KKL-1767 | CC(C)N1N=CC2=C(C(=O)NCC(N3CCOCC3)C3=CC=CS3)C=C(C3=CC=CC=C3)N=C12 | 0 |
| KKL-1768 | CNC1=C2C=CC=CC2=NC(SCC(=O)N(C)CC2=CC=C(Cl)S2)=N1 | 0 |
| KKL-1769 | FC(F)(F)C1=C(C(=O)NCC2=CC=C3OCOC3=C2)N2C=CC=CC2=N1 | 0 |

| | | |
|---|---|---|
| KKL-1770 | CC1=C(C(=O)NC2=CC=C(C(C)(C)C)C=C2)N2C=CC=CC2=N1 | 0 |
| KKL-1771 | CC1=CN2C(=NC(C(F)(F)F)=C2C(=O)NCC2=CC=C3OCOC3=C2)C=C1 | 2 |
| KKL-1772 | CC(C)OC1=CC=CC(CNC(=O)C2=C(C(F)(F)F)N=C3C=CC(C)=CN23)=C1 | 2 |
| KKL-1773 | CC(C)C1=CC=C(C2=CN=CC(N3CCN(C4=CC=NC=C4)CC3)=N2)C=C1 | 3 |
| KKL-1775 | CCC1=CC=C(CN2CCC3(CC2)OCCC2=C3C=CS2)S1 | 2 |
| KKL-1776 | C1(C2=NC=C(C3=CC=C(OCCCN4C=CN=C4)C=C3)S2)=CC=CC=C1 | 0 |
| KKL-1777 | C1=CC=C(NC2=NC3=C(S2)CCN(CC2=CC=C(C4C(C)C4)O2)CC3)C=C1 | 0 |
| KKL-1778 | C1(OC)=CC=C(CN2CCN(C(=O)CCC3=CSC(C)=N3)CC2)C=C1 | 0 |
| KKL-1779 | CCC1=CC=C(C2CC(C(F)(F)F)N3N=CC(C(=O)NCC4=CC=CS4)=C3N2)C=C1 | 0 |
| KKL-1780 | CC(=O)N1CCC2=C1C=C1C(C)=CC(C)=NC1=C2 | 0 |
| KKL-1781 | ClC1=CC=C2NC(C(=O)NC34CC5CC(CC(C5)C3)C4)=CC2=C1 | 0 |
| KKL-1782 | C12=C(OC(=O)C(CCC(O)=O)=N1)C=CC=C2 | 0 |
| KKL-1783 | N1=C2N(C(C3=CC=C(OC)C=C3)=N1)N=C(CC1=CSC(C)=N1)S2 | 0 |
| KKL-1784 | BrC1=CN=CC(C2=CC(NCCCN3C=CN=C3)=NC(C3=CC=CC=C3)=N2)=C1 | 0 |
| KKL-1785 | CC1=CN2C(=NC(C(F)(F)F)=C2C(=O)NCC2=CC(Cl)=CC(Cl)=C2)C=C1 | 16 |
| KKL-1786 | CC1=CC(NCCC2=CC=CC=C2)=NC(NC(=N)NC2=CC=C(Cl)C=C2)=N1 | 3 |
| KKL-1787 | FC1=CC=CC(OCC(=O)NC2=NC(C3=CC=CC=N3)=CS2)=C1 | 5 |
| KKL-1788 | CCCC1=CC(C(C)=O)=CC=C1OCCCON1C(=N)N=C(N)NC1(C)C | 0 |
| KKL-1789 | COC1=CC=CC=C1C1=CC=CC(CNC(=O)C2=C(OC)C=C3C=CC=CC3=C2)=C1 | 6 |
| KKL-1790 | CCOC1=CC=C(OCC2=CC=C(C(=O)NNC(=O)C3=CC=C(N(=O)=O)O3)C=C2)C=C1 | 0 |
| KKL-1791 | CCC1=CC=C(NC2=NN=C(SCC3=NOC(C)=C3)S2)C=C1 | 0 |
| KKL-1792 | CC1=CN2C(=NC(C(F)(F)F)=C2C(=O)NCC2=CC=CO2)C=C1 | 0 |
| KKL-1793 | ClC1=CN=C(NC2=NC(C3=NC=CC=C3)=CS2)C=C1 | 0 |
| KKL-1794 | C1(N2CCN(C3CS(=O)(=O)CC3O)CC2)=C(Cl)C=CC=C1 | 6 |
| KKL-1795 | CN1CCN(C2=NC(C3=CC=CC=C3)=NC(C3=CC=NC=C3)=C2)CC1 | 237 |
| KKL-1796 | FC1=CC=CC=C1C1=NC(C#N)=C(NCCC2=CC=CC=C2)O1 | 0 |
| KKL-1798 | C(CN1C(SC2=CC=NC(N3CCN(C4=CC=NC=C4)CC3)=N2)=NN=C1C1=CC=CS1)CN1CCOCC1 | 0 |
| KKL-1799 | COC1=CC=C(C2=NSC(SCC(=O)N3CCCC3)=N2)C=C1 | 1 |
| KKL-1800 | CCCC1=CC=C(S(=O)(=O)NC2=CC=C(CN3CCN(CC4CC4)CC3)C=C2)C=C1 | 3 |

| | | |
|---|---|---|
| KKL-1801 | CC1=C(C(=O)NCC2=CC=CS2)N2C=CC=CC2=N1 | 0 |
| KKL-1802 | CCCCS(=O)(=O)NC1=CC=C2C=NN(C)C2=C1 | 0 |
| KKL-1803 | CCOC1=CC=C(C2=NC(CNCCC3=CC=C(F)C=C3)=CS2)C=C1 | 0 |
| KKL-1804 | CCN(CC1=CC=C2OCOC2=C1)C(=O)CSC1=NC(NC)=C2C=CC=CC2=N1 | 1 |
| KKL-1805 | COC1=CC=C(CC(=O)NC2=C(C3=CC=CS3)N=C3C=C(C)C=CN23)C=C1 | 0 |
| KKL-1806 | CC1=C(C(=O)NCCSC2=CC=C(C)C=C2)C(C2=CC=CC=C2)=NO1 | 1 |
| KKL-1807 | FC(F)(F)C1=C(C(=O)NCC2=CC(Cl)=CC(Cl)=C2)N2C=CC=CC2=N1 | 0 |
| KKL-1808 | FC1=CC=CC(Cl)=C1COC(=O)C1=CC=C(N2N=CC(Cl)=C(Cl)C2=O)C=C1 | 4 |
| KKL-1809 | COC1=CC=C(C2=NC(C3=CC=CC(NC(=O)CC4=CC=C(F)C=C4)=C3)=NO2)C=C1OC | 4 |
| KKL-1810 | C1=C(C2=CN=C(OC(C)C3CCN(C4=NC(C(C)C)=NO4)CC3)C=N2)C=CC(S(C)(=O)=O)=C1F | 0 |
| KKL-1812 | C1=CC=CC(NC2=CC=CC=C2)=C1NCC(N(C1=CC=CC=C1)C(C)C)=O | 1 |
| KKL-1813 | C12=C(C=C(OC3CCNCC3)C(/C=C/CCCC)=C1)C=NC=C2 | 3 |
| KKL-1814 | C12=C(N=C(OC)C=C1)C(NC(=O)C1CCN(CCC3=CC=C(Cl)C=C3)CC1)=CC=N2 | 1 |
| KKL-1815 | C(N1CCC2(CC1)OCCC1=C2C=CS1)C1=CC=C2OCCOC2=C1 | 0 |
| KKL-1816 | CC1=CC(OCC(=O)NC2=CC=CC=C2)=C2C=C(Br)C=CC2=N1 | 9 |
| KKL-1817 | CN1C2=NSC(S(C)=O)=C2C(=O)N(C)C1=O | 0 |
| KKL-1818 | COC1=CC=C2C=C(C(=O)NC3=CC=C(C4CCN(C)CC4)C=C3)SC2=C1 | 1 |
| KKL-1819 | FC(F)(F)C1=CC=C2C(SCCCN3CCC4=CC=C(C#N)C=C4CC3)=CC=NC2=C1 | 0 |
| KKL-1820 | CC1=CC=C2NC(C3CCCNC3)=C(C3=CC=NC=C3)C2=C1 | 14 |
| KKL-1821 | O=C(C=CC1=CC=CO1)N1CCN(CC2=CC=C3OCOC3=C2)CC1 | 0 |
| KKL-1822 | CCC1=NC2=CC=CC=C2N1CC(O)COC1=CC(C)=CC=C1Cl | 0 |
| KKL-1823 | FC(F)(F)C1=NC(C2=CC=CC=N2)=NC(SCC2=CC=CC=C2)=C1 | 6 |
| KKL-1824 | CCC(SC1=NC2=CC=CC=C2O1)C(=O)NC1=NC=CS1 | 3 |
| KKL-1825 | CC1=CC=C(C2CC(C(F)(F)F)N3N=CC(C(=O)NCC4=CC5=C(OCO5)C=C4)=C3N2)C=C1 | 0 |
| KKL-1826 | NC(=O)C1=CC(C2=CSC=C2)=CC=C1OCCOC1=CC=CC=C1 | 0 |
| KKL-1827 | C1(C(F)(F)F)=CC(C#N)=CN2C1=NC(C(=O)NCC1=CC=CS1)=C2Cl | 1 |
| KKL-1828 | C1N(C2=NC=C(NC(NCCC3=CN(C)N=C3)=O)C=C2)CCCC1 | 0 |
| KKL-1829 | FC1=CC=CC=C1CNC(=O)N1CCN(CC2=CC=C(Br)S2)CC1 | 1 |
| KKL-1830 | O=C(NC1=CC=CC=C1N1CCOCC1)C(CC1=CC=CC=C1)NC(=O)C1=CC=CS1 | 2 |

| | | |
|---|---|---|
| KKL-1831 | COC1=CC=C(CCNC2=CC(C3=CC=CC=C3)=NC3=NC=NN23)C=C1 | 3 |
| KKL-1832 | CC1=CC(OCC(=O)NCC2=CC=CC=C2)=C2C=C(Br)C=CC2=N1 | 6 |
| KKL-1833 | COC1=C2C(N)=C(C)C=CC2=NC(C(F)(F)F)=C1 | 7 |
| KKL-1834 | CC1=NN(CC2=CC=CC=C2)C(Cl)=C1C(=O)NC1CCCCCC1 | 1 |
| KKL-1835 | C1=CC(C(C2=CC=CS2)=O)=CC2=C1CC(C(=O)N)S2 | 0 |
| KKL-1836 | CN1CCN(C2=CC=C(C(F)(F)F)C=C2NC(=O)C2=CC=CC=N2)CC1 | 7 |
| KKL-1837 | COC1=CC=C(C2=CC(NC(=O)C3=CC=C(F)C=C3Cl)=CC=C2OC)C=C1 | 0 |
| KKL-1838 | O=C(C1CCN(C2=NN=C(N3C=CC=C3)S2)CC1)N1CCC(CC2=CC=CC=C2)CC1 | 1 |
| KKL-1839 | CC1=C(S(=O)(=O)NCC2=NC(NCCCC3=CC=CC=C3)=C3C=CC=CC3=N2)C(C)=NN1 | 1 |
| KKL-1840 | C1=C2C(=CC(C3=NN=C(C4=CC(Cl)=C(OC(C)C)C=C4)S3)=C1)CCNCC2 | 4 |
| KKL-1841 | C1(NC(=O)N2CCN(CC3=CC=CS3)CC2)=CC=CO1 | 1 |
| KKL-1842 | C1=CC(CC2CCN(C(=O)NC3=NC(C4=CC=CC=N4)=CS3)CC2)=CC=C1F | 28 |
| KKL-1843 | COC1=CC=C2N=C(C)C=C(OCC(=O)NC3=CC=CC=C3OC)C2=C1 | 3 |
| KKL-1844 | CN1CCN(C2=NC(C3=CC=NC=C3)=NC(C3=CC=C(F)C=C3)=C2)CC1 | 0 |
| KKL-1845 | CSC1=CC=CC=C1C(=O)NC1=NC(C2=CC=CC=N2)=CS1 | 8 |
| KKL-1846 | C1(N)=C(NC2=CC=C(Cl)C=C2)C=CC=C1 | 3 |
| KKL-1847 | CC(C)(C)C1=CC=CC=C1OCCCON1C(=N)N=C(N)NC1(C)C | 15 |
| KKL-1848 | NC(=N)C1=CC=C(CCN2CCC(C3=NC(COCC(F)(F)F)=C(C4=CC=C(F)C=C4)O3)CC2)C=C1 | 28 |
| KKL-1849 | CCC1=CC=C(S(=O)(=O)NC2=CC(CN3CCN(CCC(C)C)CC3)=CC=C2C)C=C1 | 4 |
| KKL-1850 | CN1CCC[C@H]1COC1=C(C)C=C(/C=C/C2=CC=CC(C(O)=O)=C2)C=C1Cl | 8 |
| KKL-1851 | O=C(N1CCN(CC2=CC=C3OCOC3=C2)CC1)C1=CC2=C(CCC2)S1 | 0 |
| KKL-1852 | FC(F)(F)C1=C(C(=O)NCC2=CC=C(OC3=CC=CC=C3)C=C2)N2C=CC=CC2=N1 | 3 |
| KKL-1853 | C1=CC(C(NC2=NN=C(S(CCCC)(=O)=O)S2)=O)=NC(CCCC)=C1 | 6 |
| KKL-1854 | N1(COCC)C(=O)N=C(NS(=O)(=O)C2=CC=C(N)C=C2)C=C1 | 0 |
| KKL-1855 | C1=CC(C#N)=CC(C2=CN=C(NCC3CCC(CN(C4=CC=CN=N4)C(=O)O)CC3)C=C2)=C1 | 0 |
| KKL-1856 | C1C(N2N=NN=C2)(C(NCCC2=CC=C(OC)C=C2)=O)CCCC1 | 3 |
| KKL-1857 | NC1=CC=CC=C1NC1=CC=C2OCOC2=C1 | 2 |
| KKL-1858 | NC1=CC=CN=C1NC1=CC=C(Cl)C=C1 | 0 |
| KKL-1860 | CN1C=CN=C1CNCCS(=O)(=O)C1=CC=CC(NC2=NC=C(C3=CC(F)=CC=C3)C=C2)=C1 | 2 |

| | | |
|---|---|---|
| KKL-1861 | BrC1=CC=C(CN2CCN(C(=O)C=CC3=CC=CS3)CC2)S1 | 0 |
| KKL-1862 | CC1=CN2C(=NC(C(F)(F)F)=C2C(=O)NCC2=CC=CS2)C=C1 | 0 |
| KKL-1863 | CC1=C(C(=O)NCC2=CC=CS2)N2C=C(C)C=CC2=N1 | 4 |
| KKL-1864 | COC1=CC=C(C2=CC(=O)C3=CC(OCC(=O)OC(C)C)=CC=C3O2)C=C1 | 2 |
| KKL-1865 | C1=CC(CC2=CN=C3C(=C2)NC(=O)C(C(=O)NCCO)=C3O)=C(C(F)(F)F)C=C1F | 4 |
| KKL-1866 | C1=C(CNS(C2=CC=C(C3=C4C(=CC=C3OC3CCNCC3)C=NC=C4)S2)(=O)=O)C=CC=C1 | 33 |
| KKL-1867 | O=C(COC1=CC=CC=C1)N1CCN(C2=NC3=CC=CC=C3N=C2)CC1 | 22 |
| KKL-1868 | FC(F)(F)C1=NN=C2C=CC(Cl)=NN12 | 0 |
| KKL-1869 | CNC1=C2C=CC=CC2=NC(SC)=N1 | 0 |
| KKL-1870 | OC1(C2=CC=CC=C2)OC2=CC=CC=C2N=C1C1=CC=CC=C1 | 2 |
| KKL-1871 | O=C(NC1=CC=CC=C1N1CCOCC1)[C@H](CC1=CC=CC=C1)NC(=O)C1=CC=CS1 | 4 |
| KKL-1872 | N1=C2C(=CC=C1C)OC(SC(=O)NC1=NC=CS1)=N2 | 0 |
| KKL-1873 | CC1=C(C2=NN(CC3=CC=CC=C3)C=C2)N2C=C(C)C=CC2=N1 | 3 |
| KKL-1874 | C1(OCCNC(=O)C2=C(NC)SN=C2C)=CC=C(OC)C=C1 | 0 |
| KKL-1875 | CC1=C(C(=O)NC2=C3SC=CC3=NC(C3=CC=CC=N3)=N2)C=CO1 | 11 |
| KKL-1876 | C1(N2C(NS(C3=CC=C(N)C=C3)(=O)=O)=CC=N2)=CC=CC=C1 | 2 |
| KKL-1878 | CC1=CC=C(CNC(=O)C2=C(C(F)(F)F)N=C3C=CC(C)=CN23)O1 | 2 |
| KKL-1879 | C1(C2=NN=C(CC3=CC=C(F)C=C3)N2)=NC(NC2=CC=C(OCCN3CCOCC3)C=C2)=NC=C1 | 2 |
| KKL-1880 | CCCOC(=O)COC1=CC=C2C(=O)C(C3=CC=CC=C3OC)=COC2=C1 | 4 |
| KKL-1881 | O=C(NC1=NC(C2=NC=CC=C2)=CS1)C1=CN=CN1C1=CC=CC=C1 | 5 |
| KKL-1882 | O=C(NNC(=O)C1=CC=C2C=CC=CC2=C1)C1=CC=C(N(=O)=O)O1 | 0 |
| KKL-1883 | C1(NC2=CC=C(OC)C(CC(=O)C3=CC=CC=C3)=C2)=C2C(=NC(NC3CCNC3)=N1)C=CNC2=O | 335 |
| KKL-1884 | C1=C2C(=CC=C1)NC(C1=NC=C(N(=O)=O)S1)=N2 | 0 |
| KKL-1885 | C1(C(O)CNCCC2=CC=CC=C2)=CC=C(N)C(O)=C1 | 3 |
| KKL-1886 | C1=CC(N2C=NC(C#N)=N2)=NN2C1=NN=C2C(F)(F)F | 0 |

[1]% average; average % luminescence normalized to that of KKL-35

#### Table S2. MIC for KKL-1005 and KKL-2108

| Compound | *M. tuberculosis* | *E. coli* Δ*tolC* | *B. anthracis* |
|---|---|---|---|
| | MIC[a] | MIC | MIC |
| KKL-1005 | 4.7 | 9.3 | 9.3 |
| KKL-2108 | 6.0 | 9.3 | 5.5 |

[a] MIC; µg/mL values from at least three broth microdilution assays.

#### Table S3. Mass spectrometry analysis of eluted gel fragment from Fig. 3C

| 50S ribosomal protein L7/L12 OS = *Bacillus anthracis* | | | | | | | | |
|---|---|---|---|---|---|---|---|---|
| **Score** | **Coverage** | **# Proteins** | **# Unique Peptides** | **# Peptides** | **# PSMs** | | | |
| 1333.77 | 55.46 | 1 | 5 | 5 | 264 | | | |
| **Sequence** | **# PSMs** | **Modifications** | **XCorr** | **Charge** | **MH+ [Da]** | **ΔM [ppm]** | **RT [min]** | **# Missed Cleavages** |
| AIEEEFGVTAAAPVAVAGGAGEAAAEK | 227 | | 6.84 | 3 | 2486.2452 | 1.66 | 84.65 | 0 |
| TEFDVELTSAGAQK | 7 | | 5.08 | 2 | 1495.7263 | -0.79 | 30.35 | 0 |
| SmTVLELNDLVK | 11 | M2 (Oxidation) | 4.17 | 2 | 1377.7283 | -0.83 | 37.28 | 0 |
| LEEVGAAVEVK | 15 | | 4.09 | 2 | 1143.6267 | 0.97 | 25.11 | 0 |
| SMTVLELNDLVK | 3 | | 4.00 | 2 | 1361.7369 | 1.79 | 40.74 | 0 |
| AKLEEVGAAVEVK | 1 | | 3.70 | 2 | 1342.7578 | 0.09 | 24.47 | 1 |

#### Table S4. Bacterial strains, plasmids, and oligonucleotides

| Strain | Description | Source or reference |
|---|---|---|
| *E. coli* DH5α | For propagation of plasmids | (Chan et al. 2013) |
| *E. coli* BL21 (DE3) | For protein expression | (Studier and Moffatt 1986) |
| *E. coli* MRE600 | For *E. coli* ribosome purification | (Cammack and Wade 1965) |
| *E. coli* Δ*tolC* | Efflux deficient *E. coli* MG1655 K12 mutant (*tolC*::kan) | Keio collection (Baba et al., 2006) |
| *E. coli* Δ*tolC* Δ*ssrA* | *E. coli* Δ*tolC ssrA*::cat | (Ramadoss et al. 2013) |

| | | |
|---|---|---|
| *B. anthracis* Sterne | pXO1⁺, pXO2⁻ | (Alumasa et al. 2017) |
| *M. tuberculosis* H37Rv Δ*RD1* Δ*panCD* | H37Rv derivative; Δ*RD1* Δ*panCD*; avirulent | Gift from Jacobs Lab (Sambandamurthy et al. 2006) |
| *E. coli* pLC1021 | *E. coli* MG1655 K12 expressing luc-trpAt | (Ramadoss et al. 2013) |
| *E. coli* pLC1063 | *E. coli* MG1655 K12 strain expressing mCherry-trpAt | (Ramadoss et al. 2013) |
| *E. coli* Δ*ssrA* pLC1063 | *E. coli* MG1655 K12 Δ*ssrA* expressing mCherry-trpAt | (Ramadoss et al. 2013) |
| *E. coli prplL* | *E. coli* MG1655 K12 *expressing* 6His-tagged *E.coli rplL* | (Ramadoss et al. 2013) |
| *E. coli* Δ*tolC* p*rplL* | *E. coli* Δ*tolC expressing* 6His-tagged *E.coli rplI* | This study |
| *E. coli* BL21(DE3) pET28a*infA*-$His_6$ | Contains plasmid expressing 6His-tagged *M. tuberculosis infA* | This study |
| *E. coli* BL21(DE3) pET28a*infB*-$His_6$ | Contains plasmid expressing 6His-tagged *M. tuberculosis infB* | This study |
| *E. coli* BL21(DE3) pET28a*infC*-$His_6$ | Contains plasmid expressing 6His-tagged *M. tuberculosis infC* | This study |
| *E. coli* BL21(DE3) pET28*fusA1*-$His_6$ | Contains plasmid expressing 6His-tagged *M. tuberculosis fusA1* | This study |
| *E. coli* BL21(DE3) pET28*fusA2*-$His_6$ | Contains plasmid expressing 6His-tagged *M. tuberculosis fusA2* | This study |
| *E. coli* BL21(DE3) pET28a*tsf*-$His_6$ | Contains plasmid expressing 6His-tagged *M. tuberculosis tsf* | This study |
| *E. coli* BL21(DE3) pET28a*tuf*-$His_6$ | Contains plasmid expressing 6His-tagged *M. tuberculosis tuf* | This study |
| *E. coli* BL21(DE3) pET28a*prfA*-$His_6$ | Contains plasmid expressing 6His-tagged *M. tuberculosis prfA* | This study |
| *E. coli* BL21(DE3) pET28a*prfB*-$His_6$ | Contains plasmid expressing 6His-tagged *M. tuberculosis prfB* | This study |
| *E. coli* BL21(DE3) pET28a*frr*-$His_6$ | Contains plasmid expressing 6His-tagged *M. tuberculosis frr* | This study |
| *E. coli* BL21(DE3) pET28aTB*rplL*-$His_6$ | Contains plasmid expressing 6His-tagged *M. tuberculosis rplL* | This study |
| *E. coli* BL21(DE3) pET28aTB*smpb*-$His_6$ | Contains plasmid expressing 6His-tagged *M. tuberculosis smpB* | (Ramadoss et al. 2013) |
| *E. coli* BL21(DE3) pET28a*rplI*-NTD$His_6$ | Contains plasmid expressing 6His-tagged *E. coli rplI N terminal domain* | This study |
| *E. coli* BL21(DE3) pET28a*rplI*-CTD$His_6$ | Contains plasmid expressing 6His-tagged *E. coli rplI C terminal domain* | This study |
| *E. coli* BL21(DE3) pET28a*smpb*-$His_6$ | Contains plasmid expressing 6His-tagged *E. coli smpB* | (Ramadoss et al. 2013) |
| JW2667 | Contains plasmid pCA24N expressing *E. coli alaS* | ASKA library (Kitagawa et al. 2005) |

| | | |
|---|---|---|
| JW1865 | Contains plasmid pCA24N expressing *E. coli argS* | ASKA library (Kitagawa et al. 2005) |
| JW1855 | Contains plasmid pCA24N expressing *E. coli aspS* | ASKA library (Kitagawa et al. 2005) |
| JW0913 | Contains plasmid pCA24N expressing *E. coli asnS* | ASKA library (Kitagawa et al. 2005) |
| JW0515 | Contains plasmid pCA24N expressing *E. coli cysS* | ASKA library (Kitagawa et al. 2005) |
| JW0666 | Contains plasmid pCA24N expressing *E. coli glnS* | ASKA library (Kitagawa et al. 2005) |
| JW2395 | Contains plasmid pCA24N expressing *E. coli gltX* | ASKA library (Kitagawa et al. 2005) |
| JW3531 | Contains plasmid pCA24N expressing *E. coli glyQ* | ASKA library (Kitagawa et al. 2005) |
| JW3530 | Contains plasmid pCA24N expressing *E. coli glyS* | ASKA library (Kitagawa et al. 2005) |
| JW2498 | Contains plasmid pCA24N expressing *E. coli hisS* | ASKA library (Kitagawa et al. 2005) |
| JW0024 | Contains plasmid pCA24N expressing *E. coli ileS* | ASKA library (Kitagawa et al. 2005) |
| JW0637 | Contains plasmid pCA24N expressing *E. coli leuS* | ASKA library (Kitagawa et al. 2005) |
| JW4090 | Contains plasmid pCA24N expressing *E. coli lysU* | ASKA library (Kitagawa et al. 2005) |
| JW2101 | Contains plasmid pCA24N expressing *E. coli metG* | ASKA library (Kitagawa et al. 2005) |
| JW5277 | Contains plasmid pCA24N expressing *E. coli* pheS | ASKA library (Kitagawa et al. 2005) |
| JW1703 | Contains plasmid pCA24N expressing *E. coli* pheT | ASKA library (Kitagawa et al. 2005) |
| JW0190 | Contains plasmid pCA24N expressing *E. coli proS* | (Kitagawa et al. 2005) |
| JW0876 | Contains plasmid pCA24N expressing *E. coli serS* | (Kitagawa et al. 2005) |

| | | |
|---|---|---|
| JW1709 | Contains plasmid pCA24N expressing *E. coli thrS* | (Kitagawa et al. 2005) |
| JW3347 | Contains plasmid pCA24N expressing *E. coli trpS* | (Kitagawa et al. 2005) |
| JW1629 | Contains plasmid pCA24N expressing *E. coli tyrS* | (Kitagawa et al. 2005) |
| JW4215 | Contains plasmid pCA24N expressing *E. coli* v*alS* | ASKA library (Kitagawa et al. 2005) |
| JW2502 | Contains plasmid pCA24N expressing *E. coli* n*dk* | ASKA library (Kitagawa et al. 2005) |
| JW3249 | Contains plasmid pCA24N expressing *E. coli fmt* | ASKA library (Kitagawa et al. 2005) |
| JW3301 | Contains plasmid pCA24N expressing *E. coli tufa* | ASKA library (Kitagawa et al. 2005) |
| JW3302 | Contains plasmid pCA24N expressing *E. coli fus* | ASKA library (Kitagawa et al. 2005) |
| JW0165 | Contains plasmid pCA24N expressing *E. coli tsf* | ASKA library (Kitagawa et al. 2005) |
| JW0867 | Contains plasmid pCA24N expressing *E. coli infA* | ASKA library (Kitagawa et al. 2005) |
| JW3137 | Contains plas mid pCA24N expressing *E. coli infB* | ASKA library (Kitagawa et al. 2005) |
| JW5829 | Contains plasmid pCA24N expressing *E .coli infC* | ASKA library (Kitagawa et al. 2005) |
| JW5847 | Contains plasmid pCA24N expressing *E. coli prfB* | ASKA library (Kitagawa et al. 2005) |
| JW5873 | Contains plasmid pCA24N expressing *E. coli prfC* | ASKA library (Kitagawa et al. 2005) |
| JW1202 | Contains plasmid pCA24N expressing *E. coli prfA* | ASKA library (Kitagawa et al. 2005) |
| JW0167 | Contains plasmid pCA24N expressing *E. coli frr* | ASKA library (Kitagawa et al. 2005) |

| **Plasmid** | **Description** | **Source or reference** |
|---|---|---|

| | | |
|---|---|---|
| pLC1021 | IPTG- inducible expression of *luc*-trpAt | ASKA library (Kitagawa et al. 2005) |
| pLC1063 | IPTG- inducible expression of *mCherry*-trpAt | ASKA library (Kitagawa et al. 2005) |
| prplL | IPTG- inducible expression of *E. coli rpll*<br>pCA24N-*rplL;* cat$^R$ | (Kitagawa et al. 2005) |
| pET28a | Expression vector, IPTG-inducible; kan$^R$ | Novagen |
| pET28a-TBrplI-His6 | IPTG- inducible expression of *M. tuberculosis rpll*; kan$^R$ | This study |
| pET28a-NTDrplI-His6 | IPTG- inducible expression of *E. coli rpll* N terminal domain; kan$^R$ | This study |
| pET28a-CTDrplI-His6 | IPTG- inducible expression of *E. coli rpll C*terminal domain; kan$^R$ | This study |
| pET28a-infA-His6 | IPTG- inducible expression of *M. tuberculosis infA*; kan$^R$ | (Varshney et al. 2025) |
| pET28a-infB-His6 | IPTG- inducible expression of *M. tuberculosis infB*; kan$^R$ | (Varshney et al. 2025) |
| pET28a-infC-His6 | IPTG- inducible expression of *M. tuberculosis infC*; kan$^R$ | (Varshney et al. 2025) |
| pET28a-fusA1-His6 | IPTG- inducible expression of *M. tuberculosis fusA1*; kan$^R$ | (Varshney et al. 2025) |
| pET28a-fusA2-His6 | IPTG- inducible expression of *M. tuberculosis fusA2*; kan$^R$ | (Varshney et al. 2025) |
| pET28a-tuf-His6 | IPTG- inducible expression of *M. tuberculosis tuf*; kan$^R$ | (Varshney et al. 2025) |
| pET28a-tsf-His6 | IPTG- inducible expression of *M. tuberculosis tsf*; kan$^R$ | (Varshney et al. 2025) |
| pET28a-prfA-His6 | IPTG- inducible expression of *M. tuberculosis prfA*; kan$^R$ | (Varshney et al. 2025) |
| pET28a-prfB-His6 | IPTG- inducible expression of *M. tuberculosis prfB*; kan$^R$ | (Varshney et al. 2025) |
| pET28a-frr-His6 | IPTG- inducible expression of *M. tuberculosis frr*; kan$^R$ | (Varshney et al. 2025) |
| pET28a-SmpB-His6 | IPTG- inducible expression of *M. tuberculosis smpB*; kan$^R$ | (Varshney et al. 2025) |
| pGEM-ssr | Expresses *M. tuberculosis ssr* gene off a T7 promoter; | This study |
| pDHFR | Expresses DHFR off a T7 promoter; Ap$^R$ | New England Biolabs |

| **Oligonucleotide** | **Sequence 5' to 3'** | **Source or reference** |
|---|---|---|
| TB_L12_F | ACTTTAAGGAGATATACATGGCAAAGCTCTCCACC | This study |

| | | |
|---|---|---|
| TB_L12_R | AGTGGTGGTGGTGGTGGTGCCACTTGACGGTGACGGTG | This study |
| L12_NTD_F | ACTTTAAGAAGGAGATATACCATGTCTATCACTAAAGATCAAATCATTGAAG | This study |
| L12_NTD_R | AGTGGTGGTGGTGGTGGTGCCACCCGGAAACACCGAATTTTTC | This study |
| L12_CTD_F | AGTGGTGGTGGTGGTGGTGCTCGAGTTTAACTTCAACTTCAGCGC | This study |
| L12_CTD_R | ACTTTAAGAAGGAGATATACCATGGCTGCTGAAAAAACTGAATTC | This study |
| TBinfAfor | ACTTTAAGAAGGAGATATACATGGCCAAGAAGGACGGTG | (Varshney et al. 2025) |
| TBinfArev | AGTGGTGGTGGTGGTGGTGCTTGTACCGGTACACGATGC | (Varshney et al. 2025) |
| TBinfBfor | ATATCCATGGTGGCAGCAGGTAAGGCC | (Varshney et al. 2025) |
| TBinfBrev | ATACTCGAGCGCGCGTTCCTTCTGGACC | (Varshney et al. 2025) |
| TBinfCfor | ACTTTAAGAAGGAGATATACATGATCAGCACTGAAACCCG | (Varshney et al. 2025) |
| TBinfCrev | AGTGGTGGTGGTGGTGGTGCCCGTTCGGTGCGGCTTTG | (Varshney et al. 2025) |
| TBfusA1for | ACTTTAAGAAGGAGATATACATGGTGGCACAGAAGGACGTG | (Varshney et al. 2025) |
| TBfusA1rev | AGTGGTGGTGGTGGTGGTGCCCCTCGCCCGTCGCCTTC | (Varshney et al. 2025) |
| TBfusA2for | ACTTTAAGAAGGAGATATACATGGCCGACAGAGTGAATGCTTCCC | (Varshney et al. 2025) |
| TBfusA2rev | AGTGGTGGTGGTGGTGGTGCCCGCCGGCCCCGGCCTTT | (Varshney et al. 2025) |
| TBtsffor | ACTTTAAGAAGGAGATATACATGGCGAACTTCACTGCCG | (Varshney et al. 2025) |
| TBtsfrev | AGTGGTGGTGGTGGTGGTGCCCAGCCTGGCCCACCTCG | (Varshney et al. 2025) |
| TBtuffor | ACTTTAAGAAGGAGATATACATGGTGGCGAAGGCGAAG | (Varshney et al. 2025) |
| TBtufrev | AGTGGTGGTGGTGGTGGTGCCCCTTGATGATCTTGGTGAC | (Varshney et al. 2025) |
| TBprfAfor | ACTTTAAGAAGGAGATATACATGACGCAGCCAGTGCAG | (Varshney et al. 2025) |
| TBprfArev | AGTGGTGGTGGTGGTGGTGCCCTGATTGTCGCAACCGG | (Varshney et al. 2025) |
| TBprfBfor | ACTTTAAGAAGGAGATATACATGCCGGTAACCTTGGCTG | (Varshney et al. 2025) |
| TBprfBrev | AGTGGTGGTGGTGGTGGTGCCCGTCGTCATTACGTCGATTG | (Varshney et al. 2025) |
| TBfrrfor | ACTTTAAGAAGGAGATATACATGATTGATGAGGCTCTCTTCG | (Varshney et al. 2025) |

| | | |
|---|---|---|
| TBfrrrev | AGTGGTGGTGGTGGTGGTGCCCGA CCTCCAGCAGCTCG | (Varshney et al. 2025) |
| TB_SmpB_F | CGCGGCAGCCATATGGTGTCCAAG TCGTCGCGTG | (Dillon et al. 2017) |
| TB_SmpB_R | CTCGAGTGCGGCCGCAAGCTTCAG GTCATGCCCTTAGCGC | (Dillon et al. 2017) |
| MtbSsrAF | GAAATTAATACGACTCACTATAGGG GCTGAACGGTTTCGACTT | (Dillon et al. 2017) |
| MtbSsrAR | TGGTGGAGCTGCCGGGAATC | (Dillon et al. 2017) |
| dhfr-ns reverse | AAACCCCTCCGTTTAGAGAGGGGT TTTGCTAGTATCCGCCGCTCCAGAA TCTCAA AGCAA | (Ramadoss et al. 2013) |
| dhfr-stop reverse | TTAACCCCTCCGTTTAGAGAGGGGT TTTGCTAGTATCGCGATGGCTTCAT CCACCGACTT | (Ramadoss et al. 2013) |

**Note S1: Synthesis & Characterization of 4-azido-N-(3-(4-ethynylbenzamido)-1-phenyl-1H-1,2,4-triazol- 5-yl) benzamide (KKL-2108)**

#### GENERAL

All chemical reagents and ACS grade organic solvents were purchased from Sigma-Aldrich (St Louis, MO) unless otherwise noted. 1-Phenyl-1H-1,2,4-triazole-3,5-diamine was purchased from Life Chemicals (Kyiv, Ukraine). Flash chromatography was performed on 60-200 mm high-purity 60Å silica gel (BDH). Thin layer chromatography was performed on glass silica gel plates (Fluka) with fluorescentindicator. NMR analyses were performed on a Varian AV-360 or a DRX-400 Nuclear Magnetic Resonance (NMR) Spectrometer (Bruker). Mass spectroscopy analysis for compound confirmation was performed using an ABI 3200 QTRAP LC/MS/MS system (Applied Biosystems).

**SCHEME S1**

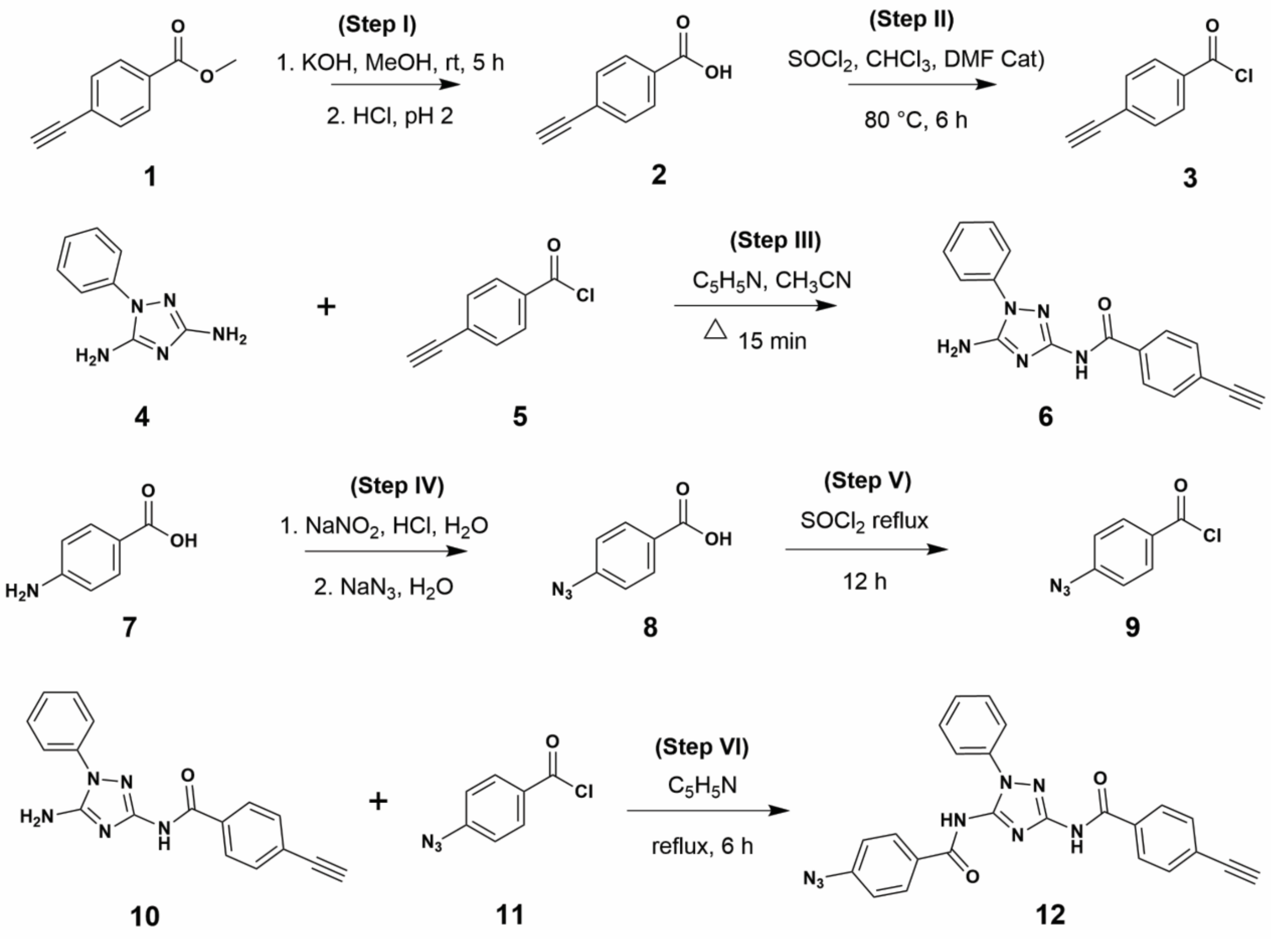


**4-ethynylbenzoic acid (2)**. Methyl-4-ethynylbenzoate (800 mg, 5.0 mmol) was added to flaskcontaining a 12 ml solution of THF: $CH_3OH$ (1:1, v/v) and stirred to dissolve. To this mixture was added 5 ml of 1 M aqueous NaOH slowly while stirring. The reaction was left to proceed at room temperature for 6 h. The volatile solvents were evaporated under reduced pressure. 30 mL water was added to the residual solid, and the resulting suspension was filtered to remove any solid material. The filtrate was transferred to a beaker and acidified with 1 M HCl to pH ~ 1. The brownish solid that precipitated was isolated by vacuum filtration, washed three times with 40 mL water, dried under vacuum and dried in a desiccator overnight. Analysis of this solid yielded the desired compound (**1**) as a fine powder in quantitative yield. $^{1}$H-NMR (400 MHz, DMSO-$d_6$): $\delta$ 4.44 (s,1H); 7.59 (d, $J$ = 8.2 Hz, 2H); 7.95 (d, $J$ = 8.2 Hz, 2H); 12.9 (s, 1H). $^{13}$C-NMR (100 MHz, DMSO-$d_6$): $\delta$ 83.9, 84.3, 126.8, 130.3, 132.9, 168.0.

**4-ethynylbenzoyl chloride (3).** 4-ethynylbenzoic acid (380 mg, 2.6 mmol) was added to a flask containing 5 mL chloroform. Thionyl chloride (0.95 mL, 13 mmol) was added followed by 2 drops of dimethylformamide. The flask was fitted with a condenser and transferred to an oil bath at 75 °C. This reaction mixture was refluxed for 12 h and the volatile solvents were removed by vacuum distillation at 80 °C. The dark brown crude residue was used in the amidation reaction without further purification.

**N-(5-amino-1-phenyl-1H-1,2,4-triazol-3-yl)-4-azidobenzamide (5).** Compound **4** (400 mg, 2.3 mmol) was added to a round-bottomed flask containing 5 mL dry acetonitrile and 0.5 mL pyridine. A solution of 4-ethynylbenzoyl chloride (416 mg, 2.5 mmol) in 3 mL dry acetonitrile was added dropwise while stirring the mixture. The flask was fitted with a condenser and the reaction was refluxed at 95 °C for 15 min and cooled -20 °C. The precipitate that formed was isolated by vacuum filtration and dissolved in methanol. TLC analysis showed >60% conversion. The crude product was purified by flash chromatography using 5% methanol in dichloromethane to obtain white crystals, 233 mg (33.4 % recovery). $^{1}$H-NMR (400 MHz, DMSO-$d_6$): $\delta$ 3.17 (s, 1H); 6.55 (s, 2H); 7.36 (t, $J$ = 8.4 Hz, 1H); 7.45 – 7.62 (m, 6H); 7.94 (d, $J$ = 8.4 Hz, 2H); 10.6 (s, 1H). $^{13}$C-NMR (100 MHz, DMSO-$d_6$): $\delta$ 69.1, 82.9, 122.5, 125.0, 126.8, 128.2, 129.4, 131.7, 134.0, 137.2, 154.0, 164.4.

**4-azidobenzoic acid (7).** 4-aminobenzoic acid (2 g, 14.6 mmol) was added to a 250 mL round-bottom flask equipped with a stir-bar on an ice bath maintained at 0 – 5 °C. 2N HCl (100 ml) was added and the mixture was stirred to dissolve the solid. A solution of sodium nitrite (2.5 g, 36.5 mmol) dissolved in water was added dropwise with vigorous stirring over 15 min. The mixture was stirred for an additional 15 min until it was completely clear of any precipitated material. A solution of sodium azide (4.8 g, 73.8 mmol) dissolved in water was slowly added to the reaction over 30 min, resulting in a white precipitate. The reaction mixture was left to stir for an additional 30 min at 0 °C. The solid was isolated by vacuum filtration, washed with cold and dried under vacuum to afford the desired product in quantitative yield. $^1$H-NMR (400 MHz, $CDCl_3$): $\delta$ 7.61 (d, $J$ = 8.2 Hz, 2H); 7.99 (d, $J$ = 8.2 Hz, 2H); 12.3 (s, 1H). $^{13}$C-NMR (100 MHz, $CDCl_3$): $\delta$ 119.9, 123.1, 129.7, 136.9, 167.5.

**4-azidobenzoyl chloride (8).** Thionyl chloride (5 mL, 27.6 mmol) was added to a flask containing 4-azido benzoic acid (250 mg, 1.5 mmol). This flask was fitted with a condenser and transferred to a pre-heated oil bath maintained at 90 – 100 °C. The reaction mixture was stirred overnight and excess $SOCl_2$ was removed by vacuum distillation at 90 °C. The resulting crude residue was used in the coupling step without further purification due to the sensitivity of the product to moisture.

**4-azido-N-(3-(4-ethynylbenzamido)-1-phenyl-1H-1,2,4-triazol-5-yl)benzamide (9, KKL-2108).** Compound **5** (230 mg, 0.76 mmol) was added to 5 mL pyridine in a dry round-bottomed flask and stirred to dissolve. A solution of 4-azidobenzoyl chloride (152 mg, 0.84 mmol) in 1 mL pyridine was added to the reaction flask while stirring. The flask was fitted with a condenser and the reaction was refluxed at 100 °C for 12 h. The reaction mixture was cooled to room temperature and poured into a beaker with crushed ice. The precipitate was isolated by vacuum filtration, washed with water, and dried in a desiccator. Purification of the crude product was performed by flash chromatography over silica gel using 5% methanol in dichloromethane to obtain 123 mg of the desired product (36% yield). $^{1}$H-NMR (400 MHz, DMSO-$d_6$): $\delta$ 3.18 (s, 1H); 7.37 (t, $J$ = 8.4 Hz, 1H); 7.40 (t, $J$ = 8.4 Hz, 2H); 7.43 – 7.57 (m, 6H); 7.81 (d, $J$ = 8.4 Hz, 2H); 7.96 (d, $J$ = 8.4 Hz, 2H); 10.8 (bs, 2H). $^{13}$C-NMR (100 MHz, DMSO-$d_6$): $\delta$ 68.8, 82.2, 121.9, 124.8, 126.5, 127.9, 129.0, 129.5, 130.9, 133.7, 136.9, 152.9, 153.2, 163.8, 164.0. TOF MS ES (+): observed 449.1494, calculated 449.1475.

**MS Analysis Report for KK-2108**

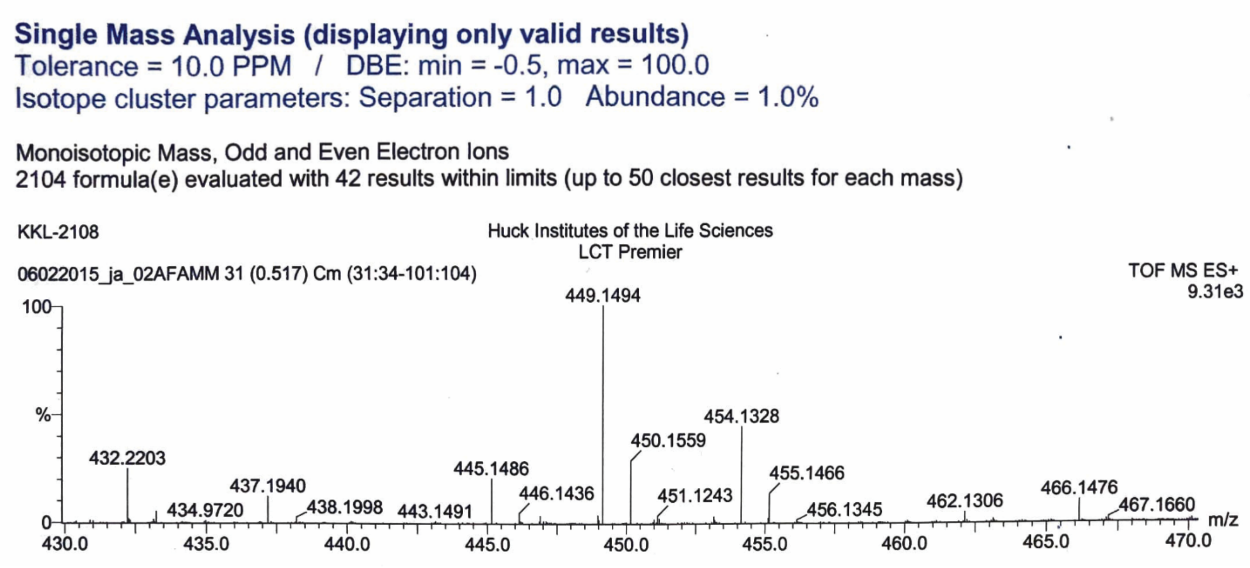
